# From comparative gene content and gene order to ancestral contigs, chromosomes and karyotypes

**DOI:** 10.1101/2022.09.28.509880

**Authors:** Qiaoji Xu, Lingling Jin, Chunfang Zheng, Xiaomeng Zhang, James Leebens-Mack, David Sankoff

## Abstract

To reconstruct the ancestral genome of a set of phylogenetically related descendant species, we use the Raccroche pipeline for organizing a large number of generalized gene adjacencies into contigs and then into chromosomes. Separate reconstructions are carried out for each ancestral node of the phylogenetic tree for focal taxa. The ancestral reconstructions are monoploids; they each contain at most one member of each gene family constructed from descendants, ordered along the chromosomes. We design and implement a new computational technique for solving the problem of estimating the ancestral monoploid number of chromosomes *x*. This involves a “***g***-mer” analysis to resolve a bias due long contigs, and gap statistics to estimate ***x***. We find that the monoploid number of all the rosid and asterid orders is ***x* = 9**. We show that this is not an artifact of our method by deriving ***x* ≈ 20** for the metazoan ancestor.

---

Evolutionary inference on a set of species in a biological family, order or higher grouping, implies the reconstruction of ancestral phenotypes or genotypes. Phenotypic reconstruction, essentially genome-free, can be derived from comparative macroscopic or microscopic evidence from extant forms or fossils, while inference of genotypes is based on the genome sequences. In this work, we focus on analyses of annotated genes and the chromosomal ordering of these genes in the genomes of extant organisms.

The **genome-free** inference of the basic (or monoploid) ancestral chromosome number *x*, based on the values of *x* for a very large number of extant species, has a long history in plant evolutionary biology, exemplified in Grant’s ground-breaking 1963 work [1, pp. 483-487]. More recently, sophisticated combinatorial optimization techniques and Bayesian inference approaches have been developed to infer ancestral chromosome numbers [2– 4], but these approaches do not aim to elucidate the genetic composition nor the chromosomal structure of the ancestral species. In contrast, the present **genome-based** study, based on all the common gene adjacencies (including “gapped” adjacencies) among species within a phylogenetic context seeks to recover the largest possible consistent subset of these adjacencies, organized into hypothetical ancestral chromosomes of a monoploid ancestor. One such set of chromosomes is constructed for each ancestral node of the phylogenetic tree describing the relationship among the analyzed species.

Inference about ancestral genome structure is difficult in the plant kingdom (as reviewed in [5]). Adjacencies are disrupted in plant genomes by whole genome duplication followed by random deletion of duplicate genes (“fractionation”), in addition to niche-specific expansion and contraction of gene families, chromosomal rearrangements, fissions and fusions and by rampant invasions and culling of transposons, which typically comprise the majority of the genome, and other processes. Much of the work on reconstruction, e.g. [5,6], relies on a bottom-up, greedy stepwise inference of “contiguous ancestral regions”, incorporating external information such as on whole genome duplication events, without particular attention to the number and nature of individual chromosomes.

The focus on monoploidy in our method, however, permits a single-step reconstruction of ancestral chromosomal fragments, **contigs**, without any recourse to information external to the given set of phylogenetically-related genomes. The maximum weight matching (mwm) algorithm embedded in the Raccroche pipeline [7,8] assures a robust monoploid reconstruction; each ancestor contains at most one representative of each gene family, organized into a number *x* of ancestral chromosomes, the “basic number”, and ordered along the chromosomes in a way most consistent with the gene order in their extant descendants. Somewhat unexpectedly, as illustrated in Figure 1, the method produces more clear-cut inferences on clades of more remotely related species, such as six genomes each from a different monocot order [8], or six eudicot species from different orders [9], than sampling of lineages within orders or families, where the results are degraded by high levels of noise [10].

**Fig. 1.**
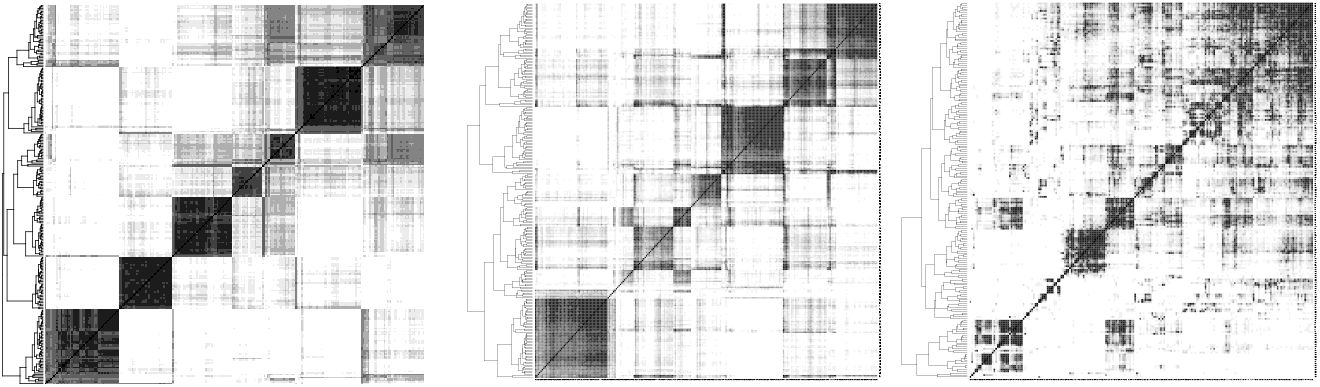
Clear-cut results of Raccroche across monocot (left) and eudicot (center) orders, compared to noisy results for intra-ordinal analysis of Fagales (right). Heat maps compare an optimal clustering of ancestral chromosomal fragments with itself, with dark cells representing two fragments which co-occur on the same chromosome in several extant genomes.

In this paper, we trace the origin of this noise to a severe bias that arises when analyzing a clade of closely related genomes. We devise a new method to eliminate this bias and thus mitigate the resulting noise. In addition we introduce new methods to determine the statistically optimal number of chromosomes and reconstruct these chromosomes automatically.

We use our pipeline to determine the monoploid number of the common ancestor of all sampled species for each order analyzed as well as ancestors represented by internal nodes for each phylogeny. For each of six rosid orders Fagales, Cucurbitales, Malpighiales, Myrtales, Malvales and Sapindales, and five asterid orders - Asterales, Gentianales, Lamiales, Solanales and Ericales, species were chosen based largely on the availability of annotated chromosome-level genome sequences representing many or most of the families in each order.

A result of our genome-based method is that the monoploid number of the rosid and asterid orders is determined to be *x* = 9, compared to the *x* = 7 or *x* = 8 estimated from a recent genome-free study [4].

## Methods

### Generating sets of long contigs

To infer gene content and gene order for each chromosome in each ancestral genome in a phylogeny, we identify a large number of generalized [11] (or “gapped” [12]) gene adjacencies, allowing for example, up to 7 spacer genes between the two considered adjacent, from all chromosomes in the set of input genomes and then infer adjacencies for each ancestral node in the species phylogeny. To do this, for each ancestor, graphs generated with all phylogenetically informative generalized adjacencies as vertices and edges joining any two adjacencies that each contain one of the 5’ and 3’ ends of the same gene, are analyzed using the mwm algorithm. This outputs inferred linear ancestral “contigs”, each containing up to several hundred genes.

The data used for this work are annotated, chromosome-level or other high-quality genome sequences, accessible on the CoGe [13, 14] platform, or uploaded to a dedicated repertoire on this platform from public sources, as well as phylogenies for each of the orders studied, as extracted from recent literature and databases [15, 16]. The only pre-processing software required was the SynMap [13, 14] comparative genomics package, also on the CoGe platform, which produces syntenically validated homology identification between genomes (orthologs) and within single genomes (paralogs). The term “gene” here is used broadly to refer to gene families, or sets of homologous genes in the extant genomes as well as the hypothetical ancestral genes inferred by our procedures.

An important observation is that the lengths of the contigs constructed from the mwm output are highly variable, ranging from a single adjacency to several hundred in some cases. The lengths of the longest few contigs provide a measure of the conservation of gene order among the extant genomes, up to several hundred when reconstructing the ancestor of a plant family or order, compared to less than a hundred for analyses of more distantly related species in a more inclusive taxon such as the monocot or eudicot clades (Figure 1a,b). On the other hand, the extreme variability of contig lengths encountered at the family or order level can lead to ambiguous or distorted clustering of contigs into inferred ancestral chromosomes.

### Clustering the chromosomal co-occurrence matrix and the long-contig bias

To group contigs into clusters reflecting ancient chromosomes, we match each contig against the chromosomes of the extent genomes, and count the number of times any two contigs match the same chromosome, taking account of their ordering, possibly twice or more within a single genome. The resulting co-occurrence matrix, smoothed by a correlation analysis of pairs of contigs [8], is then submitted to a complete-link clustering analysis to distinguish the contigs, and hence the gene content, appropriate to each hypothetical ancestral chromosome. Once contig content of each chromosome is posited, the data on relative order of each pair of contigs on a chromosome is submitted to a Linear template Ordering Problem routine to locate them along the chromosome. It is at this point that the variability of contig length leads to serious biases, due to the longest contigs tending to group together to produce unrealistically large clusters, as illustrated in Figures 2 and 3 for a 7-chromosome analysis of a Fagales ancestor. The reason for this lies in the gene families with more than one (but ≤ 10) representatives in some extant genomes. Our focus on small gene families (larger families are excluded from the analysis) rather than inferred orthologs for our ancestral contig reconstructions avoids error in orthology assignment, such as those due to widespread whole genome duplication events in plant lineages, while at the same time increasing overlap in “gene” adjacencies among analyzed genomes. This inclusion, however, allows the mwm to join distantly homologous or non-homologous generalized adjacencies when assembling the ancestral contigs. This may result in splicing of two, three or more part-contigs from different chromosomes. Thus the various long contigs of that result tend to involve many genes in common, deriving from several chromosomes in the extant genomes. For shorter contigs, this can also happen, but is rare.

**Fig. 2.**
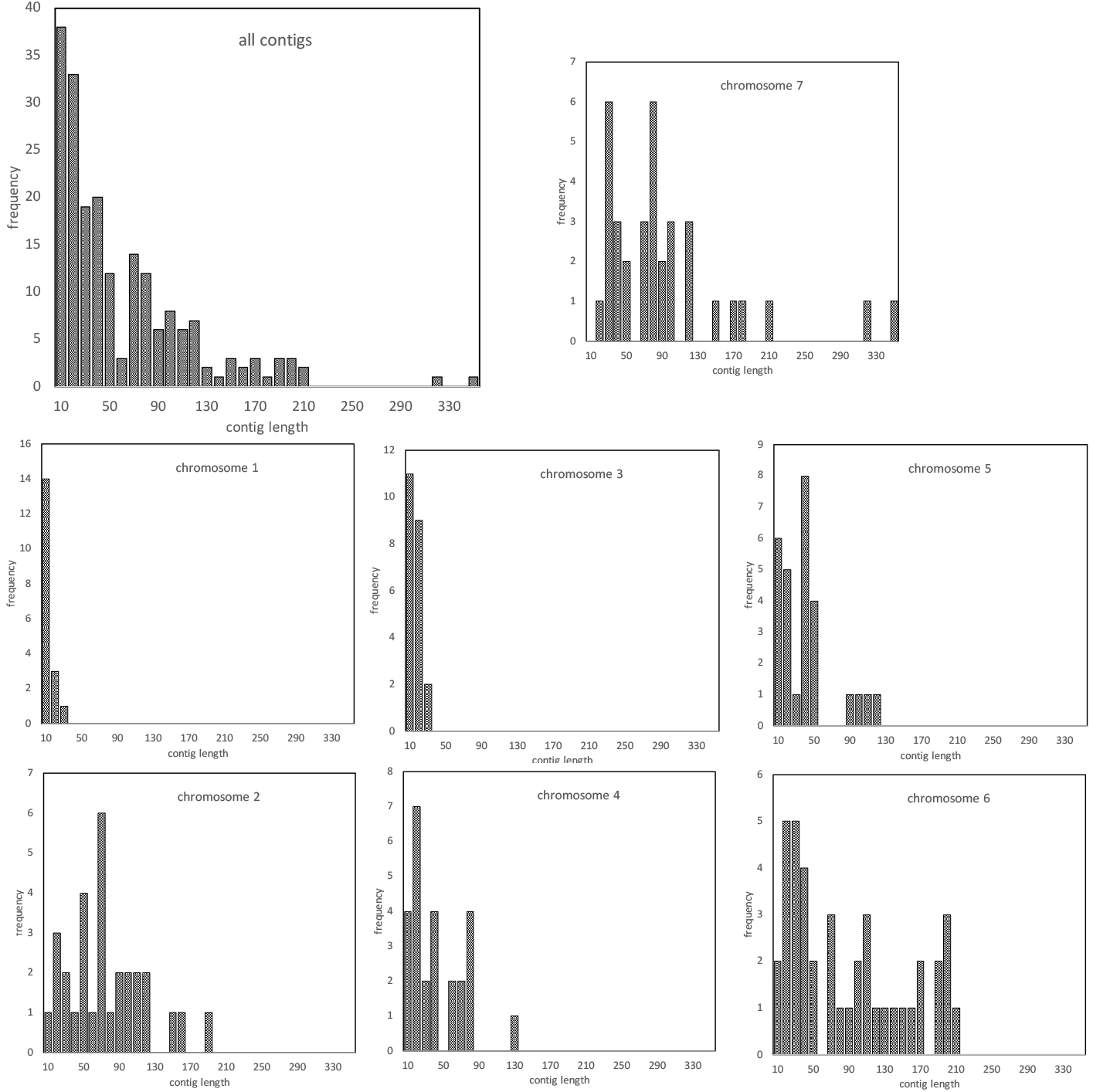
Contigs per chromosome, Fagales Ancestor 2 (which we use as an example throughout; see Figure 8). N.B. Frequency scale differs among chromosomes.

**Fig. 3.**
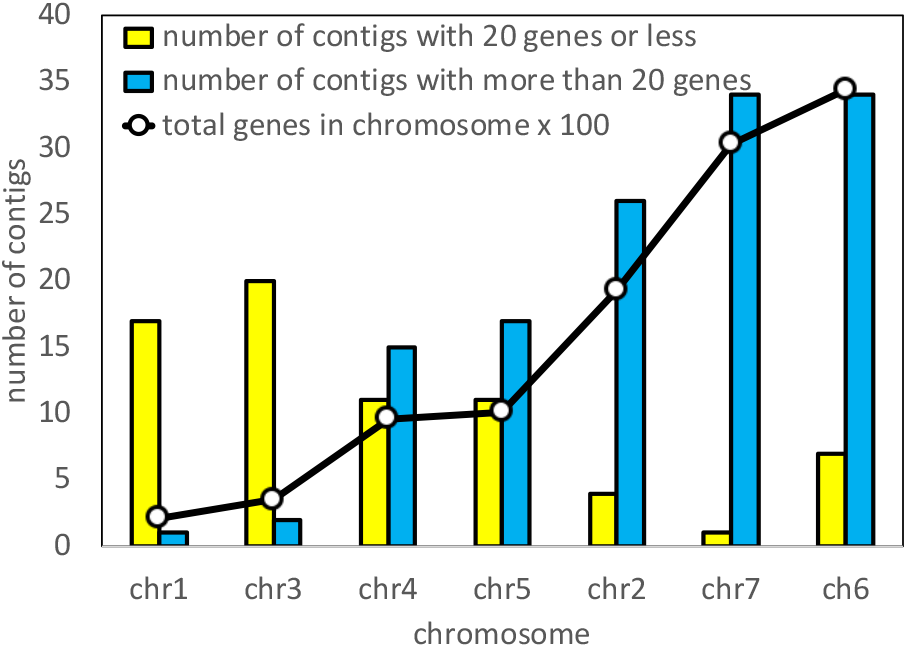
Statistics on the chromosomes in Figure 2, showing bias in the assignment of long contigs.

The effect of this artifact is apparent not only in imbalances among the inferred chromosomes, as in Figures 2 and 3, but also very noisy heat maps as in Figure 1c.

### Introducing g-mers to remove bias and noise

As illustrated with the case of Fagales in Figures 2 and 3, the presence of extreme-length contigs produces biases in counting contig co-occurrences, leading to an unbalanced set of chromosomes. This would seem a severe problem with the Raccroche method, especially when applied to sets of closely related genomes. It is possible, however, to completely remove the length bias by simply cutting each contig of length *L* into approximately *L/g* contigs of length *g*, called *g*-mers, exempting of course for contigs where *L* ≤ *g* already. We can then carry out the cluster analysis based on the *g*-mers derived from all the contigs.

A clustering may be visualized by constructing a “heat map” comparing the cluster to itself, as in Figure 4, which shows the improvement in distinctness and size balance of an ancestral genome reconstruction through the use of *g*mers. The figure suggests that at least in this example, any choice of *g* results in a clear improvement.

**Fig. 4.**
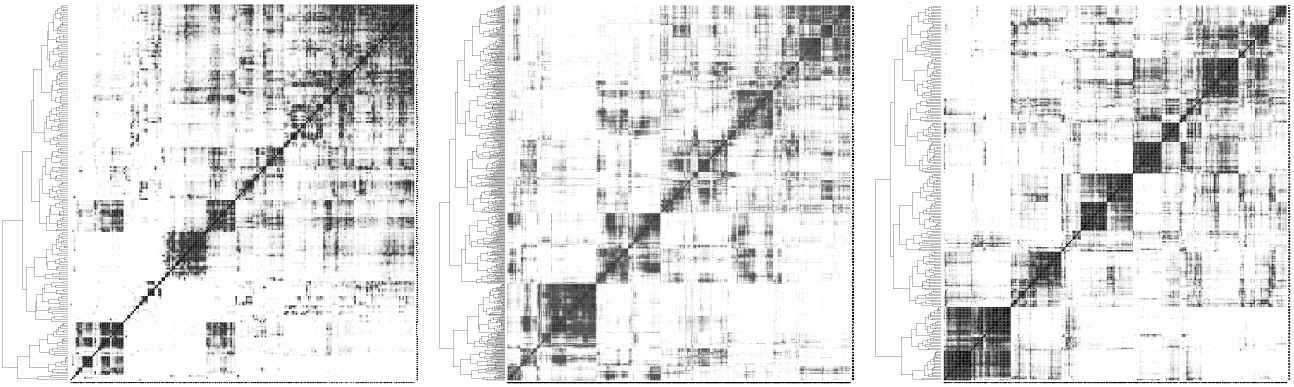
Heat map of Fagales Ancestor 2 based on full contigs (left) and on 20-mers (center) and 40-mers (right). The hierarchical cluster for each heat map depicted at the left-hand side. Shades of grey in cells, representing frequency of chromosomal contig co-occurrence in extant genomes, controlled to have equal darkness proportions across all heat maps.

### Sampling of Maximum Weight Matching Solutions

The mwm algorithm that we invoked to find an optimal matching of the adjacencies does not return a unique solution. Indeed, given the massive amount of generalized adjacencies in our analyses, there may be many thousands of equally optimal solutions, usually quite similar 95% *±* 3% but exhibiting considerable amount of variation in gene content and gene order among the ancestral contigs.

The mwm algorithm constructs all these optimal sets of matchings without taking into account the properties of the contigs they determine or the clustering used to build chromosomes. In particular, whether they give rise to neat clusters in the complete-link analysis or not, does not influence the mwm constructions.

For this reason, we sample a number (100 or 50 in this study) of optimal mwm solutions. For each *g*, then, our problem becomes one of searching among these solutions for one that gives the clearest clustering pattern, towards which end we implement the following definition and analysis.

### Gap statistics to determine the basic chromosome number x

To determine the number of chromosomes in an ancestral genome, we cut the hierarchical clustering at a series of levels, starting near the root and proceeding towards the leaves, at each step increasing the number of clusters *k* by 1.

The gap statistic method [17] tests the significance of the *k*-cluster analysis for *k* = 2, 3, *· · ·* against a null hypothesis that there is no clustering, i.e., *k* = 1. A plot of this gap statistic, as on the left in Figure 5, for a *k*-chromosome analysis shows a rapid, though concave, rise for *k* = 2, 3, *· · ·*, representing real improvements in the explanatory power of larger *k*, until a point where the rate of the increase drops visibly, becoming a slow linear trend measuring non-explanatory overfitting by excessive chromosome numbers. The point where one trend gives way to the other may be taken as an estimate of the basic chromosome number *x*. Since this typically varies among the mwm samples, we plot all the values, plus their mean (indicated by back dots in the display), on a single graph as a first step in the search for the best value of *x*. There are various methods for detecting the inflection point of a curve transitioning from one trend to another. The intersection of linear fits to the gap statistic for first few values of *k*, and for the last few *k* [10], proves to be unstable and misleading estimate, largely due to the variable concavity of the first trend. The method known as “kneedles” [18], based on finding the point of maximum curvature in the plot, is biased towards low values of *k*, also because of the template concavity of the initial trend. And there are many other methods, but the most appropriate approach for our data is to fit least squares line to the noisy trend, based on *k* = 12, …, 20 and to simply take *x* to be largest *k* for which the gap statistic exceeds the prediction of this line.

**Fig. 5.**
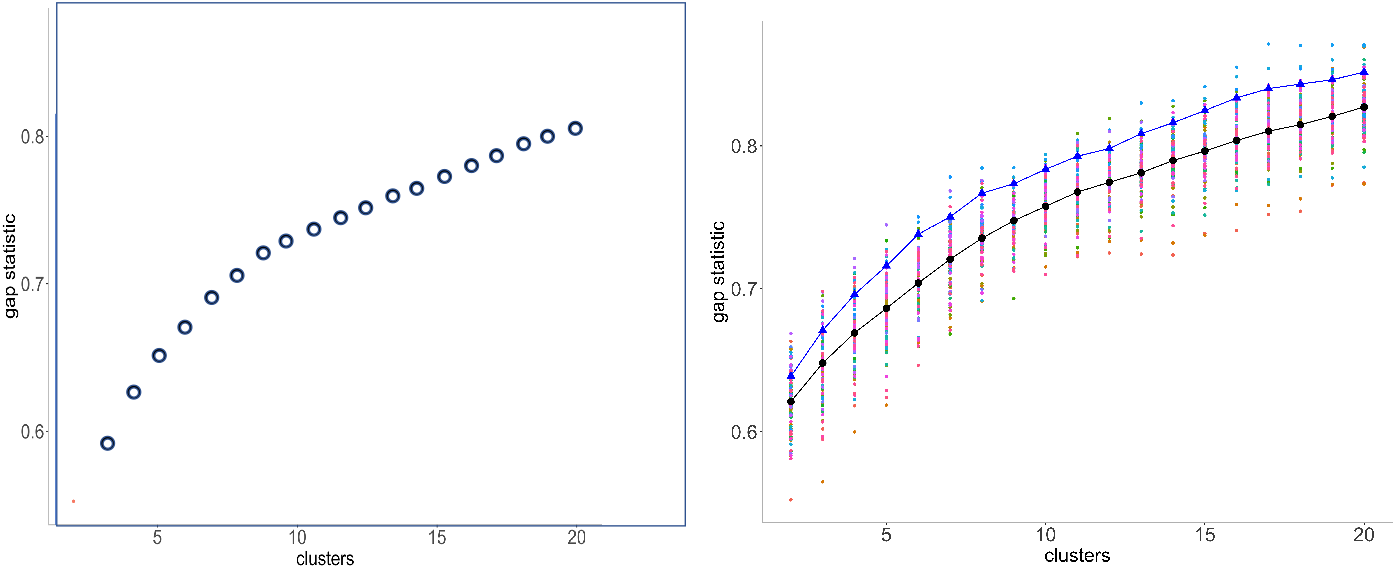
Left: Typical gap statistic for a single mwm sample. Right: Gap statistics for 100 samples with Fagales Ancestor 2. Black: means. Blue: means of 10 “best (as described in text).

As stressed above, since the mwm samples vary as to how clear a clustering they produce, we are not directly interested in the mean gap statistics, but seek instead the samples with the highest values of this statistic. Thus for each value of *k*, we note the 10 best values out of the hundred samples, and retain those samples that appear in the 10 best at least twice for 4 ≤ *k* ≤ 10 In the applications to be described below, this generally resulted in a choice of 9-15 samples. (For the metazoan example introduced later, we surveyed *k* for 4 ≤ *k* ≤ 20 to find the best mwm runs.) The mean gap statistics for these samples appear as blue dots in Figure 5.

We consider the inflection point of the gap statistics in terms of the blue dots as the most pertinent to the estimate of *x*. And we consider the clustering with the highest score as the best choice to represent the ancestral monoploid. This clustering can be slightly adjusted, without changing *k*, using the Dynamic Cutting routine [19], which corrects for deeply nested subclustering as well as outlier contigs. Heat maps for the reconstructions in this paper are available in Supplementary material A.

### Choice of g for g-mers

For each ancestor, we first break down the contigs into *g*-mers as described above, calculate a new co-occurrence matrix, construct a hierarchical cluster and carry out the gap statistic analysis for each of several values of *g*. This involves choosing the best mwmsample and cluster number *k*. To see the effect of choice of *g* on the properties of the reconstructed ancestor, we can use the following statistics.

#### Coherence

The coherence of the construction is reflected in the resemblance between each chromosome, in either an extant or ancestral descendant genome, and some chromosome of its immediate ancestor. Then for each chromosome we calculate the maximum proportion of its genes originating in any chromosome of its immediate ancestor. We average this over all chromosomes of the descendant genome. And take an overall average over all descendants in the phylogeny, separately for extants and ancestors.

#### Coverage

This is simply the number of genes in the reconstructed genome.

#### Choppiness

When painting an extant genome by the colors of the chromosomes of the nearest ancestor, as illustrated in Figure 12, we define the choppiness by counting the number of single colour regions (*>* 300 Kb) on all the extant chromosomes. This an indicator of how much genome rearrangement has intervened between the ancestor and its descendent.

Figure 6 suggests that for *g >* 10, there is little change in the quality of the reconstruction for *g* up to 40, at least. While the levels of the evaluation statistics vary from order to order, as seen in the Supplementary materials B, there is no systematic dependence on *g*.

**Fig. 6.**
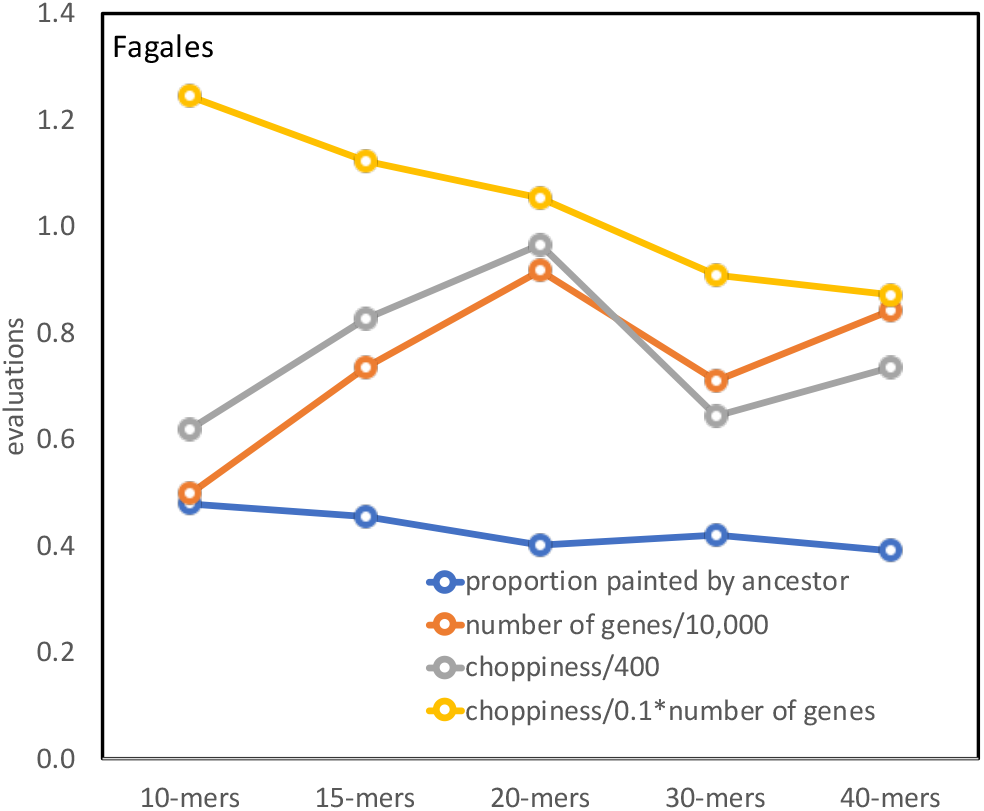
Reconstruction quality as a function of *g*. Averaged over all extant Fagales genomes.

### Alternative clustering

Our final partition of the contigs into discrete clusters does not retain any interchromosomal relationships. However, the stepwise decomposition of the higher order links in the hierarchical clustering, as a “greedy” procedure, may lead to suboptimal results. This is mitigated by our use of dynamic tree cutting, which can redress inappropriate hierarchical constraints, and even more important, by the extensive sampling of mwm solutions, from which the most cleanly separated reconstructions are selected.

Approaches such as *k*-means are also possible, but this incurs stability issues with the co-occurrence matrix and does not possess the contig ordering properties of the hierarchical clustering.

Perhaps more important is that the matrix of contig co-occurrences, as well as the derived correlations between contigs, are situated in a very high dimensional space. With such data, much of the variance resides in relatively few dimensions. Principal Component Analysis (PCA) allows us to home in on these important dimensions, relegating the rest to noise. The clustering can then be done based on the coordinates of the contigs in these few dimensions only. Using the HCPC package for R, we produce the two-dimensional display for a Fagales Ancestor 1 in Figure 7. Similar plots for the other ten orders are presented in Supplementary materials E.

**Fig. 7.**
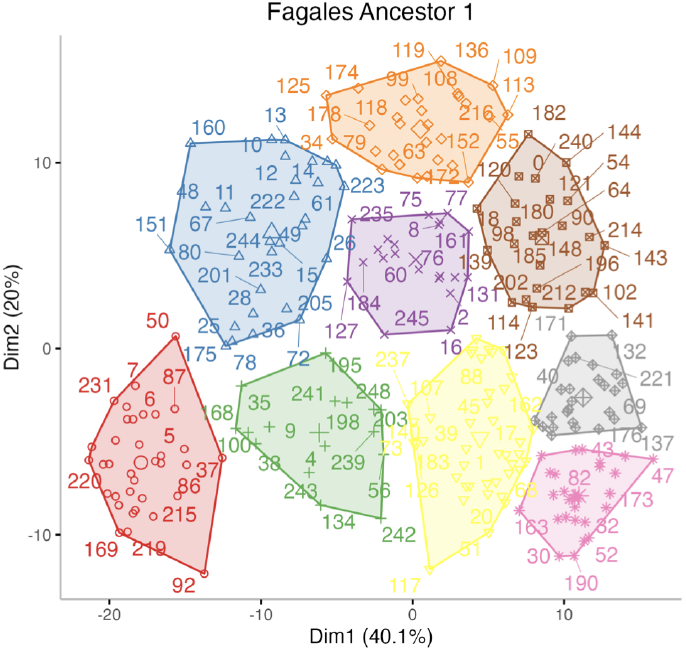
Nine clusters in first two principal components for Fagales Ancestor 1 20-mer data

**Fig. 8.**
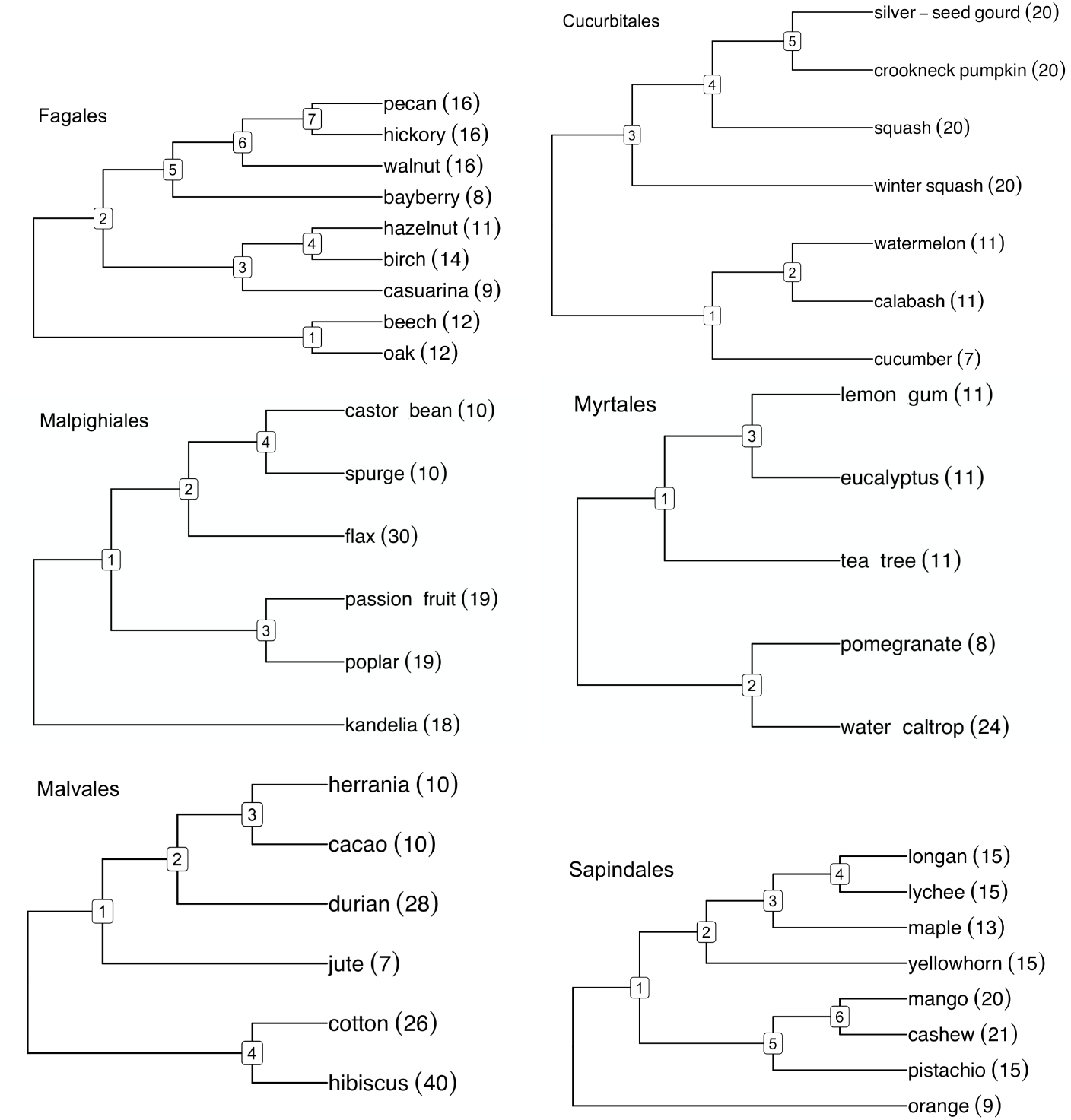
Phylogenies of rosid orders with haploid numbers of chromosomes.

**Fig. 9.**
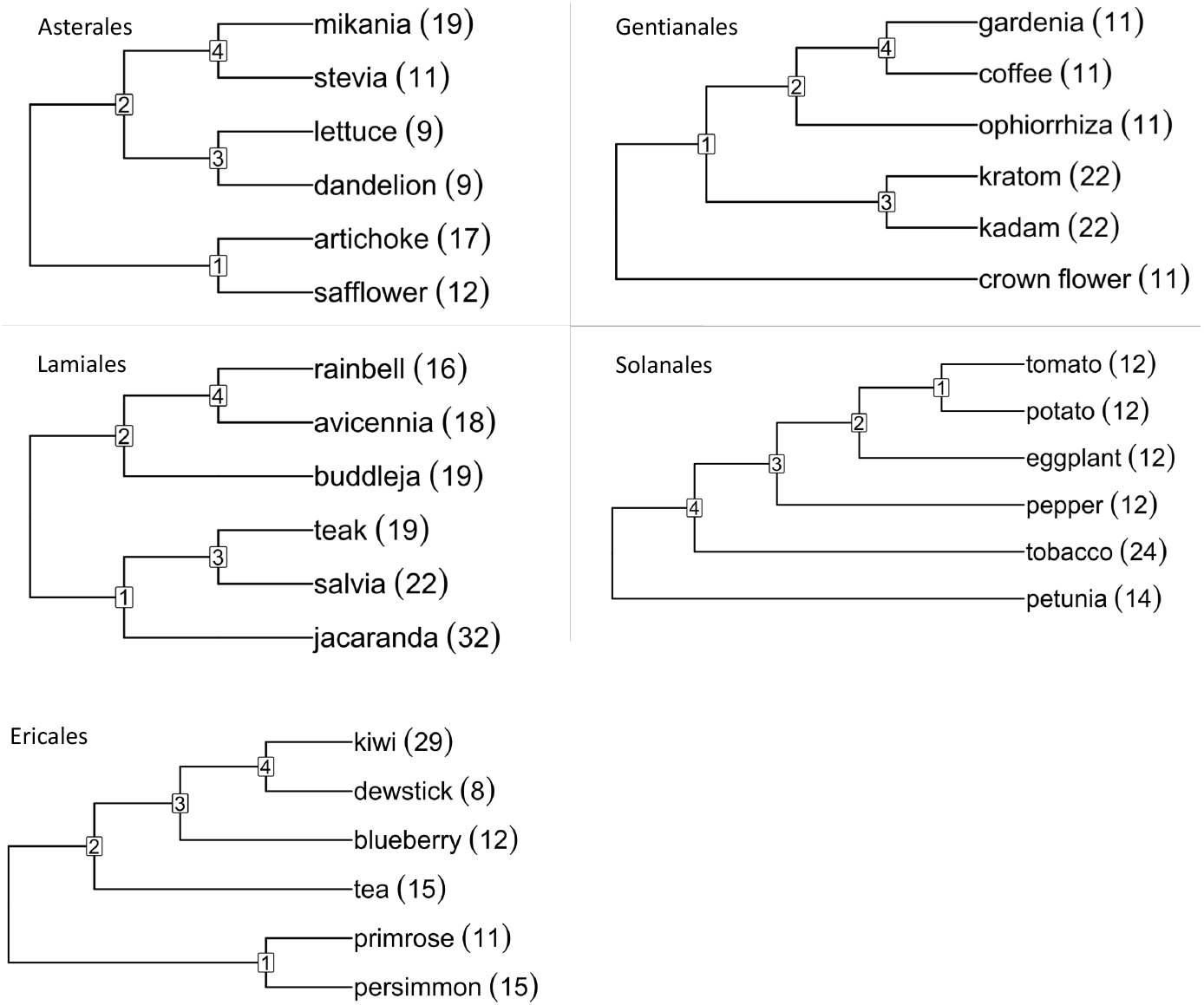
Phylogenies of asterid orders with haploid numbers of chromosomes.

**Fig. 10.**
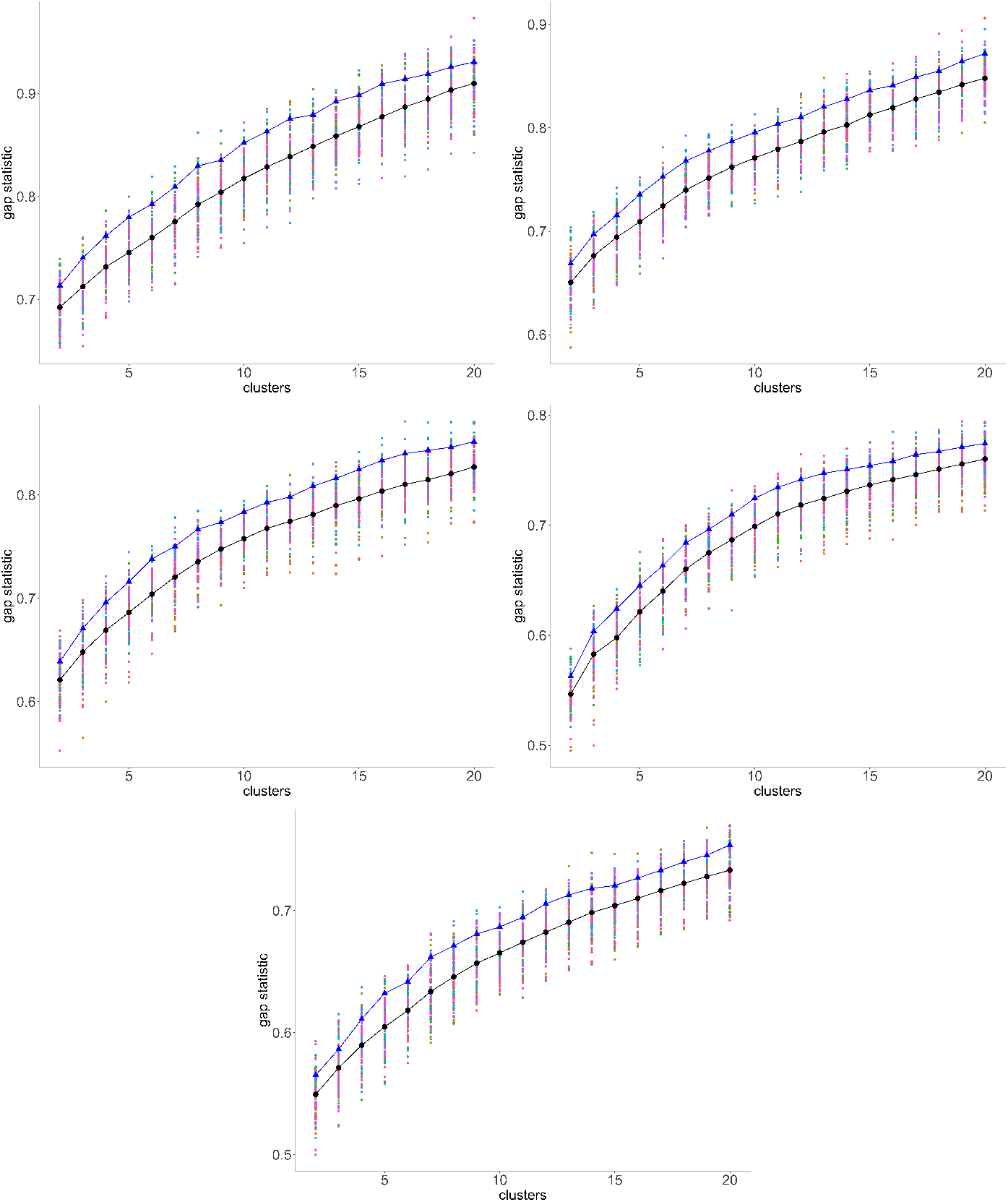
Gap statistics for Fagales for 100 samples; means and means of 10 best (in blue).

## Results

The Angiosperm Phylogeny Group circumscription of flowering plant orders and families, version IV [15], includes eight fabid and eight malvid orders (plus Vitales) as making up the rosids as well as seven campanulid and eight lamiid orders (plus Ericales) as constituting the asterids. We wished to include as many orders as possible in our study, ideally with access to at least six genomes with high quality, preferably chromosome-level, assemblies, distributed among at least three different families. At the time of data collection, we could obtain suitable data from three fabid orders, Fagales [20–27], Cucurbitales [28–33], Malpighiales [34–39]; three malvid orders, Myrtales [40–44], Malvales [45–50] and Sapindales [51–61]; one campanulid order, Asterales [62–68]; and three lamiid orders, Gentianales [69–75], Lamiales [76–82] and Solanales [83–88], plus Ericales [89–94]. Other orders with sufficient genomes available were not selected, such as Fabales, because representative genomes from only one or two families were available, or Brassicales, which is the subject of a concurrent publication.

In the case of our data, there is some uncertainty about locating the inflection point within *±*2 for any particular ancestor and any particular *g*-mer analysis. But the inflection point does not display any sensitivity to *g* in the range, say, from 15-50, although the overall gap score plot may be shifted upwards or downwards for different *g*. Further there does not seem to be any tendency for the estimate of *x* to vary from ancestor to ancestor within an order; which is understandable as the basic chromosome number would tend to be the same across a single order. More complete data appears in Supplementary materials C.

Though the gap statistics curves all display similar shapes, the increment in the significance level from *k* − 1 to *k* is subject to considerable statistical fluctuation, which is the reason for the uncertainty in determining *x*, even for the best mwm samples. To attack this problem, under the hypotheses that the choice of *g* in the range from 10-15 to 40-50 is of little consequence, and that all the ancestors in an order have the same monoploid number, we take the average of the gap statistic across all these ancestors and all the *g* as most likely to reveal the trends in the order.

In addition, to amplify the visual impression of the tendencies in gap statistic, we can plot the increment of this quantity from *k* − 1 to *k* instead of the statistic itself. We display this increment in Figure 11.

**Fig. 11.**
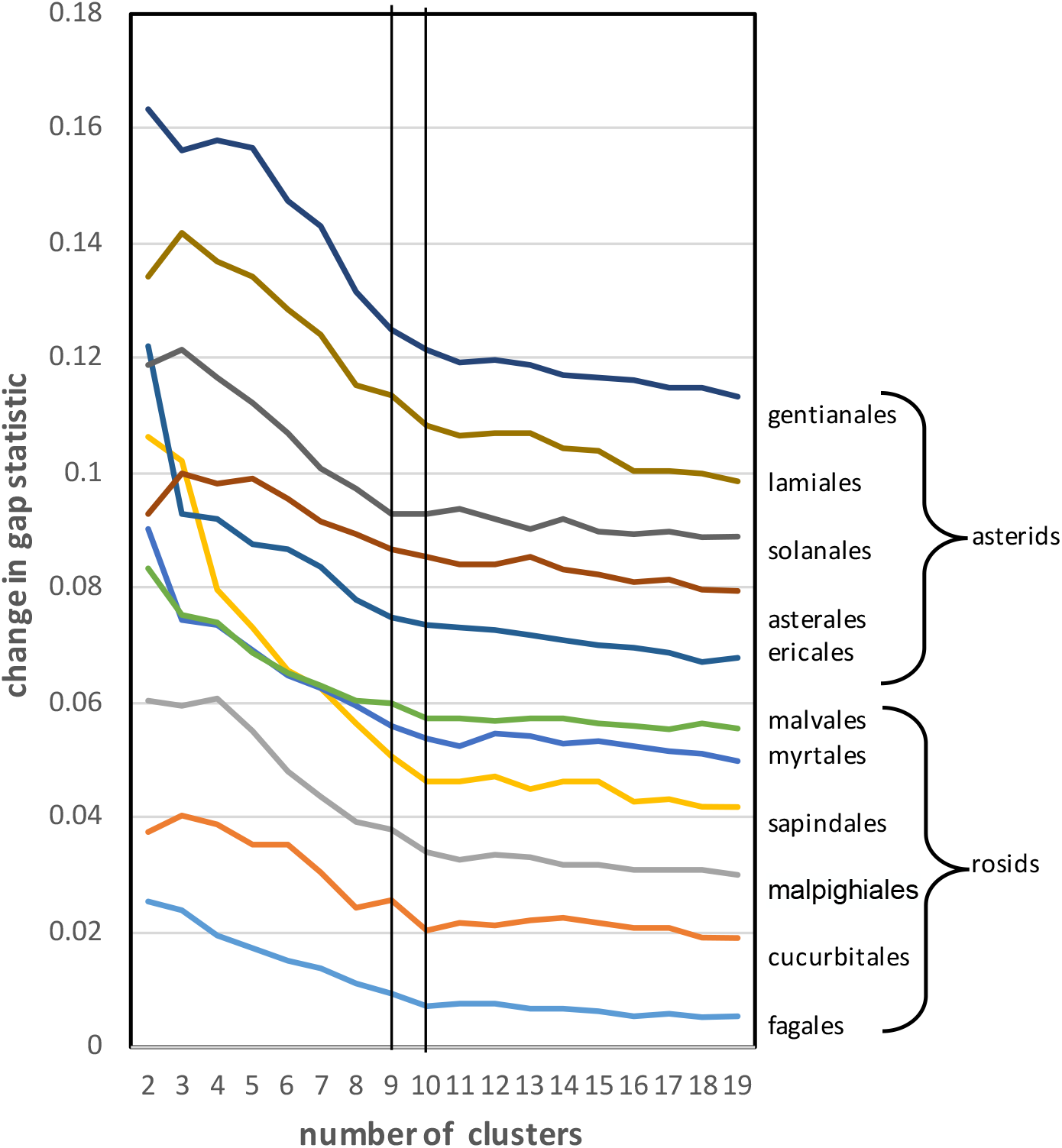
Transition between meaningful and noisy phases of increase in gap statistic for 11 core eudicot orders. The *y*-axis values are displaced +0.01 for each order in the list above the previous item. Vertical lines at *k* = 9 and *k* = 10 highlight that for most orders, the increment at *k* = 10 is in line with the trend of uninformative additional clustering at higher values of *k* while for *k* = 9 the increment exceeds this trend.

Once the ancestral reconstructions are completed, we can visualize the evolution of an extant genome from its most recent ancestor. We assign a colour to each chromosome in this ancestor, and then assign that colour to any region of an extant chromosome that matches with a contig in the ancestral chromosome. Examples are shown in Figure 12. Further examples are in the Supplementary materials D.

**Fig. 12.**
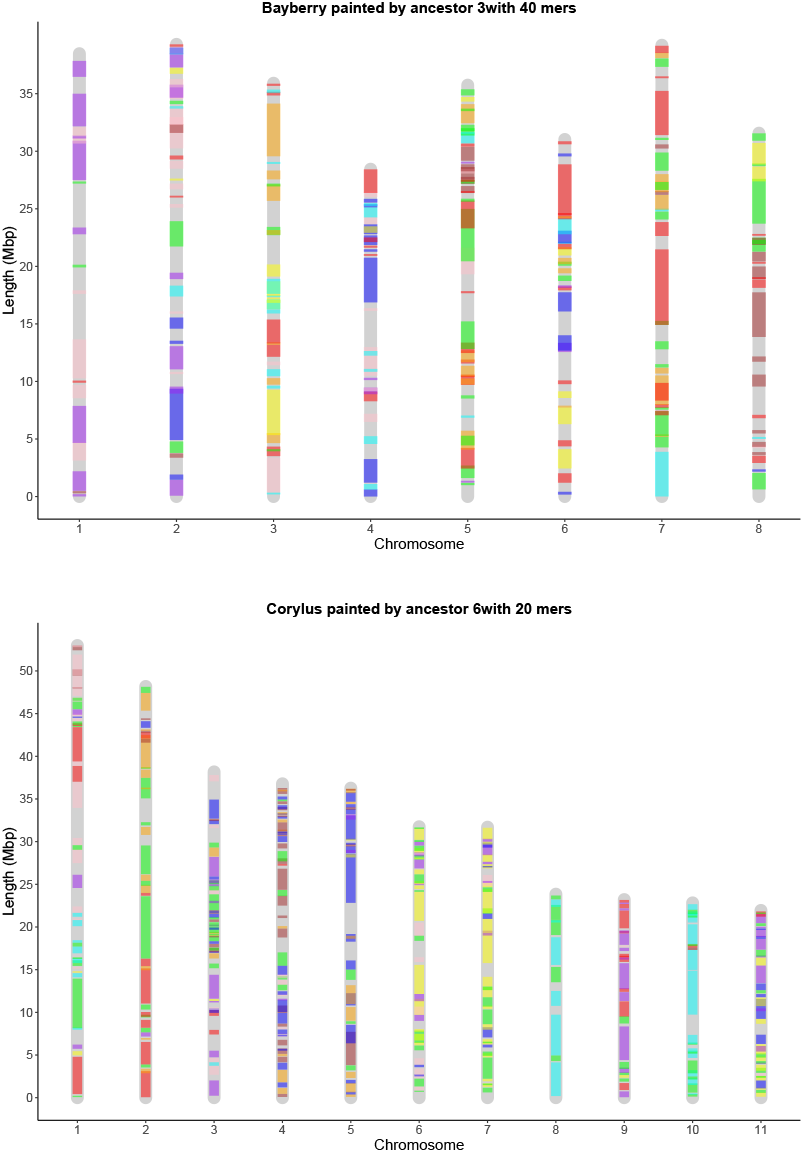
Painting the chromosomes of the extant genomes using colors corresponding to the ancestral chromosomes. Example of ancestral Fagales colors on the oak and the bayberry genomes.

The reconstruction analysis within each of the 11 core eudicot orders was carried out independently of the other orders, including the identification of the gene families. To what extent do the ancestors of the various orders resemble each other? To answer this for a particular pair of orders, we first have to determine which gene families in one order correspond to a gene family in the other. This can be done by finding pairs of genes in extant genomes that are orthologous. Once these are identified we can determine the co-occurrence of gene families across the chromosomes of the ancestral genomes in the two orders. Figure 13 gives the results of this for the Fagales and Mapighiales orders. It can be seen that for the most part, we can identify corresponding chromosomes for the two orders.

**Fig. 13.**
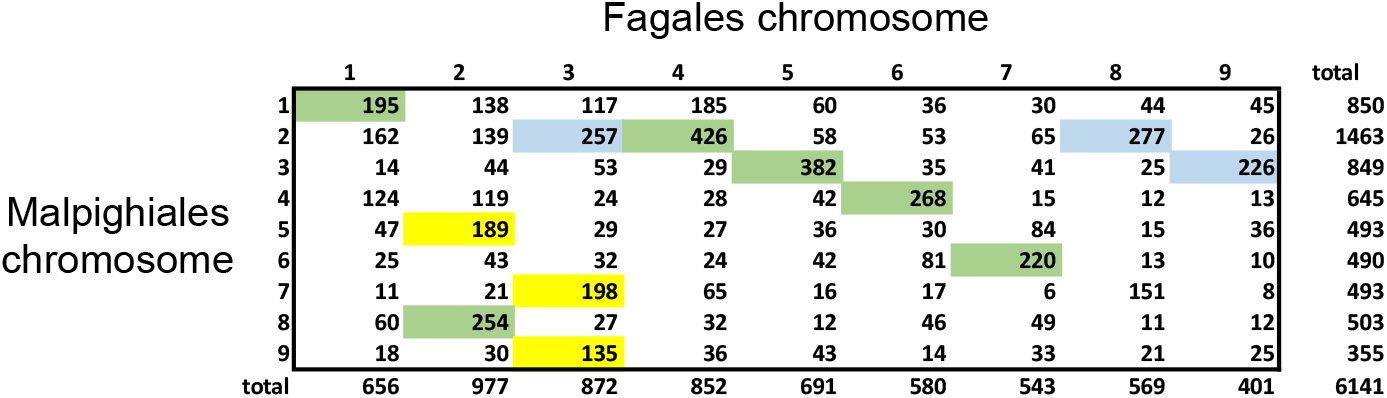
Gene families shared between Fagales and Malpighiales ancestors. For each Malpighiales ancestral chromosome, the yellow or green cell indicates the Fagales chromosome that shares a maximum number of gene families. For each Fagales ancestral chromosome, the blue or green cell indicates the Malpighiales chromosome that shares a maximum number of gene families. There are six green cells indicating closely related chromosomes in the two independently calculated ancestors. A total of 8419 gene families were reconstructed in the Malpighiales ancestor, of which 2278 were not recovered in Fagales. A total of 9424 gene families were reconstructed in the Fagales ancestor, of which 3283 were not recovered in Malpighiales.

Another aspect of the consistency of our reconstruction is a comparison with the PCA-based reconstruction. Figure 14 shows that although there are many genes that do not fit the general pattern, we can still identify, in most cases a 1-1 correspondence between the two sets of chromosomes. In the figure, 7 out of 9 chromosomes correspond in this way.

**Fig. 14.**
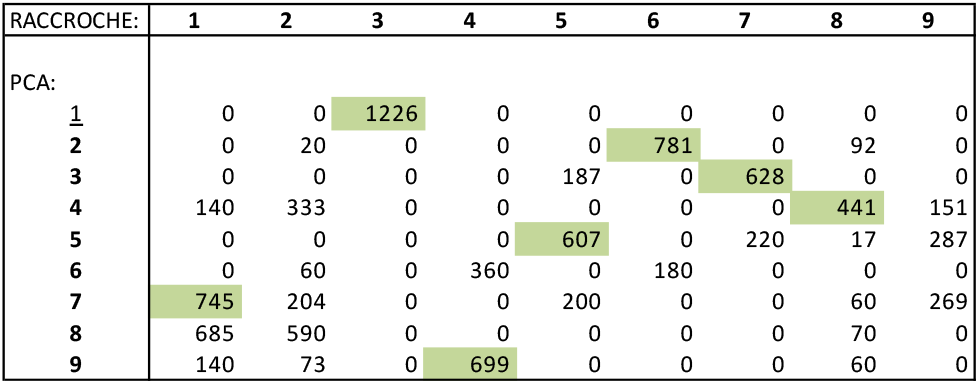
Gene families shared between hierarchical clustering-based chromosomes and PCA-based chromosomes. A green cell indicates that a chromosome from one method shares a maximum number of gene families with a single chromosome from the other. method.

### No upper limit on x

Our reconstruction of monoploid ancestors of rosid ancestors, not only at the highest nodes, but for somewhat more recent ancestors, all seem to have monoploid number *k* ≤ 9. These results are eminently plausible, but still may provoke the question of whether Raccroche would even be able to detect a higher *k* for an ancestor if this were warranted.

Lacking any knowledge of ground truth about ancient plant karyotypes, we could have recourse to simulations, and simulation protocols have been used successfully in studying plant evolution [8]. But simulations of plant evolution starting with an *x* = 20, say, ancestor, and known parameters for chromosome fusion, whole genome duplication, and other processes, could only be very speculative, generating unrealistic versions of extant genomes to test the Raccroche reconstruction.

Instead, we venture outside the plant world to an evolutionary domain where the monoploid ancestor is agreed to have *x* around 20, namely the animals, or metazoans [95]. We used two out of the five genomes from those studied in [95], namely the lancelet *Branchiostoma floridea*, representing the deuterostomes, and the sponge *Ephydatia muelleri* [96]. The annotated genome files from the other three species in [95] being publicly unavailable, we substituted *Octopus sinensis* [97] from the phylum Mollusca as a representative of the protostomes, the cnidarian *Acropora millepora* [98] and the placozoan *Trichoplax adhaerans* [99]. These five species represent major branches of the animal kingdom, including the subkingdoms Porifera (sponges) and Eumetazoa, the latter branching into the placozoan and cnidarian phyla and the bilaterians protostomes and deuterostomes, as in Figure 15. We note that the time scale is 6 to 10 times as long as that of the plant orders we have focused on. Not surprisingly, given the well-known lack of conserved gene order among early metazoan lineages [100], Raccroche produces relatively short contigs with these data.

**Fig. 15.**
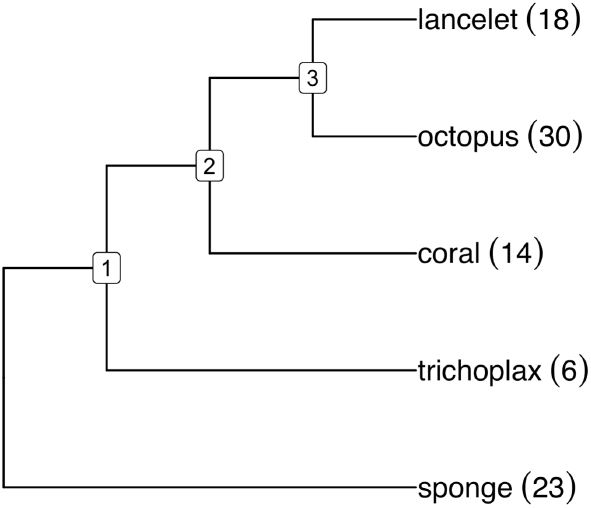
Metazoan phylogeny with haploid numbers of chromosomes

The results of our analysis is summarized in the significance increment graph Figure 16. Here the monoploid number appears to be between 15 and 21.

**Fig. 16.**
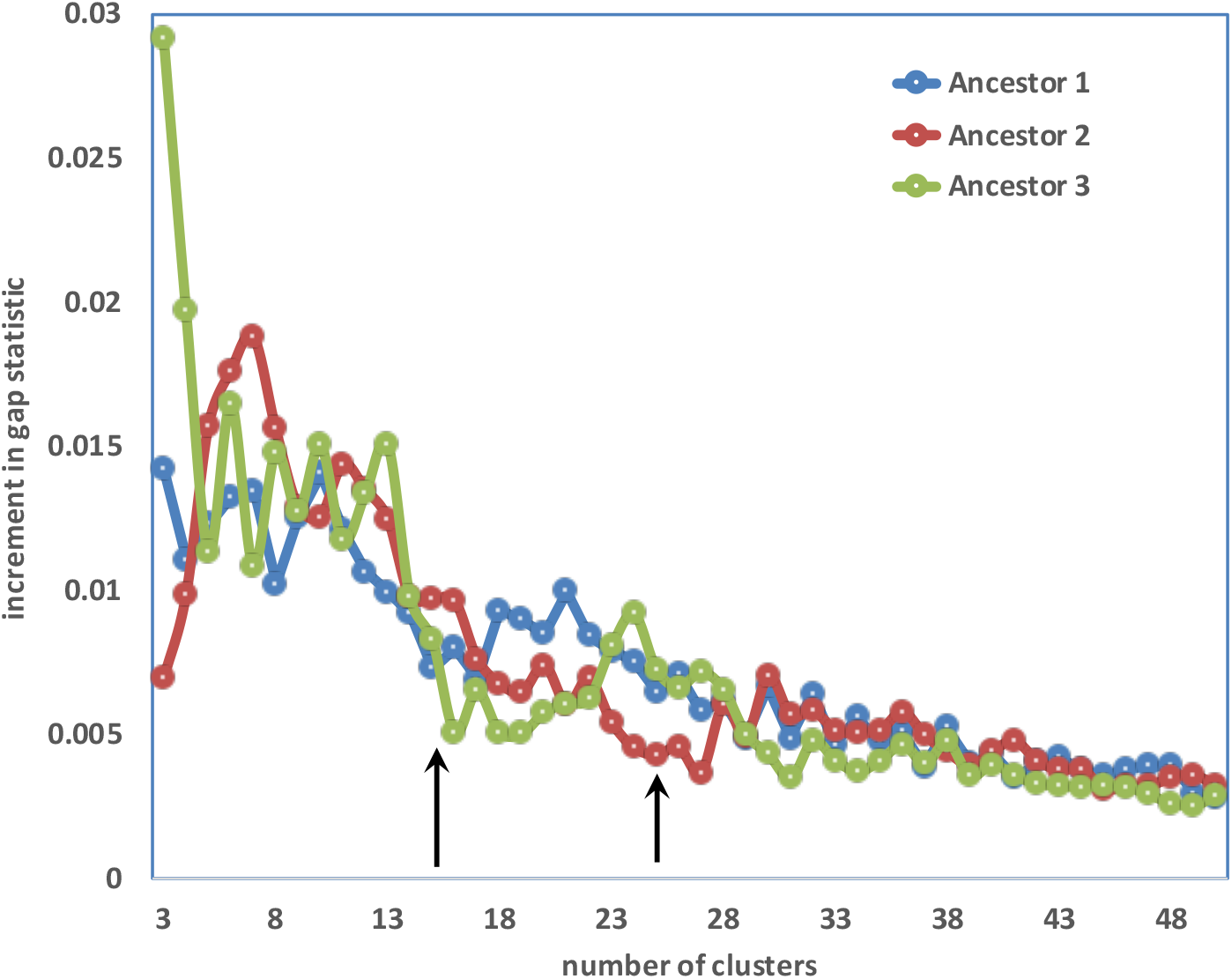
Significance increment graph for cluster-based metazoan karyotypes

## Discussion

Grant’s visionary work on basic chromosome number of ancestral plants [1] predated genomics by several decades, but made use of data on many thousands of species to produce excellent estimates. Modern genome-free approaches [2–4] use sophisticated statistical methodology on greatly expanded data sets to improve and automate this line of research.

With the rise of molecular approaches to evolution and genomics, however, it behooves us to investigate whether the gene order on the chromosomes of a set or related extant genomes carry a signal about the basic chromosome number.

Despite their demonstrated ability to estimate gene content and to some extent gene order in reconstructed ancient genomes, the problem of delimiting chromosomes in an automated way has proved difficult [5]. Our approach differs from previous methods in that it focuses solely on monoploid reconstructions, whether or not this corresponds to the ploidy of the hypothesized ancestors. This is done in a purely automated way, given the gene orders on the chromosomes or scaffolds of extant genomes, as well as their phylogenetic relationships, without taking into account supplementary information or hypotheses in the process. The use of mwm, chromosomal co-occurrence matrices and gap statistics to achieve the monoploid reconstruction is entirely novel.

The result of applying our method to eleven rosid and asterid orders, without directly referencing chromosome number of the extant genomes, is that the basic chromosome number of these core eudicots is nine. This is somewhat higher than the value of eight recently obtained by genome-free methods [4] using chromosome numbers of many thousands of extant species, but not at all inconsistent with Grant’s original assessment [1, p.486].

Our reconstructions must be considered estimates; we only recover around 10,000 gene families, less than half of what we expect from plant genomes, even those which have not undergone whole genome duplication. Second the assignment of these “genes” to chromosomes, and their ordering along the chromosomes varies somewhat from one mwm to another, even among those which results in the clearest clustering. Nevertheless, every stage of our pipeline, which is not influenced by any information or data outside the input genomes, produces global or locally optimal results. Moreover, we have several indications of consistency, including chromosome-by-chromosome correspondences among the ancestors from different core eudicot orders, which are constructed independently from entirely different genomes. This is also clear from the parallel plots of gap statistics increments and the switch between meaningful increase and noisy increase. We also have correspondences between the results from different clustering methods. Furthermore we know that these results are not an artifact of some limitation in the detection power of our method; it successfully estimated an *x* value twice as large as the core eudicots for the ancestral metazoan genomes. This work opens up new directions for research into the evolution of the chromosomal structures of plants and other organisms.

