## Supplements for "From comparative gene content and gene order to ancestral contigs, chromosomes and karyotypes"

***A. Heat maps, Eleven orders, all ancestors*** $g = 20$

***B. Evaluation graphs. Eleven orders***

***C. Gap statistics. For*** $g = 20$ ***each ancestor, eleven orders***

***D. Painted extant genomes*** $g = 20$

***E. PCA clustering*** $g = 20$

### Supplement A. Heat maps, Eleven orders, all ancestors $g = 20$

Heat maps also available for 15-mers, 30-mers and 40-mers.

**Ancestor 1 with 40-Mers**

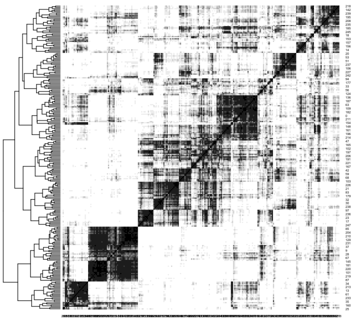

**Ancestor 2 with 40-Mers**

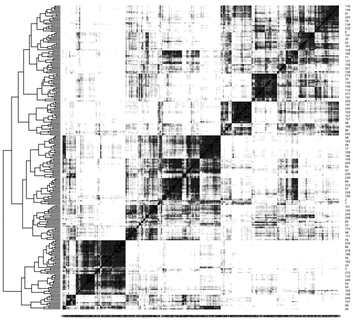

**Ancestor 3 with 40-Mers**

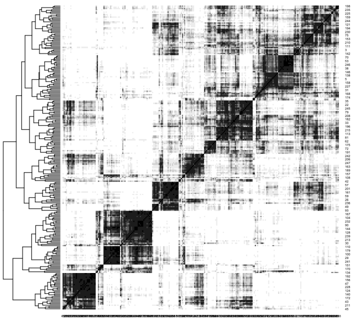

**Ancestor 4 with 40-Mers**

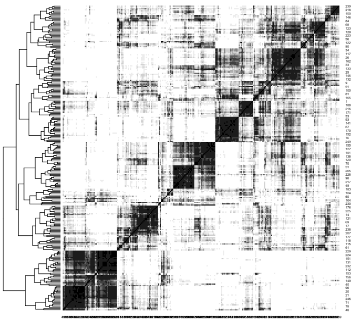

**Ancestor 5 with 40-Mers**

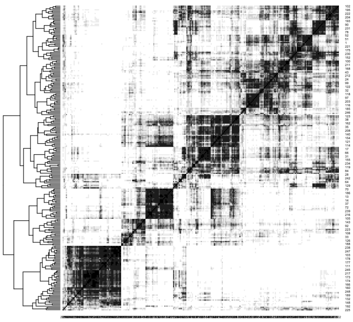

**Ancestor 6 with 40-Mers**

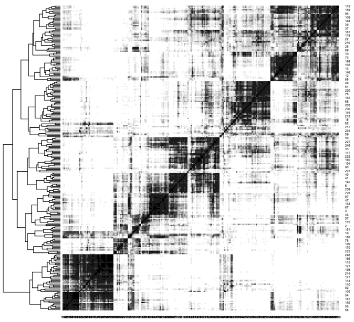

**Ancestor 7 with 40-Mers**

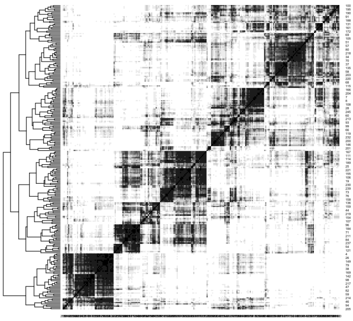

### Fagales

Ancestor 1 with 20-Mers

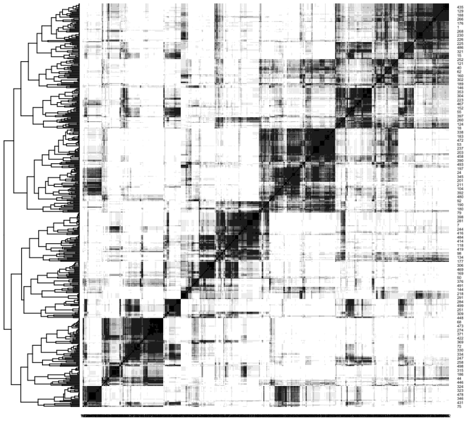

Ancestor 2 with 20-Mers

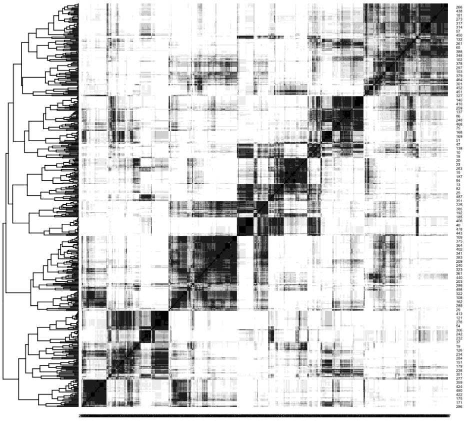

Ancestor 3 with 20-Mers

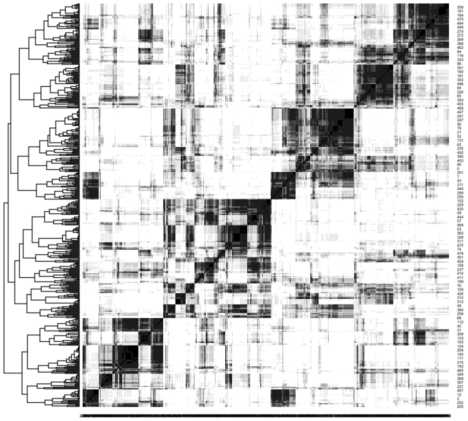

Ancestor 4 with 20-Mers

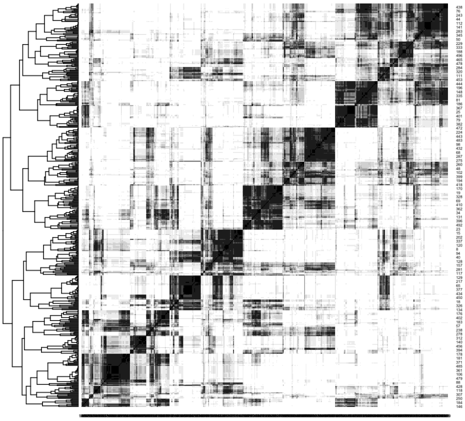

Ancestor 5 with 20-Mers

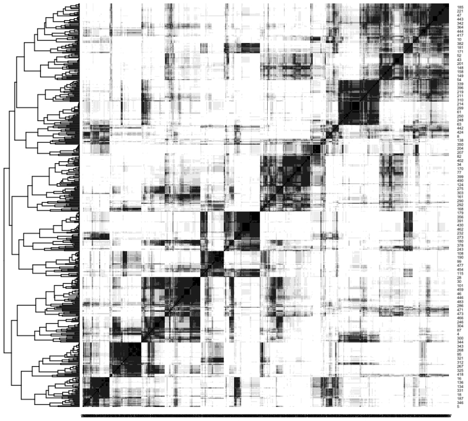

### Cucurb

**Ancestor 1 with 30-Mers**

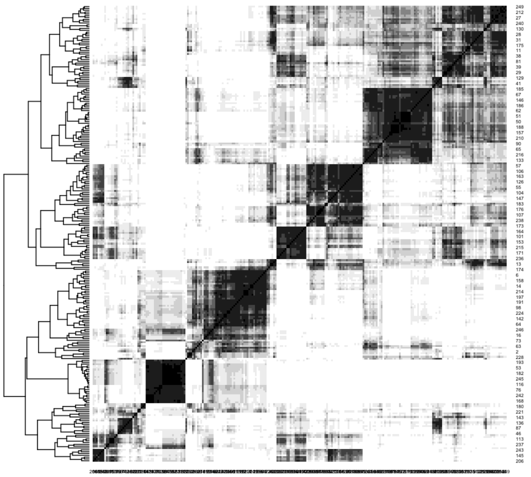

**Ancestor 2 with 30-Mers**

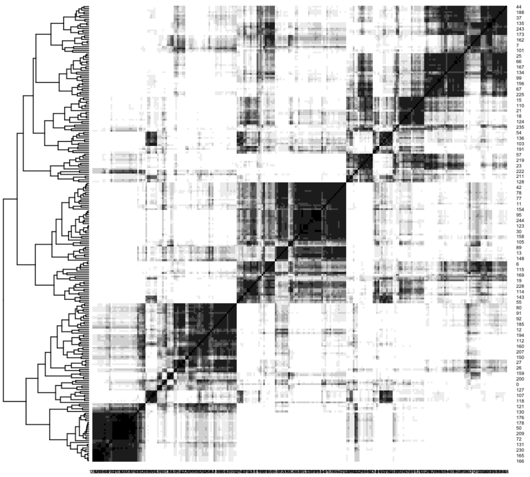

**Ancestor 3 with 30-Mers**

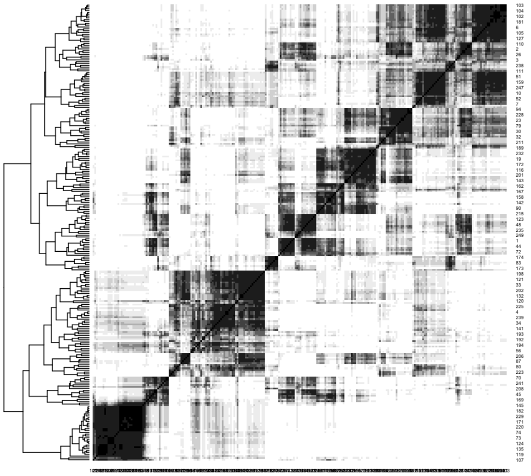

**Ancestor 4 with 30-Mers**

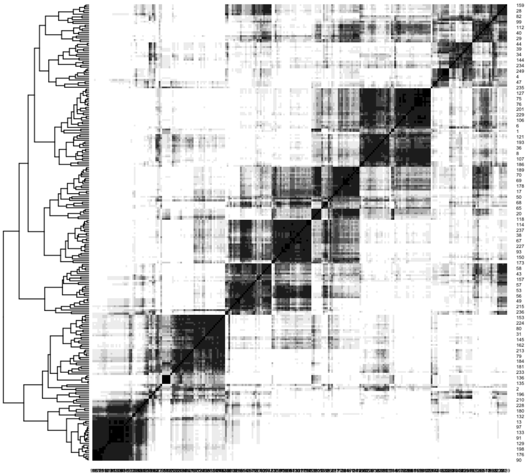

### Malpighiales

**Ancestor 1 with 20-Mers**

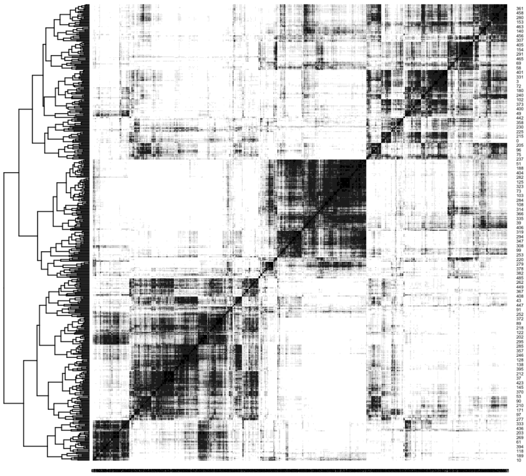

**Ancestor 2 with 20-Mers**

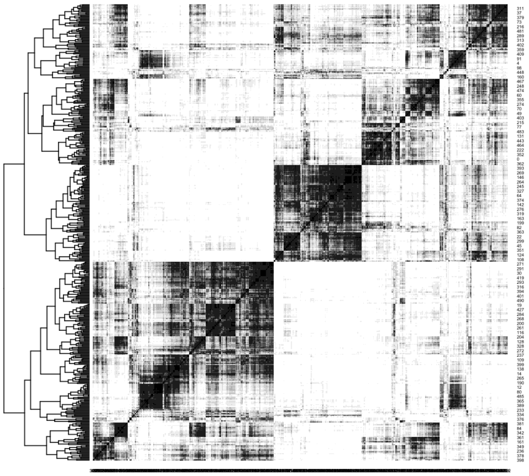

**Ancestor 3 with 20-Mers**

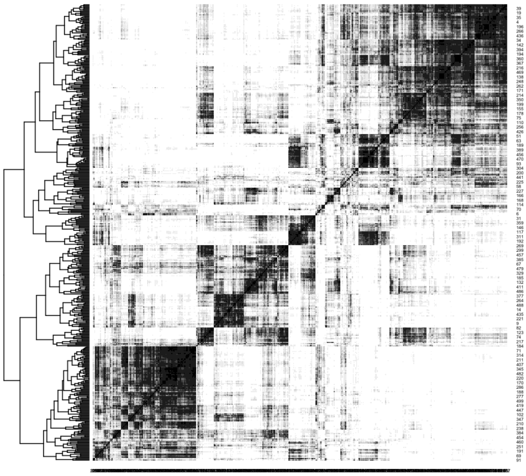

### Myrtales

##### Ancestor 1 with 30-Mers

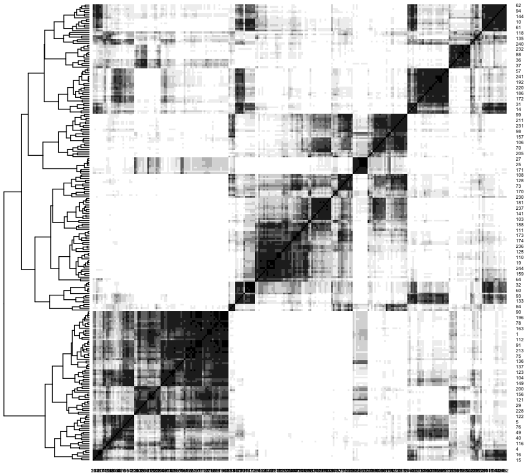

##### Ancestor 2 with 30-Mers

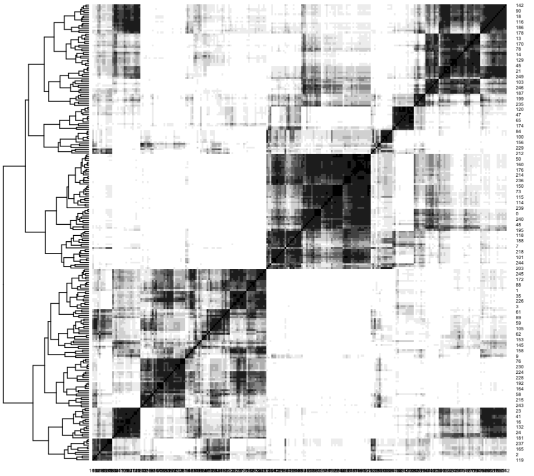

##### Ancestor 3 with 30-Mers

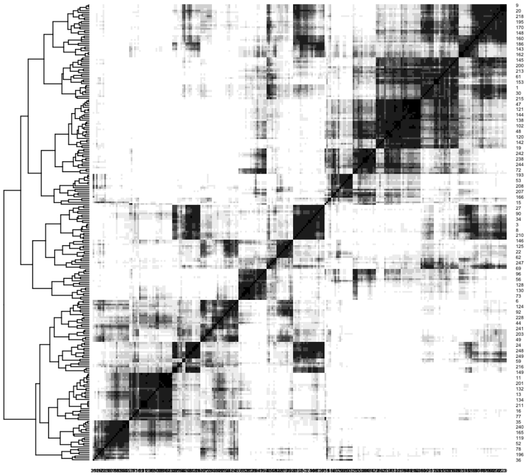

##### Ancestor 4 with 30-Mers

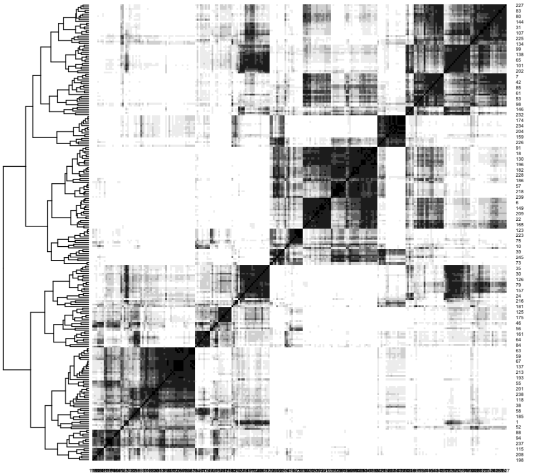

### Malvales

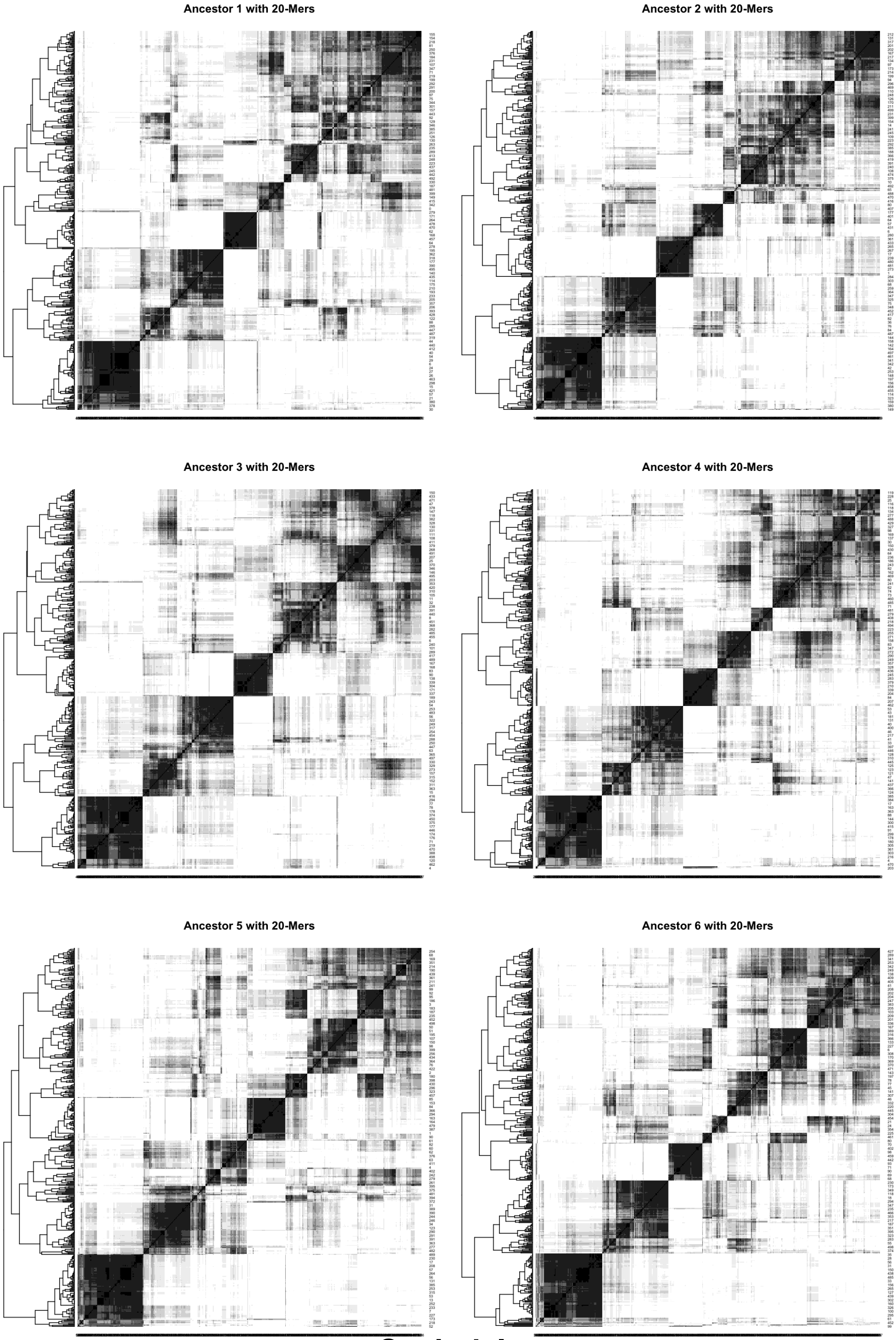
Ancestor 1 with 20-Mers
Ancestor 2 with 20-Mers
Ancestor 3 with 20-Mers
Ancestor 4 with 20-Mers
Ancestor 5 with 20-Mers
Ancestor 6 with 20-Mers

### Sapindales

**Ancestor 1 with 20-Mers**

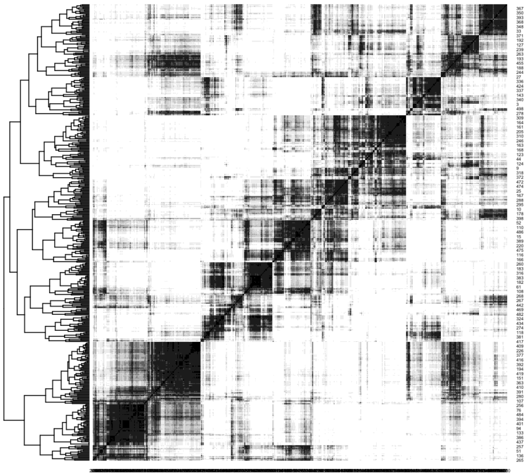

**Ancestor 2 with 20-Mers**

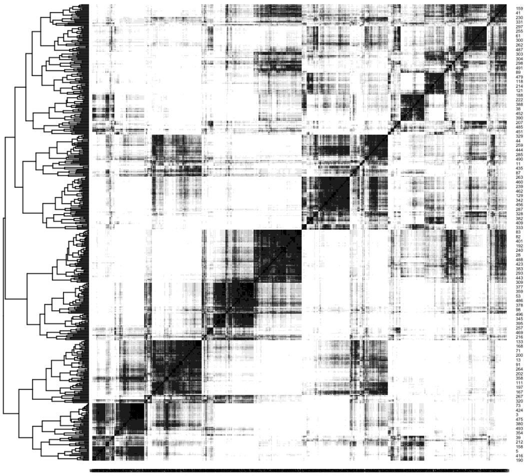

**Ancestor 3 with 20-Mers**

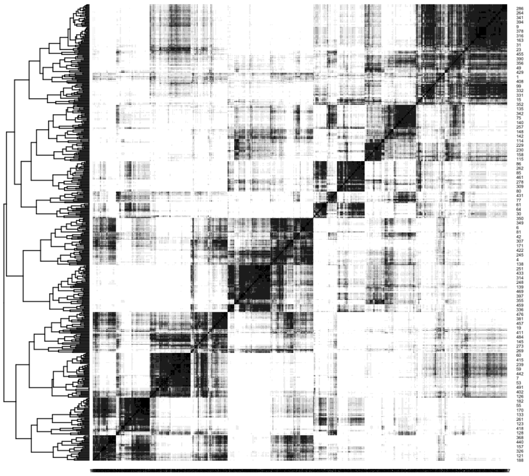

**Ancestor 4 with 20-Mers**

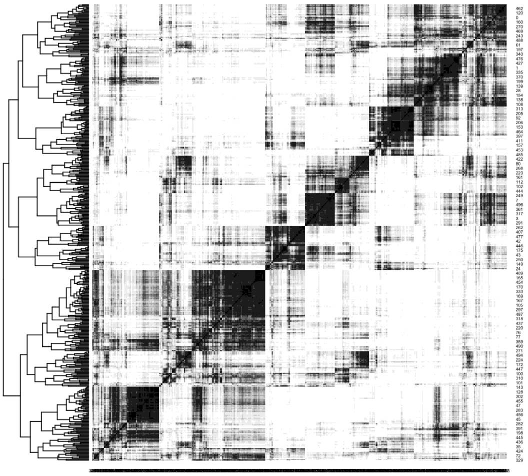

### Asterales

**Ancestor 1 with 40-Mers**

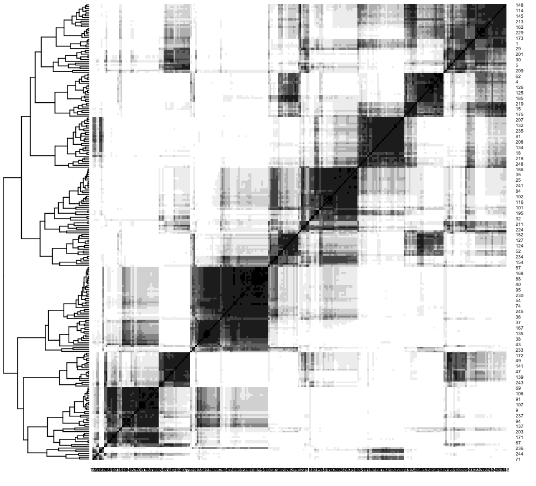

**Ancestor 2 with 40-Mers**

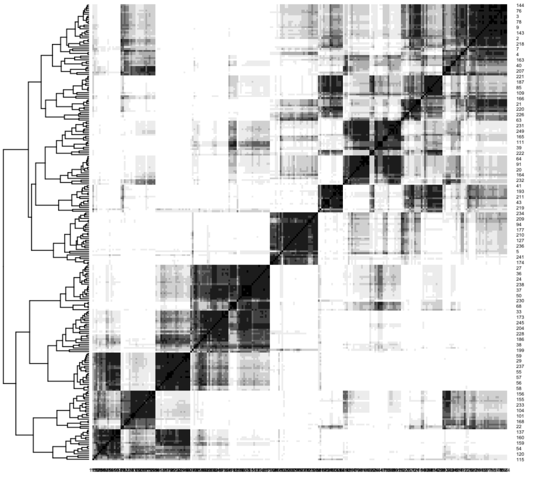

**Ancestor 3 with 40-Mers**

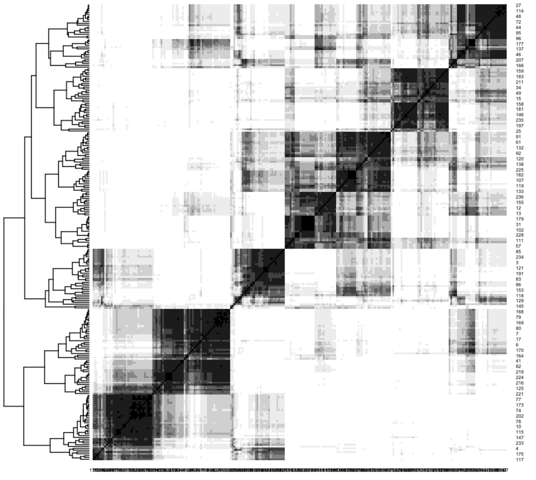

**Ancestor 4 with 40-Mers**

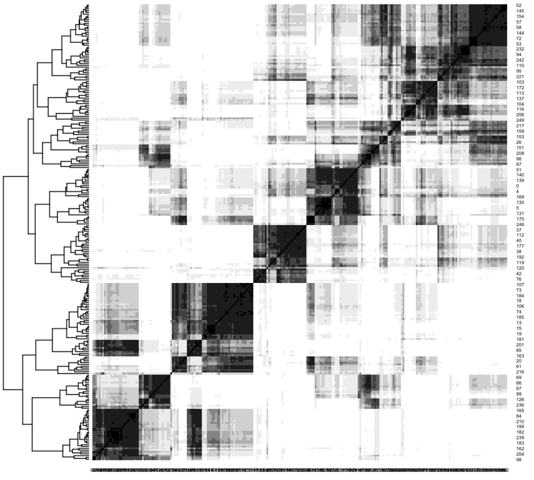

### Gentianales

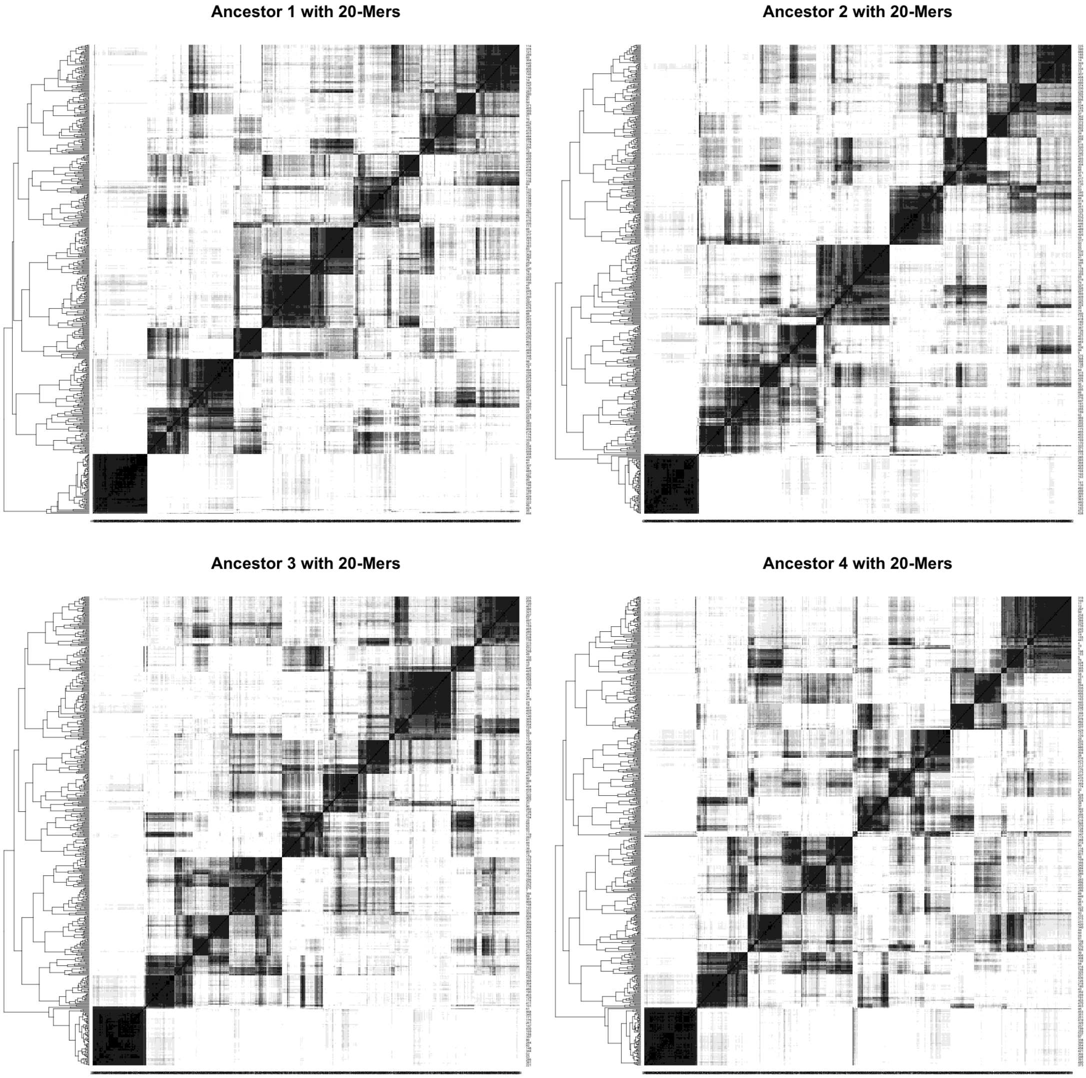
Ancestor 1 with 20-Mers
Ancestor 2 with 20-Mers
Ancestor 3 with 20-Mers
Ancestor 4 with 20-Mers

**Lamiales**

**Ancestor 1 with 20-Mers**

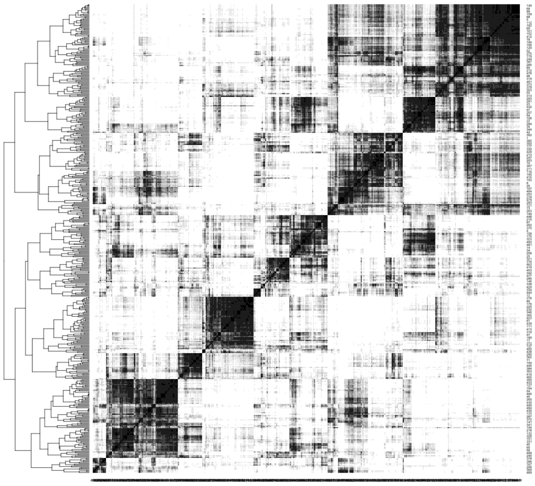

**Ancestor 2 with 20-Mers**

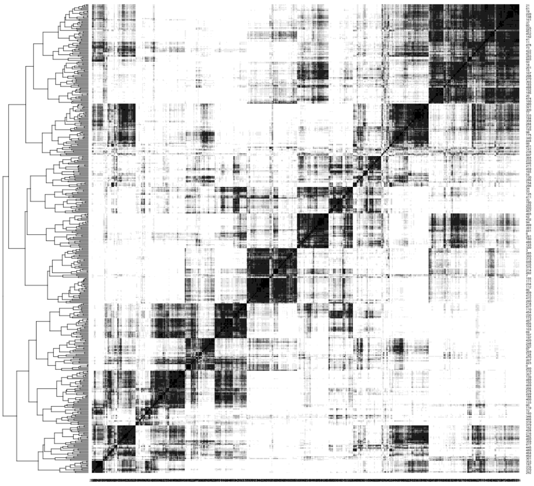

**Ancestor 3 with 20-Mers**

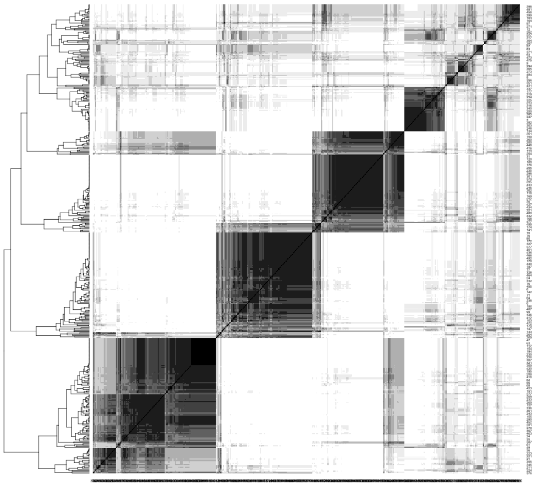

**Ancestor 4 with 20-Mers**

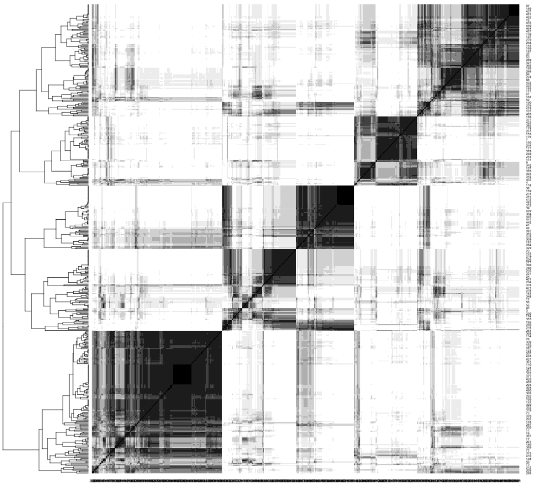

### Solanales

Ancestor 1 with 20-Mers

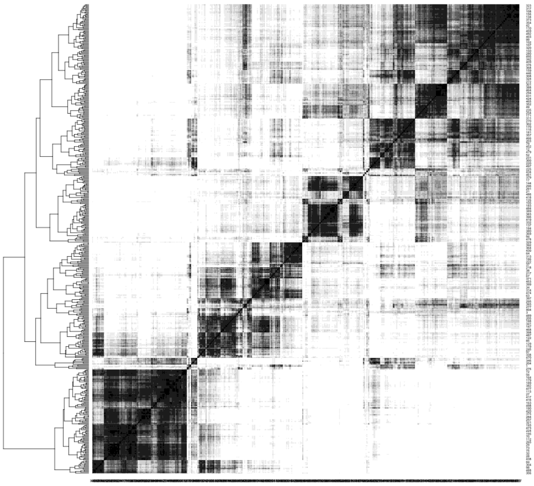

Ancestor 2 with 20-Mers

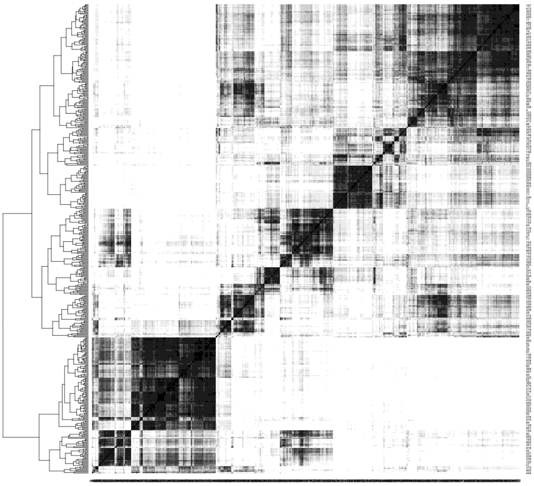

Ancestor 3 with 20-Mers

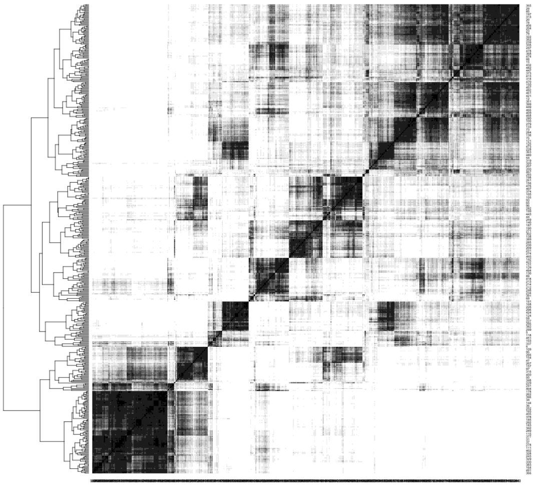

Ancestor 4 with 20-Mers

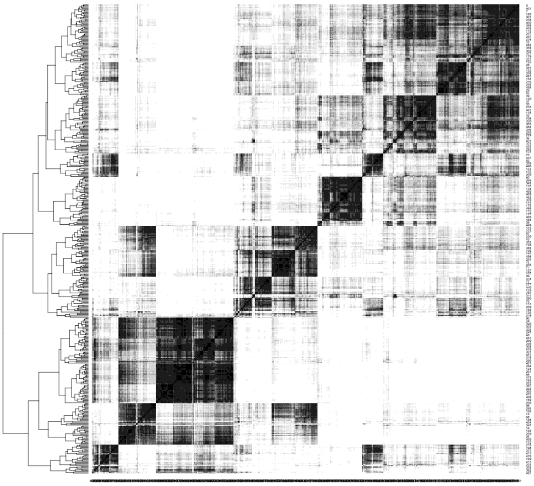

### Ericales

Springer Nature 2021 LaTeX template

### Supplement B. Evaluation graphs. Eleven orders

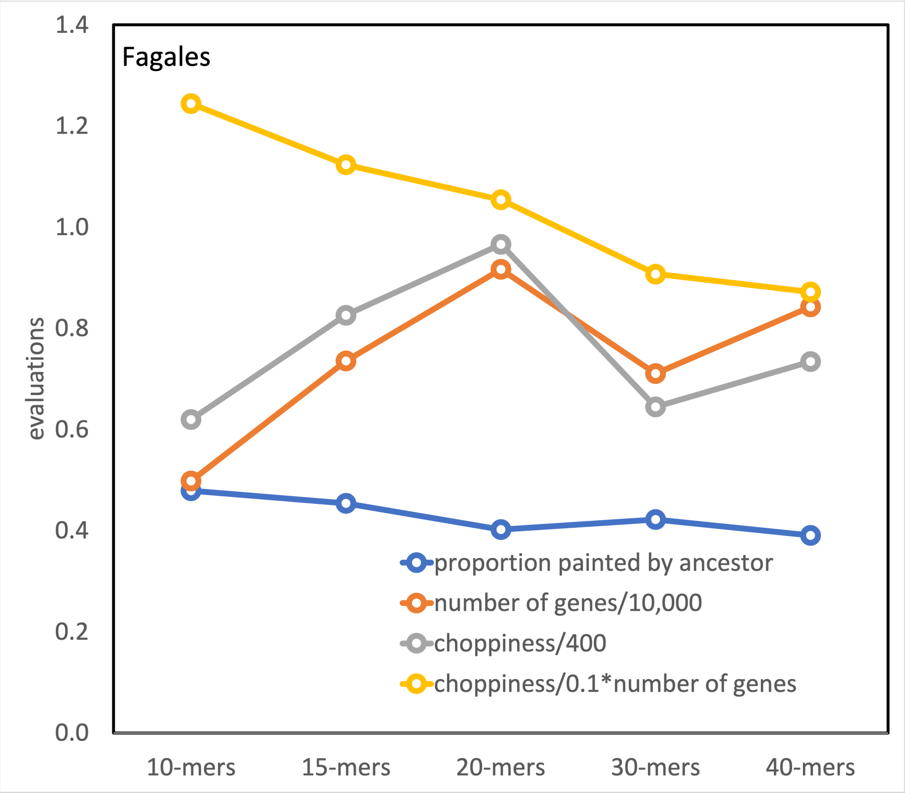

Fagales
evaluations
1.4
1.2
1.0
0.8
0.6
0.4
0.2
0.0
10-mers
15-mers
20-mers
30-mers
40-mers
proportion painted by ancestor
number of genes/10,000
choppiness/400
choppiness/0.1*number of genes

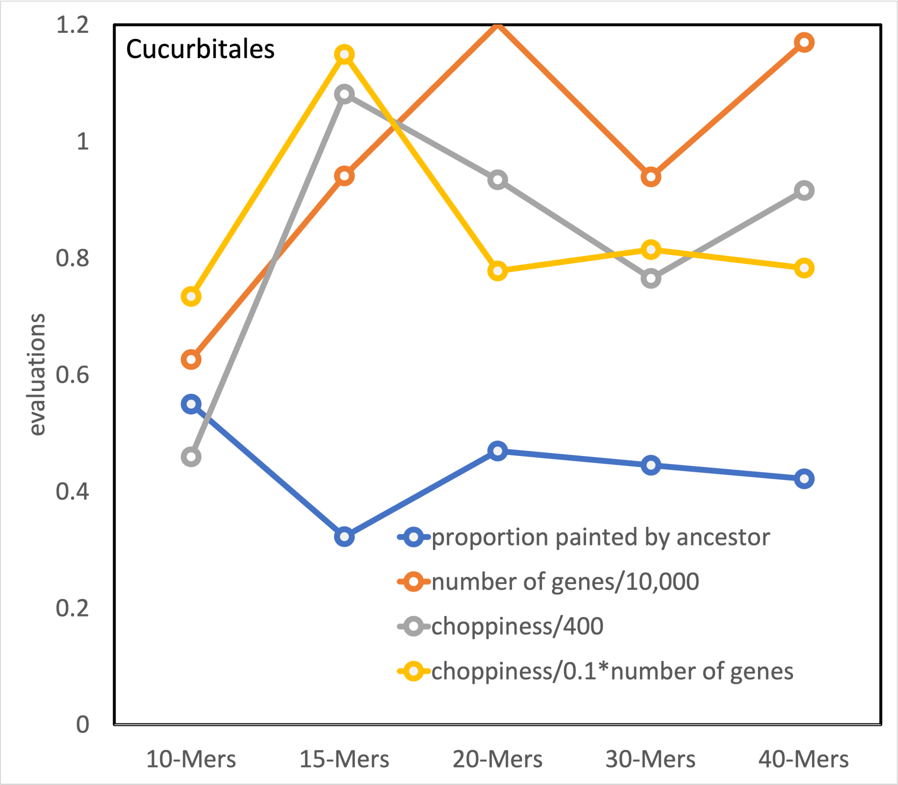
Cucurbitales
evaluations
1.2
1
0.8
0.6
0.4
0.2
0
10-Mers
15-Mers
20-Mers
30-Mers
40-Mers
proportion painted by ancestor
number of genes/10,000
choppiness/400
choppiness/0.1*number of genes

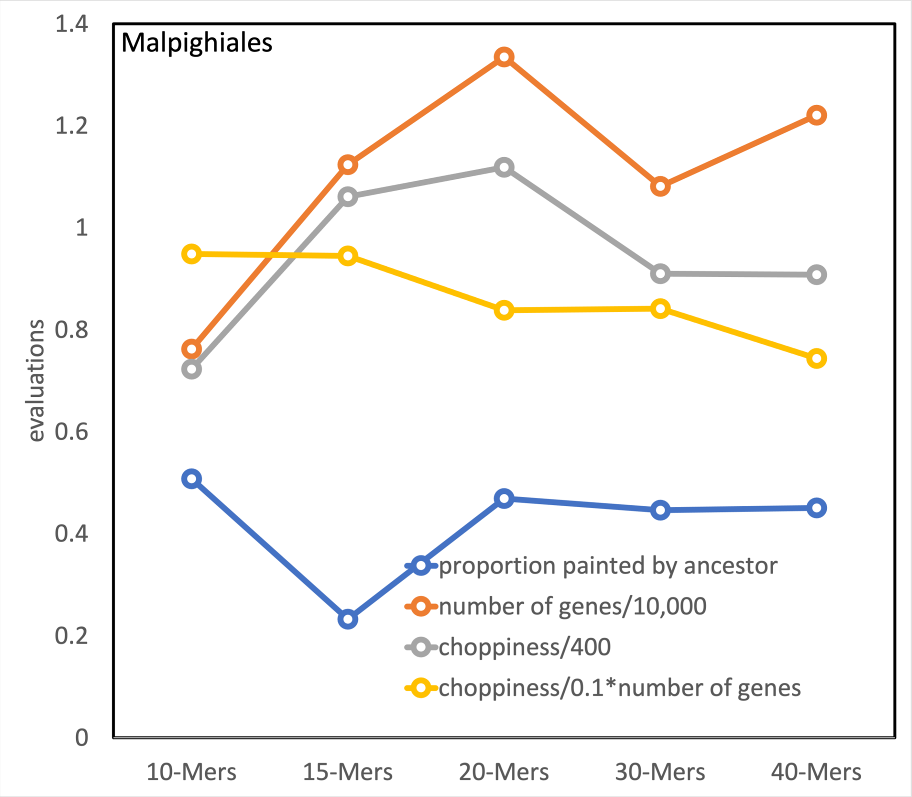
Malpighiales
1.4
1.2
1
0.8
0.6
0.4
0.2
0
evaluations
10-Mers
15-Mers
20-Mers
30-Mers
40-Mers
proportion painted by ancestor
number of genes/10,000
choppiness/400
choppiness/0.1*number of genes

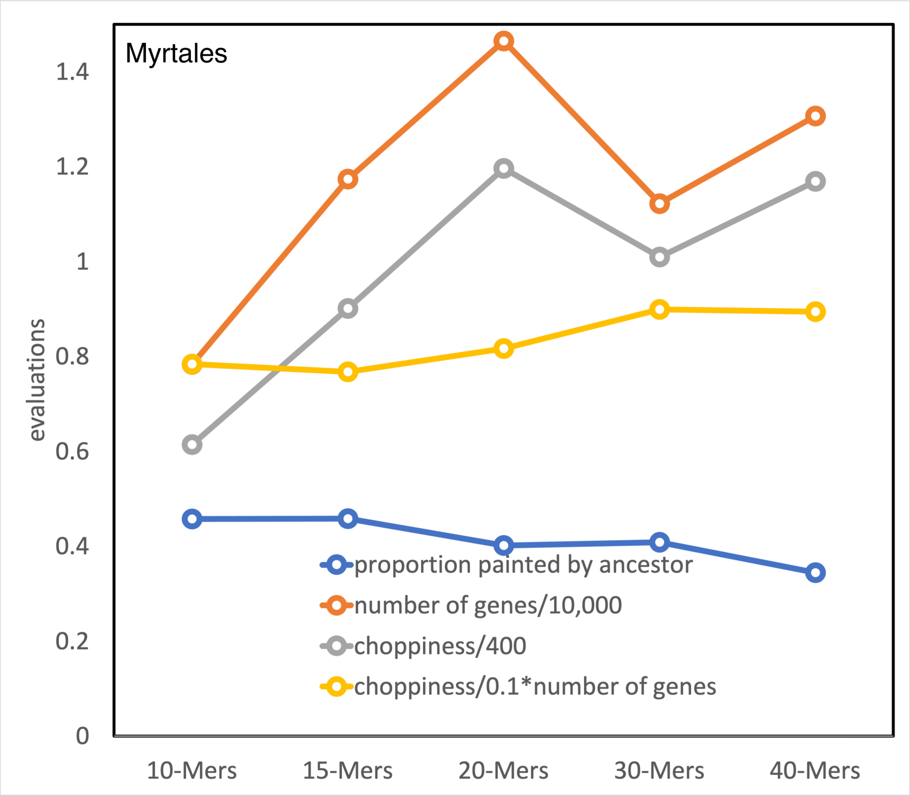
Myrtales
evaluations
1.4
1.2
1
0.8
0.6
0.4
0.2
0
10-Mers
15-Mers
20-Mers
30-Mers
40-Mers
proportion painted by ancestor
number of genes/10,000
choppiness/400
choppiness/0.1*number of genes

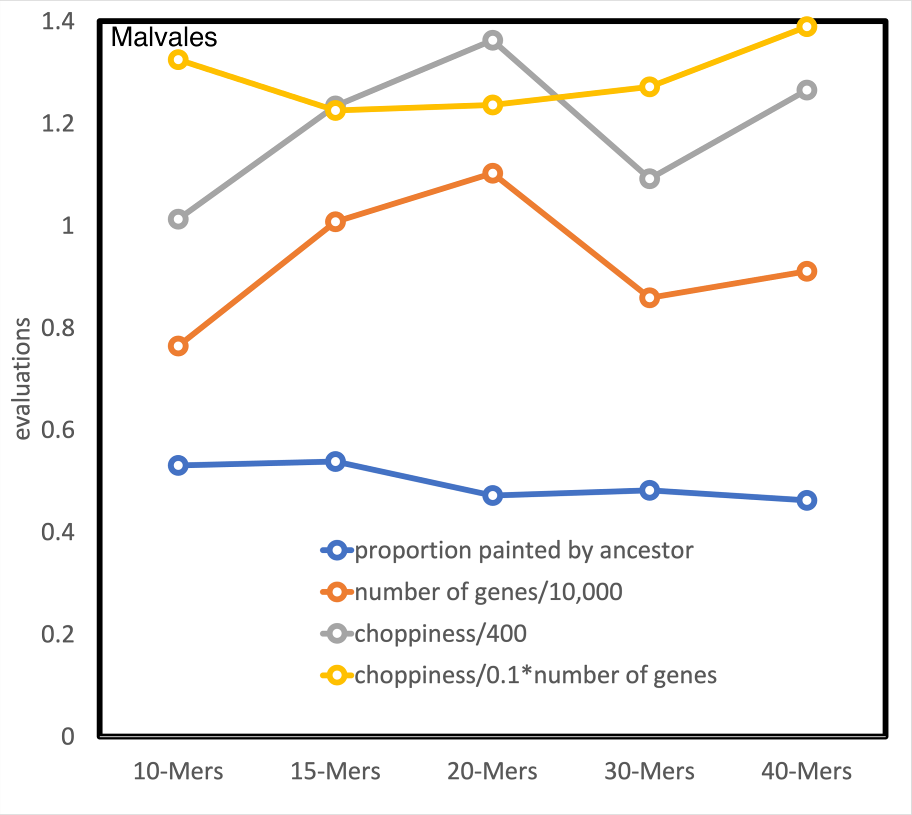
Malvales
evaluations
1.4
1.2
1
0.8
0.6
0.4
0.2
0
10-Mers
15-Mers
20-Mers
30-Mers
40-Mers
proportion painted by ancestor
number of genes/10,000
choppiness/400
choppiness/0.1*number of genes

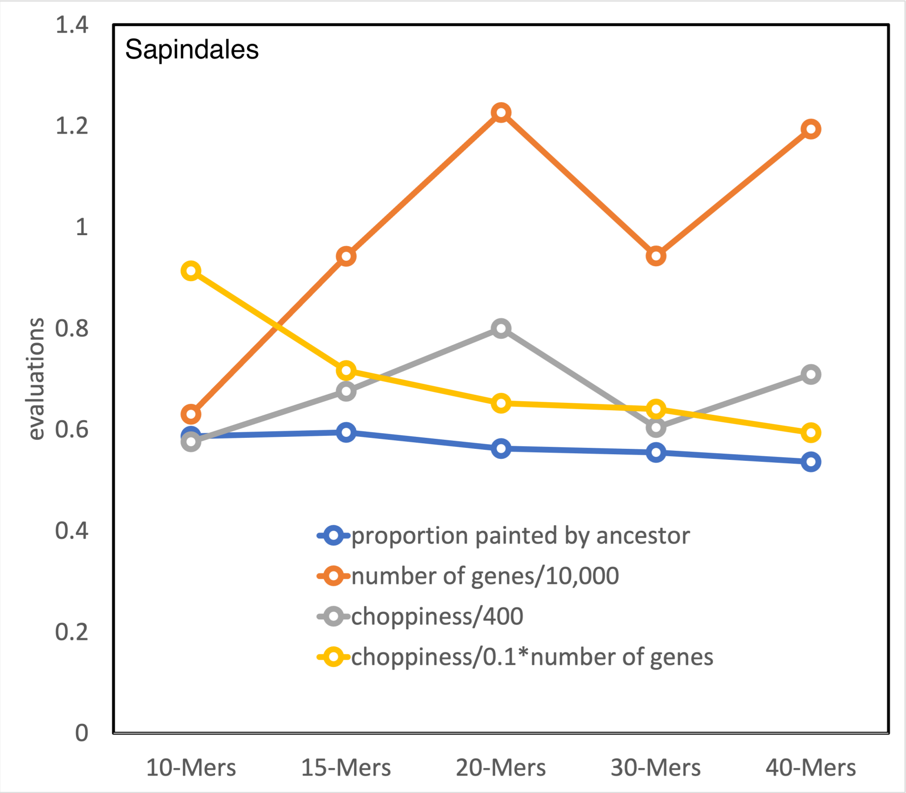

Sapindales
evaluations
1.4
1.2
1
0.8
0.6
0.4
0.2
0
10-Mers
15-Mers
20-Mers
30-Mers
40-Mers
proportion painted by ancestor
number of genes/10,000
choppiness/400
choppiness/0.1*number of genes

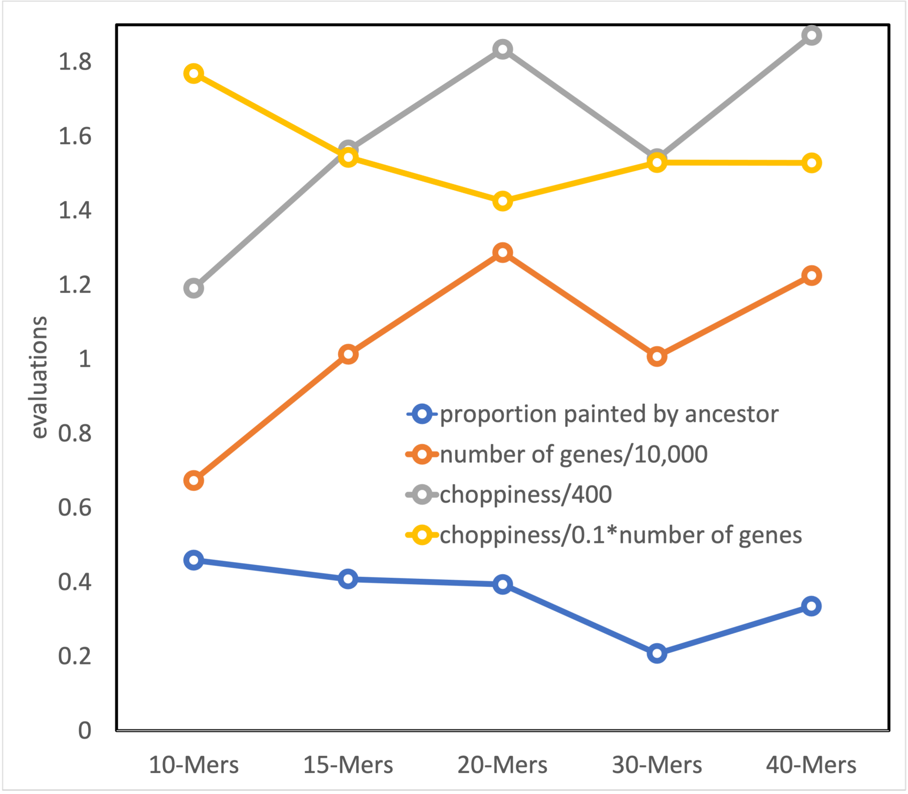

evaluations
1.8
1.6
1.4
1.2
1
0.8
0.6
0.4
0.2
0
10-Mers
15-Mers
20-Mers
30-Mers
40-Mers
proportion painted by ancestor
number of genes/10,000
choppiness/400
choppiness/0.1*number of genes

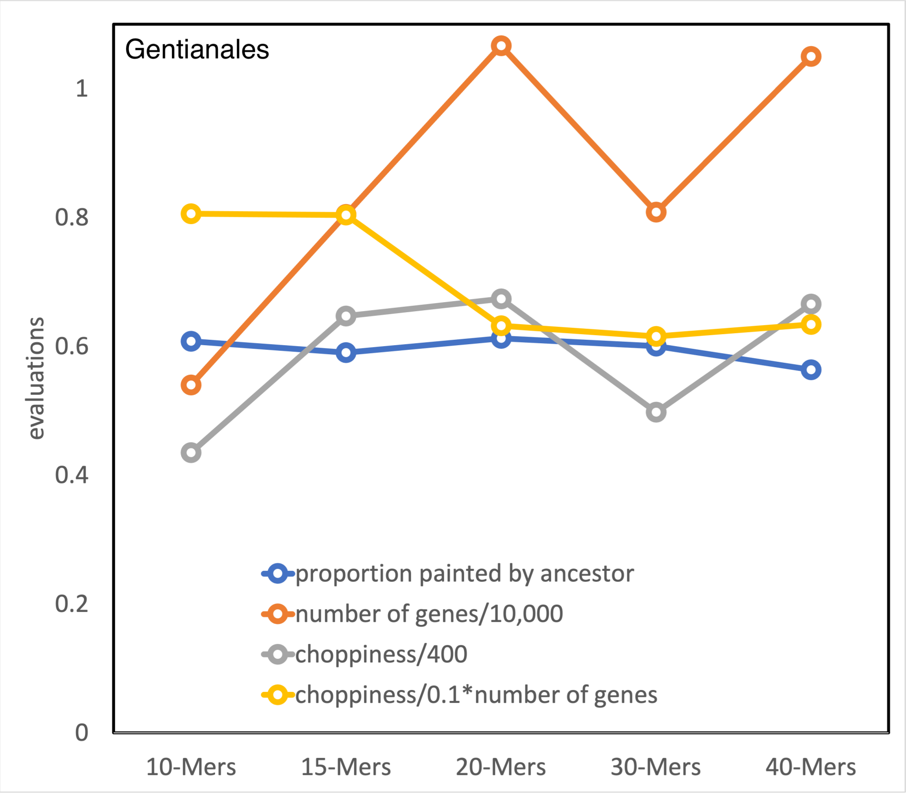

Gentianales
evaluations
1
0.8
0.6
0.4
0.2
0
10-Mers
15-Mers
20-Mers
30-Mers
40-Mers
proportion painted by ancestor
number of genes/10,000
choppiness/400
choppiness/0.1*number of genes

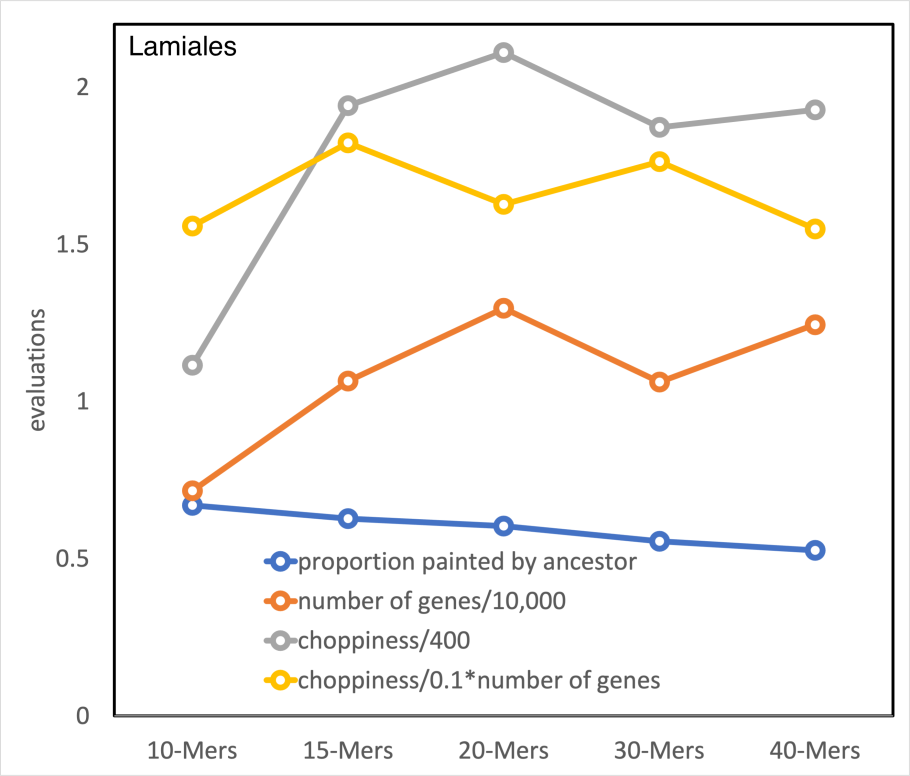

Lamiales
evaluations
2
1.5
1
0.5
0
10-Mers
15-Mers
20-Mers
30-Mers
40-Mers
proportion painted by ancestor
number of genes/10,000
choppiness/400
choppiness/0.1*number of genes

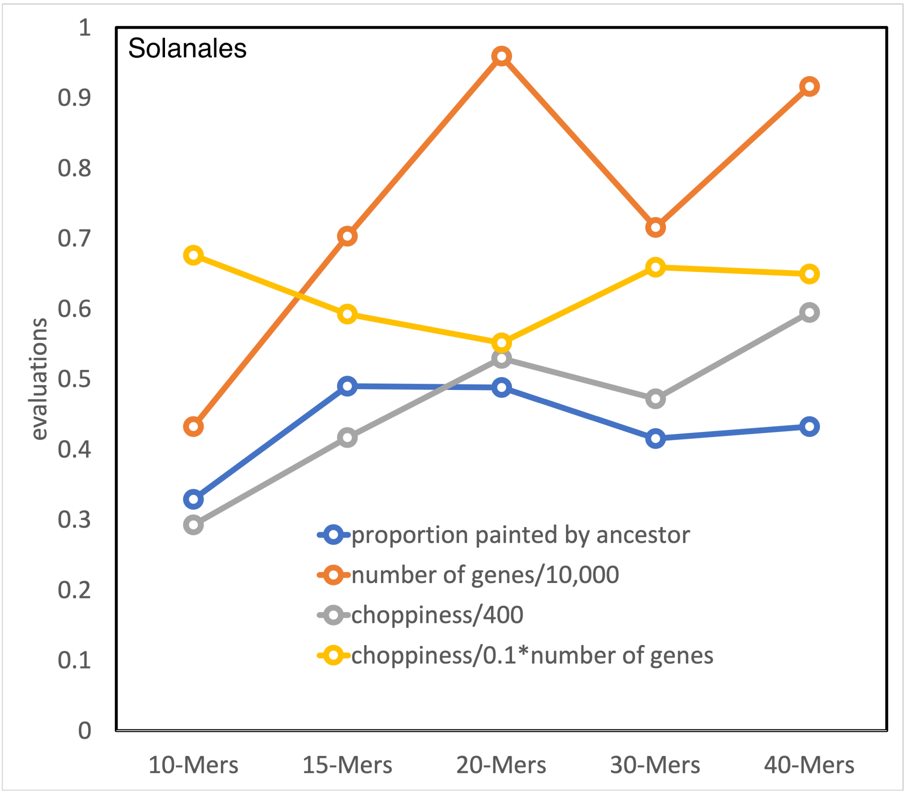

Solanales
evaluations
1
0.9
0.8
0.7
0.6
0.5
0.4
0.3
0.2
0.1
0
10-Mers
15-Mers
20-Mers
30-Mers
40-Mers
proportion painted by ancestor
number of genes/10,000
choppiness/400
choppiness/0.1*number of genes

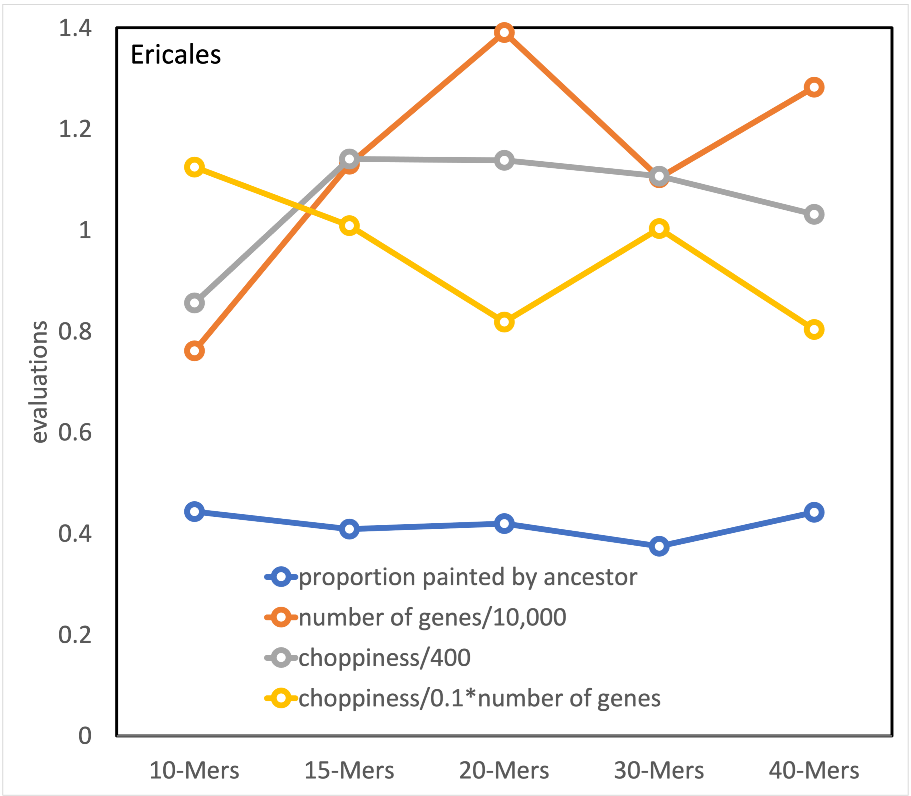

Ericales
1.4
1.2
1
0.8
0.6
0.4
0.2
0
evaluations
10-Mers
15-Mers
20-Mers
30-Mers
40-Mers
proportion painted by ancestor
number of genes/10,000
choppiness/400
choppiness/0.1*number of genes

### Supplement C. Gap statistics. For $g = 20$ each ancestor, eleven orders

Gap statistics plots also available for 15-mers, 30-mers and 40-mers.

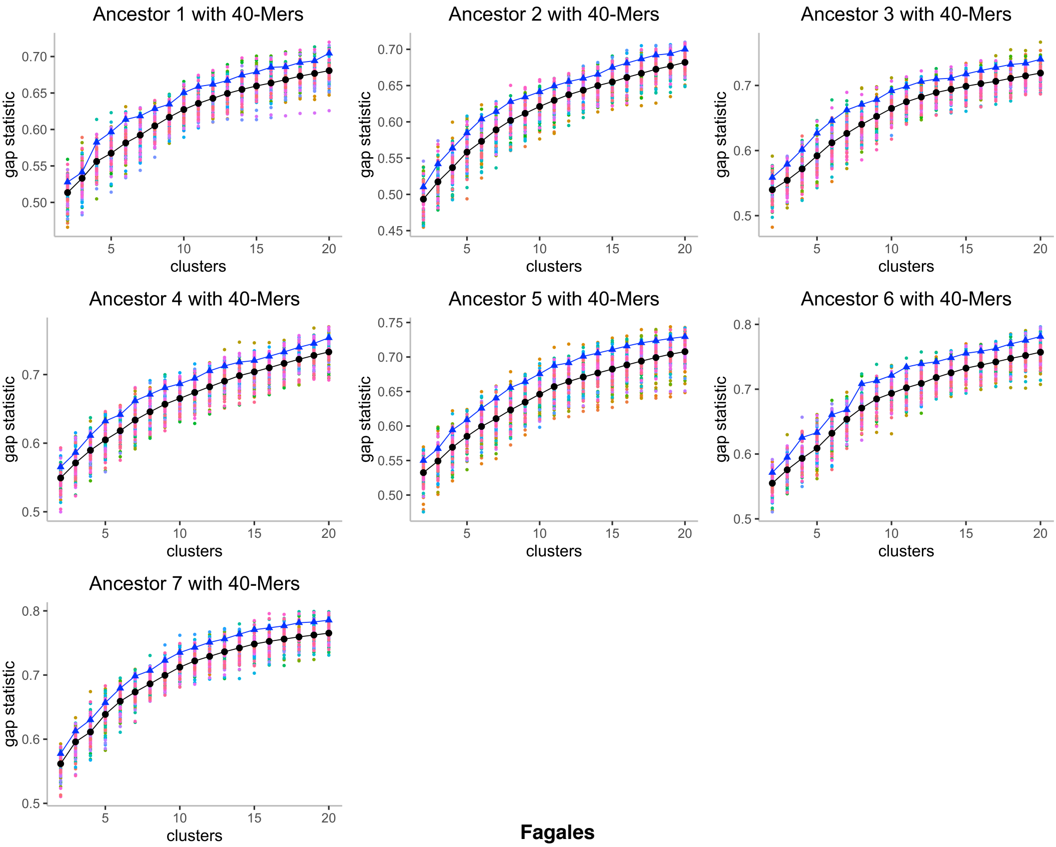
Ancestor 1 with 40-Mers
Ancestor 2 with 40-Mers
Ancestor 3 with 40-Mers
Ancestor 4 with 40-Mers
Ancestor 5 with 40-Mers
Ancestor 6 with 40-Mers
Ancestor 7 with 40-Mers
gap statistic
clusters

**Fagales**

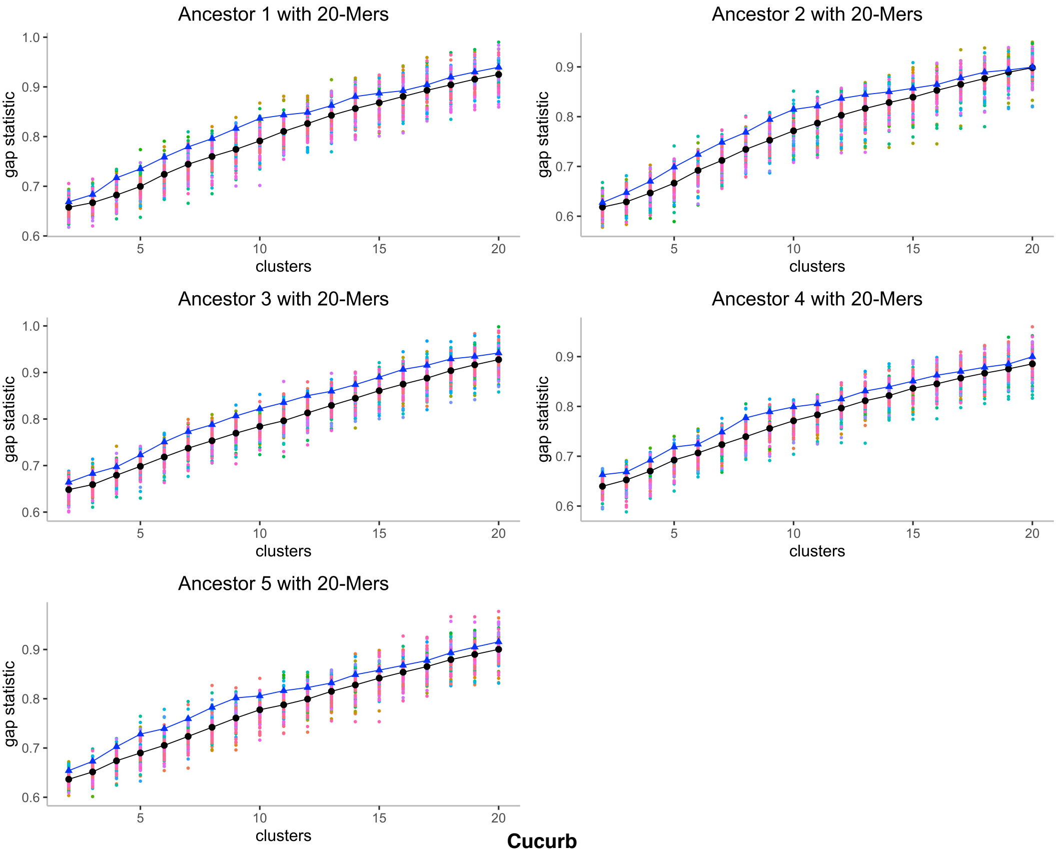
Ancestor 1 with 20-Mers
Ancestor 2 with 20-Mers
Ancestor 3 with 20-Mers
Ancestor 4 with 20-Mers
Ancestor 5 with 20-Mers
gap statistic
clusters

**Cucurb**

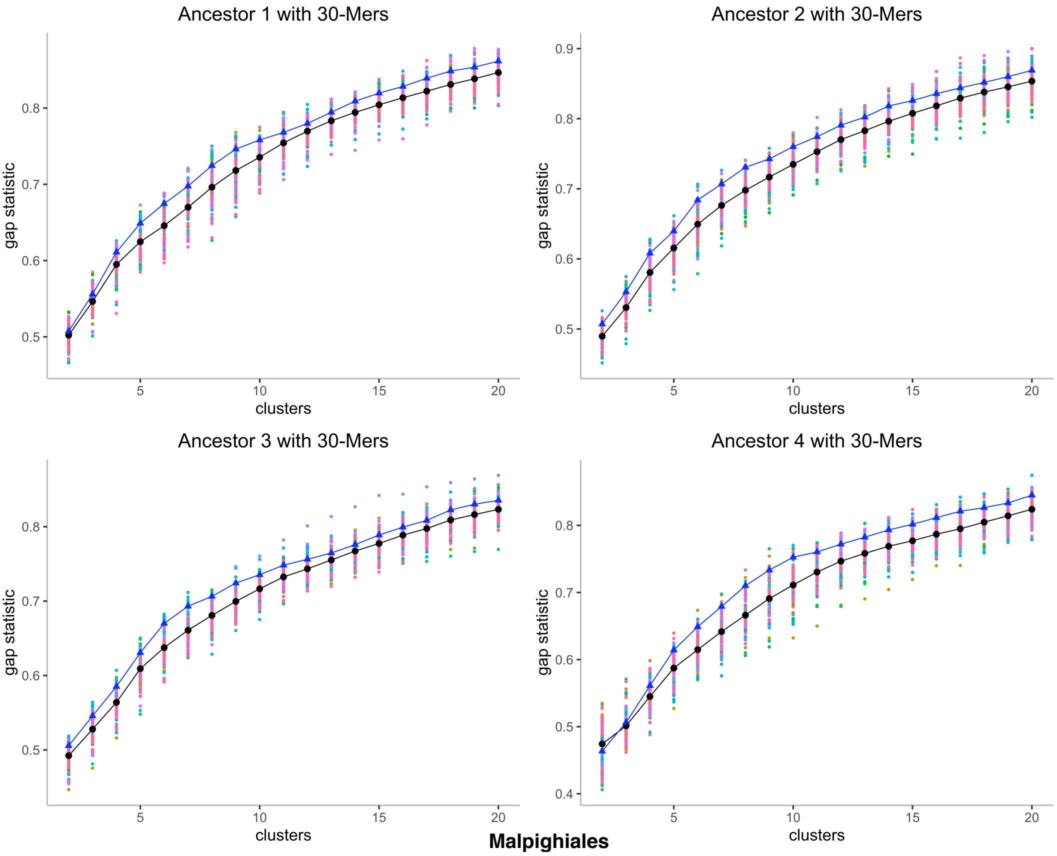
Ancestor 1 with 30-Mers
Ancestor 2 with 30-Mers
Ancestor 3 with 30-Mers
Ancestor 4 with 30-Mers
gap statistic
clusters
Malpighiales

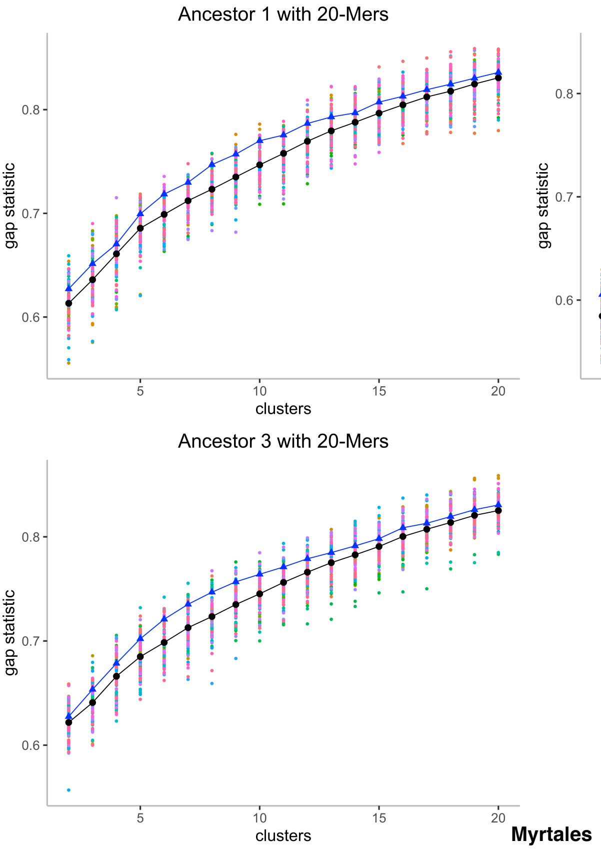
Ancestor 1 with 20-Mers
Ancestor 2 with 20-Mers
Ancestor 3 with 20-Mers
gap statistic
clusters
0.6
0.7
0.8
5
10
15
20

**Myrtales**

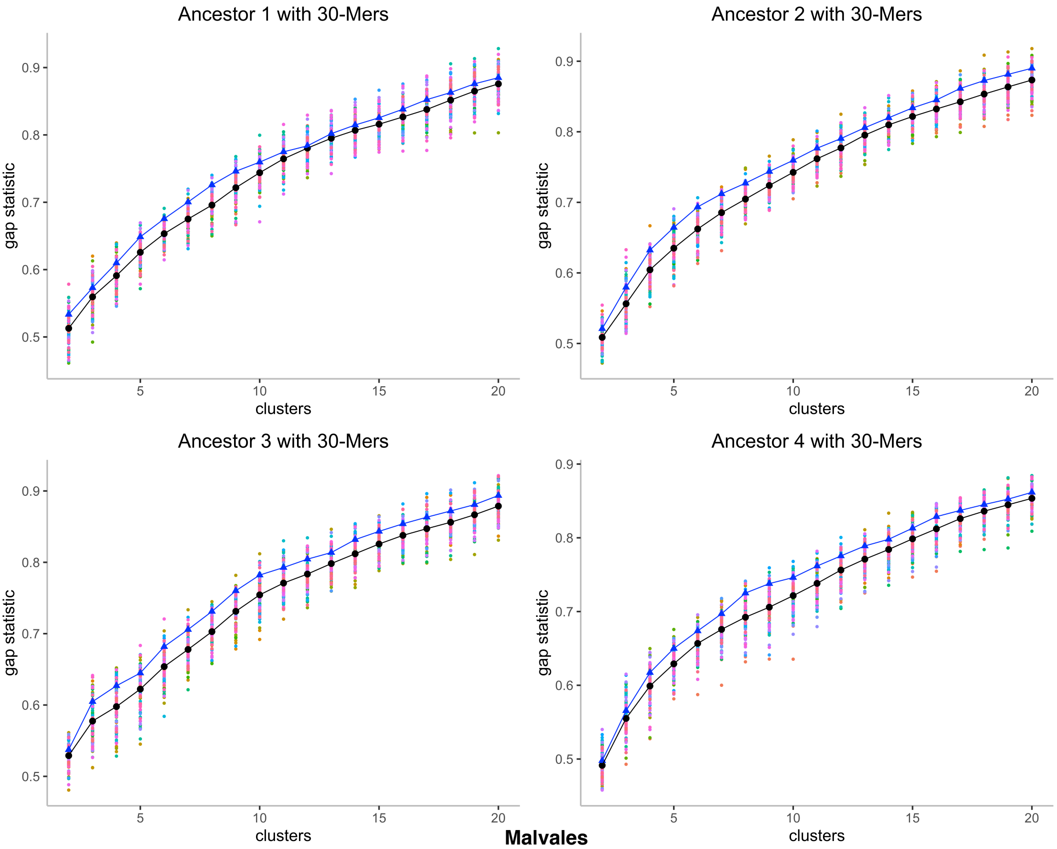
Ancestor 1 with 30-Mers
Ancestor 2 with 30-Mers
Ancestor 3 with 30-Mers
Ancestor 4 with 30-Mers
gap statistic
clusters
0.5
0.6
0.7
0.8
0.9
5
10
15
20

**Malvales**

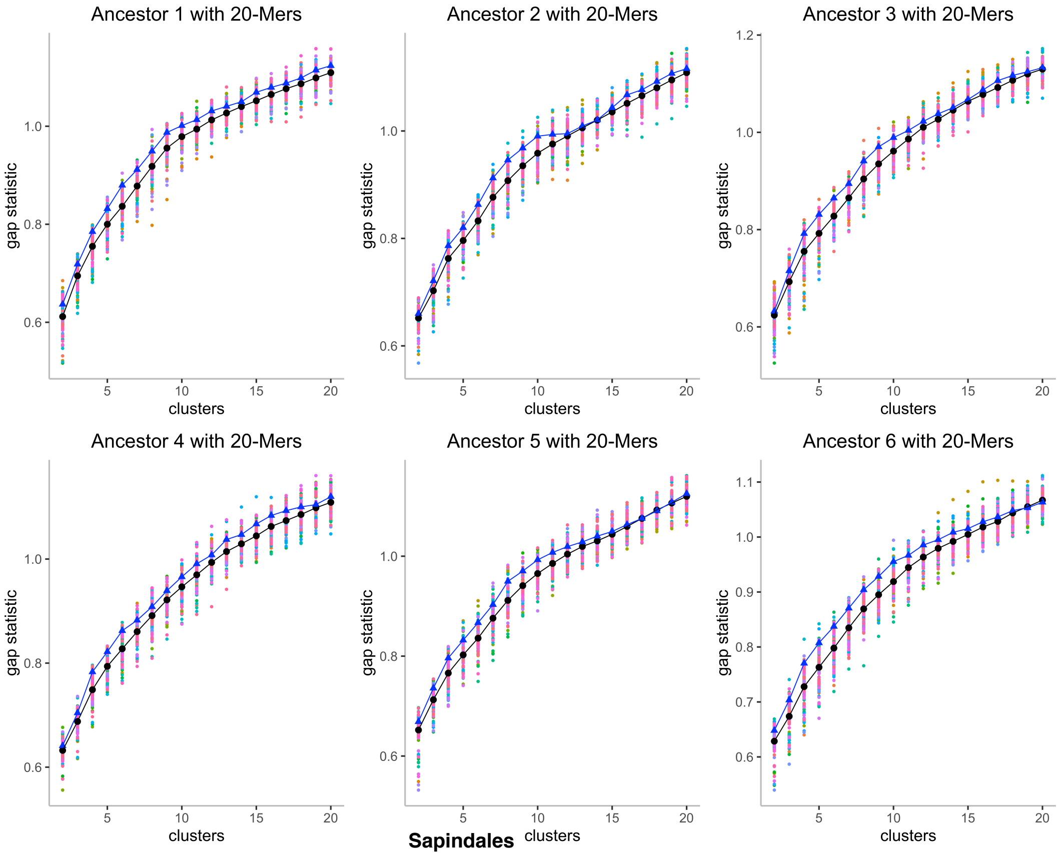

Ancestor 1 with 20-Mers
Ancestor 2 with 20-Mers
Ancestor 3 with 20-Mers
Ancestor 4 with 20-Mers
Ancestor 5 with 20-Mers
Ancestor 6 with 20-Mers
gap statistic
clusters

**Sapindales**

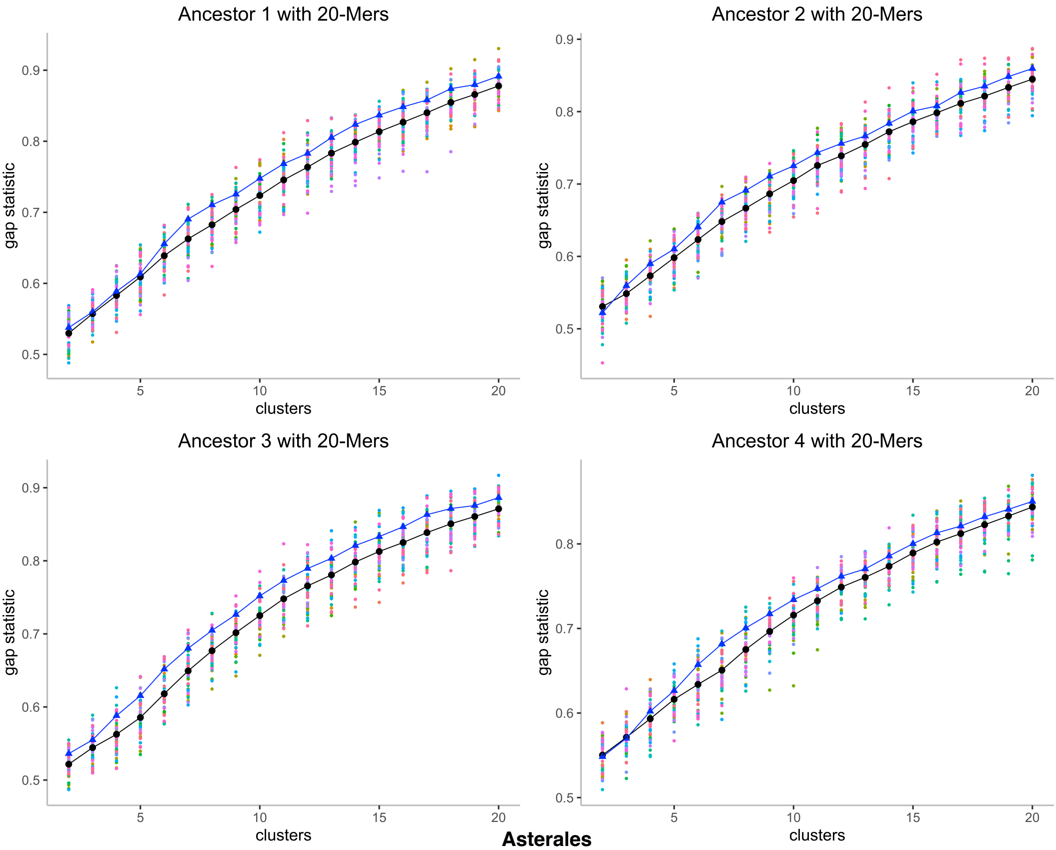
Ancestor 1 with 20-Mers
Ancestor 2 with 20-Mers
Ancestor 3 with 20-Mers
Ancestor 4 with 20-Mers
gap statistic
clusters
0.5
0.6
0.7
0.8
0.9
5
10
15
20

**Asterales**

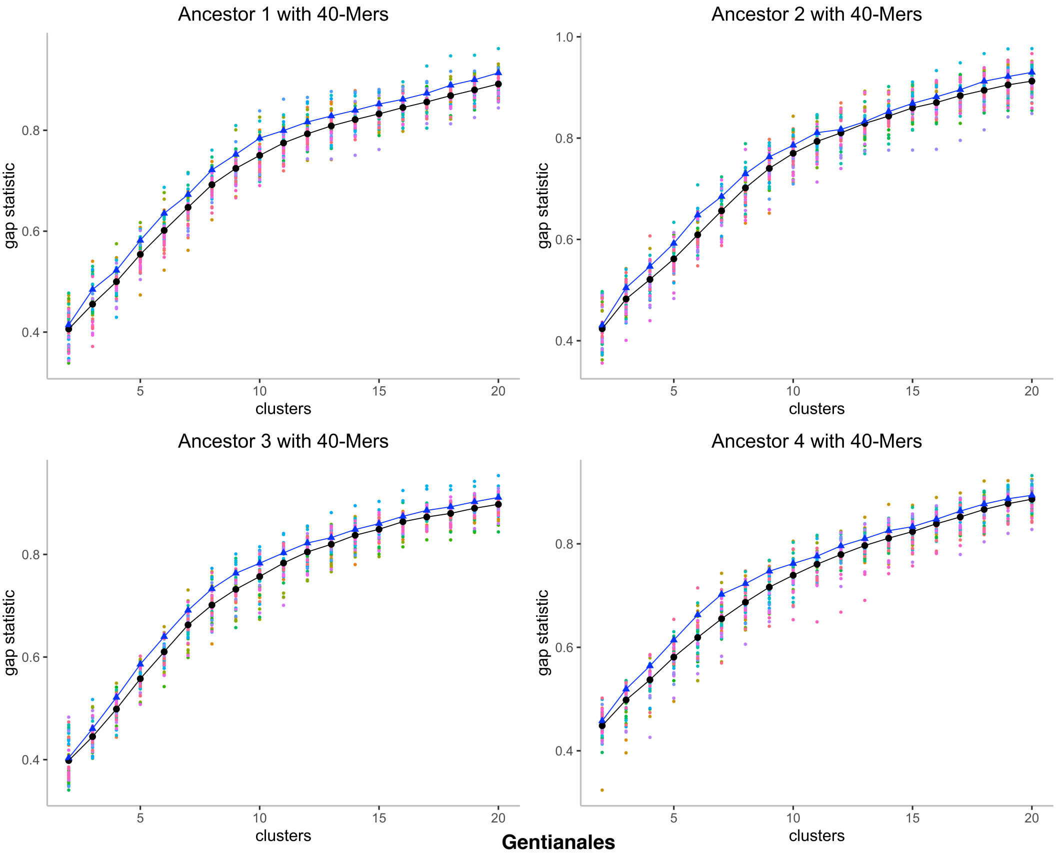

**Gentianales**

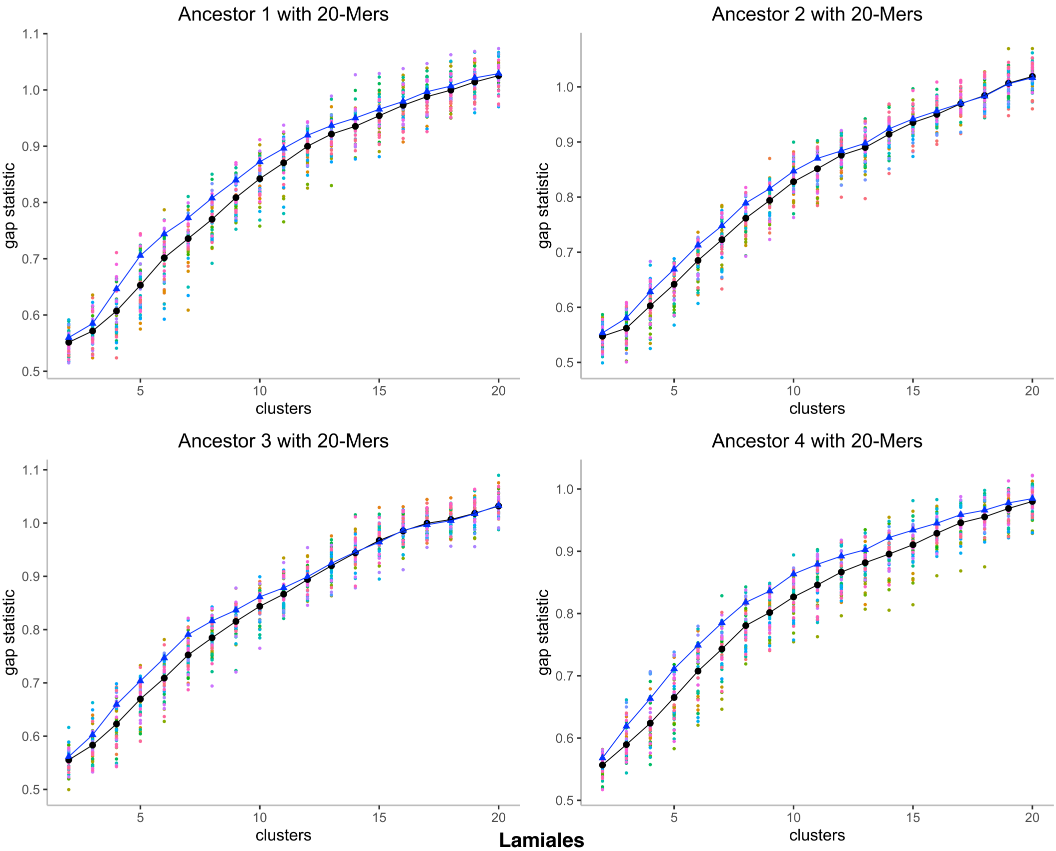
Ancestor 1 with 20-Mers
Ancestor 2 with 20-Mers
Ancestor 3 with 20-Mers
Ancestor 4 with 20-Mers
gap statistic
clusters
1.1
1.0
0.9
0.8
0.7
0.6
0.5
5
10
15
20

**Lamiales**

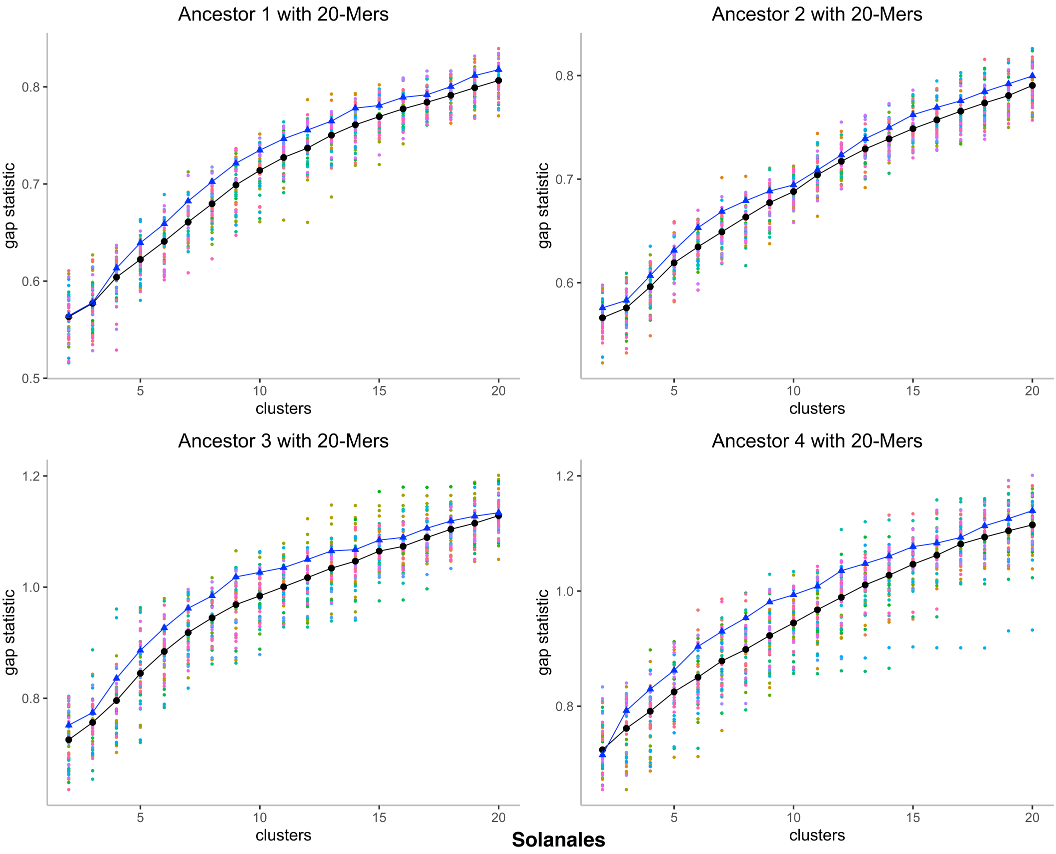
Ancestor 1 with 20-Mers
Ancestor 2 with 20-Mers
Ancestor 3 with 20-Mers
Ancestor 4 with 20-Mers
gap statistic
clusters
Solanales

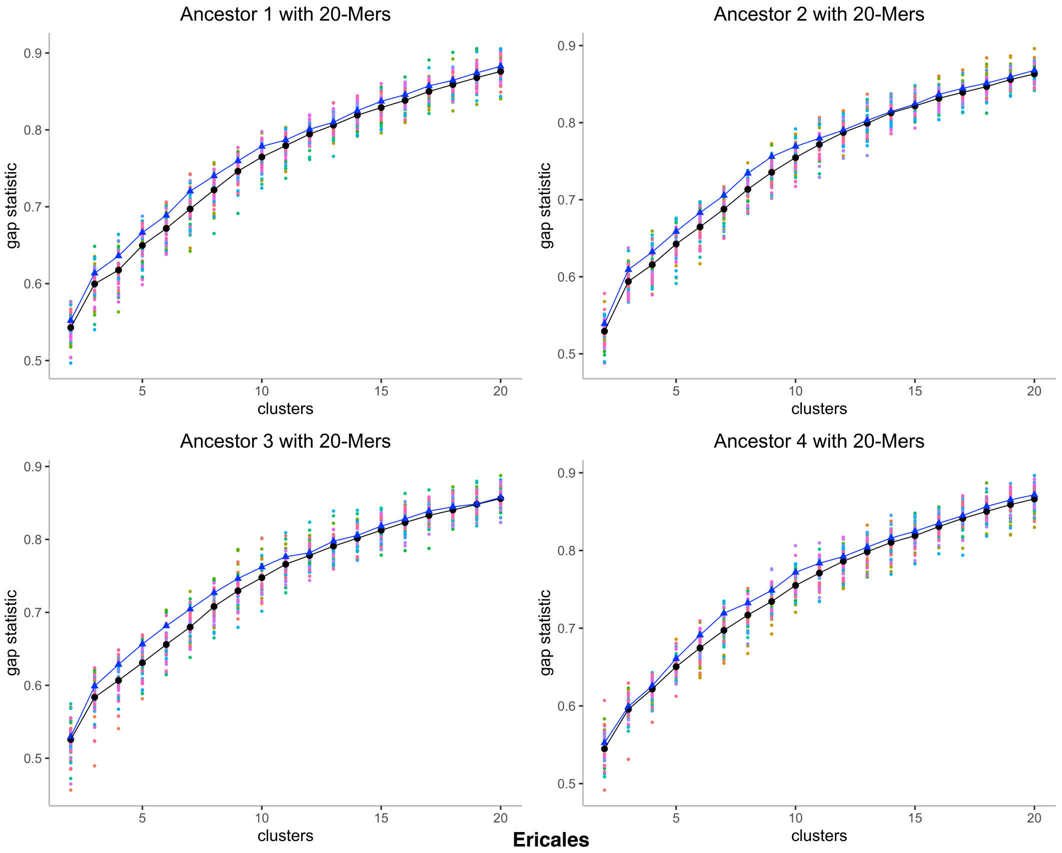
Ancestor 1 with 20-Mers
Ancestor 2 with 20-Mers
Ancestor 3 with 20-Mers
Ancestor 4 with 20-Mers
gap statistic
clusters
0.5
0.6
0.7
0.8
0.9
5
10
15
20
Ericales

### Supplement D. Painted extant genomes $g = 20$

Missing genomes where chromosome-level assemblies not available. Paintings = available for 15-mers, 20-mers, 30-mers and 40-mers.

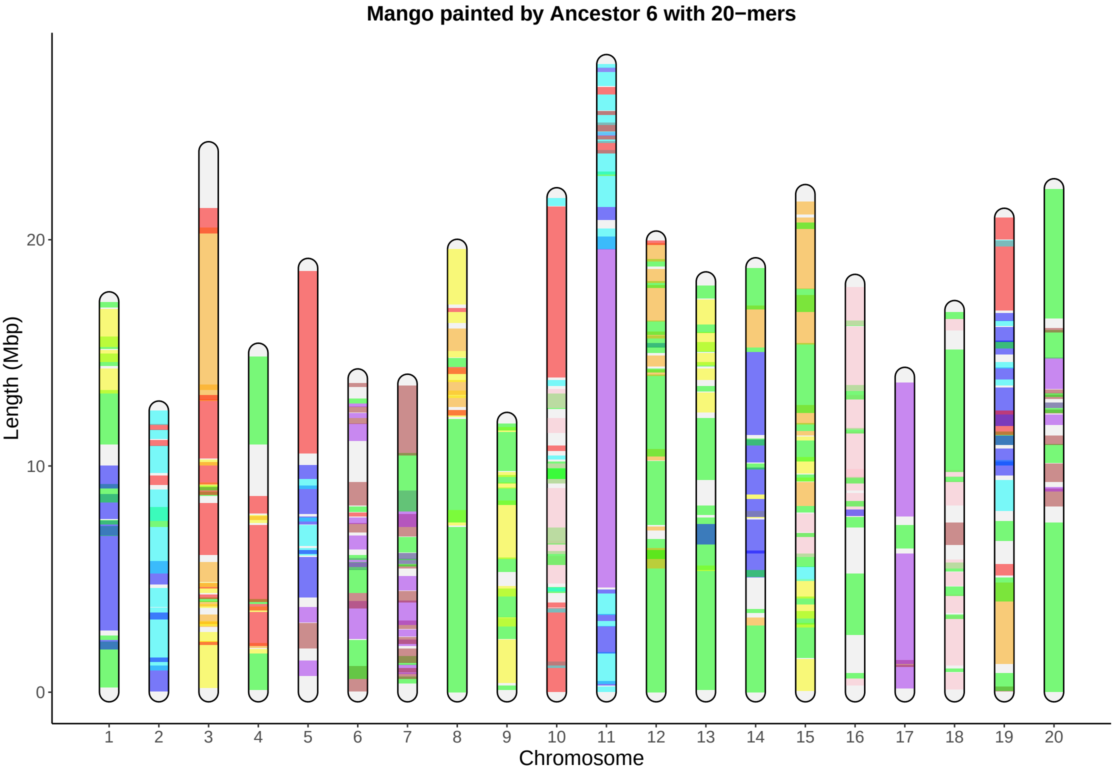
Mango painted by Ancestor 6 with 20−mers
Length (Mbp)
20
10
Chromosome

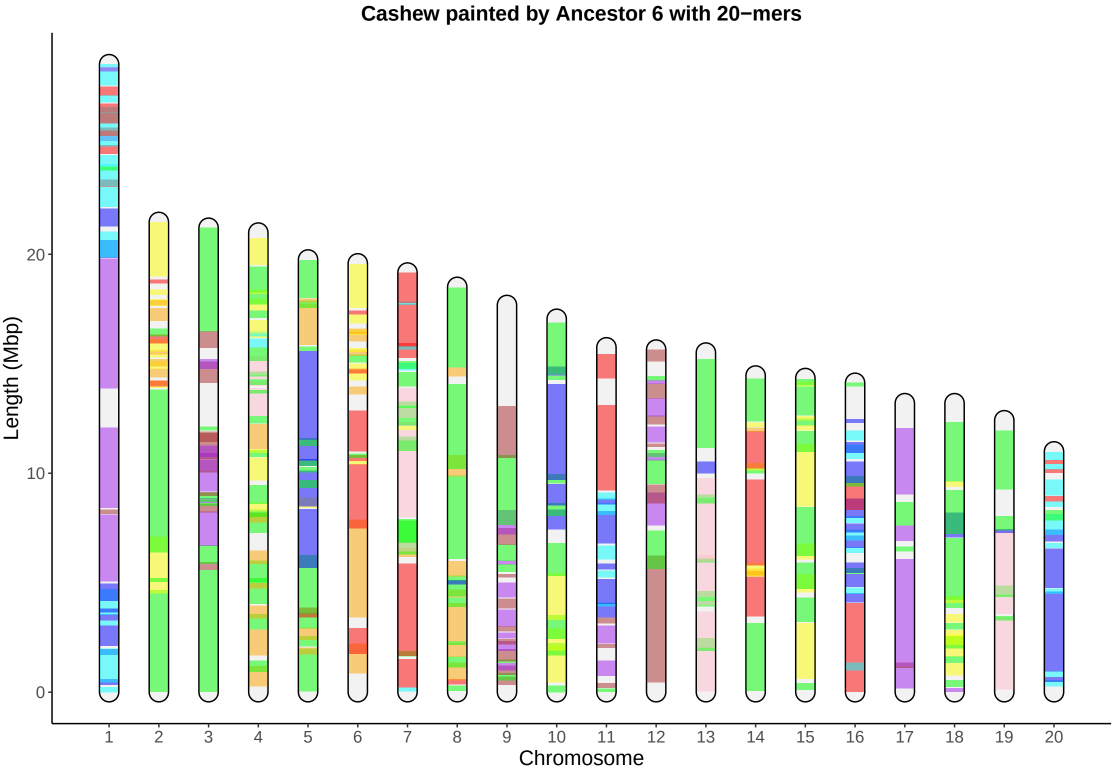
Cashew painted by Ancestor 6 with 20−mers
Length (Mbp)
0
10
20
Chromosome

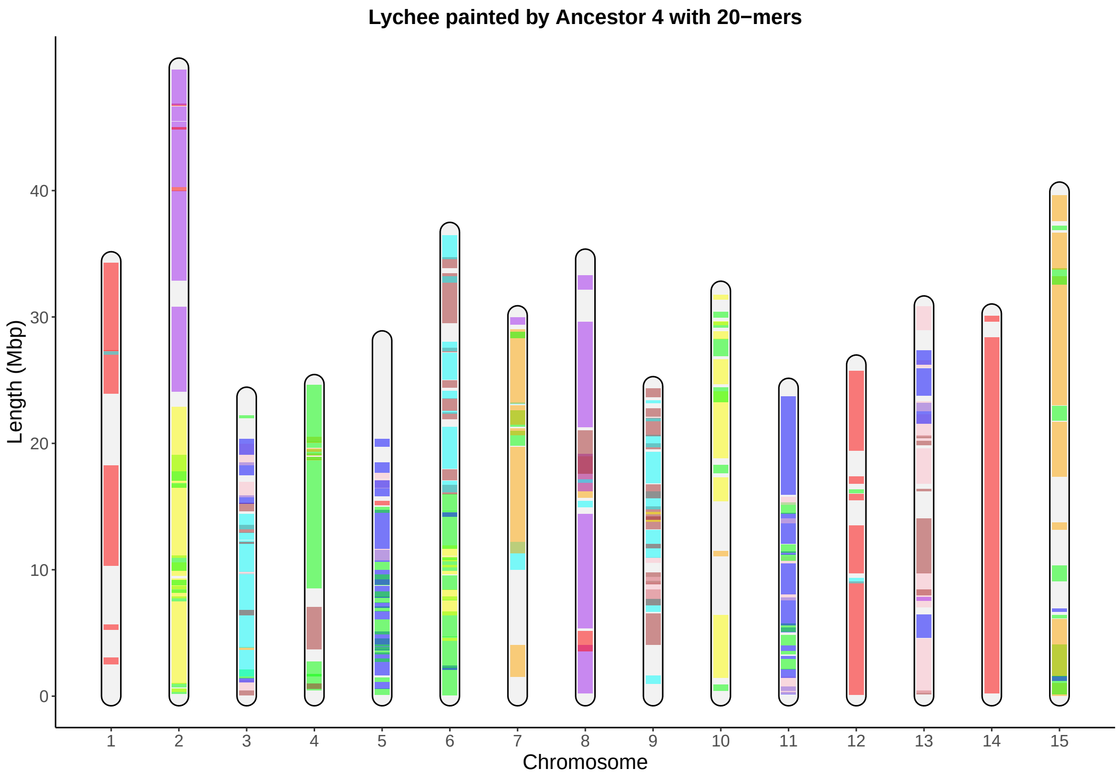
Lychee painted by Ancestor 4 with 20−mers
Length (Mbp)
0
10
20
30
40
Chromosome

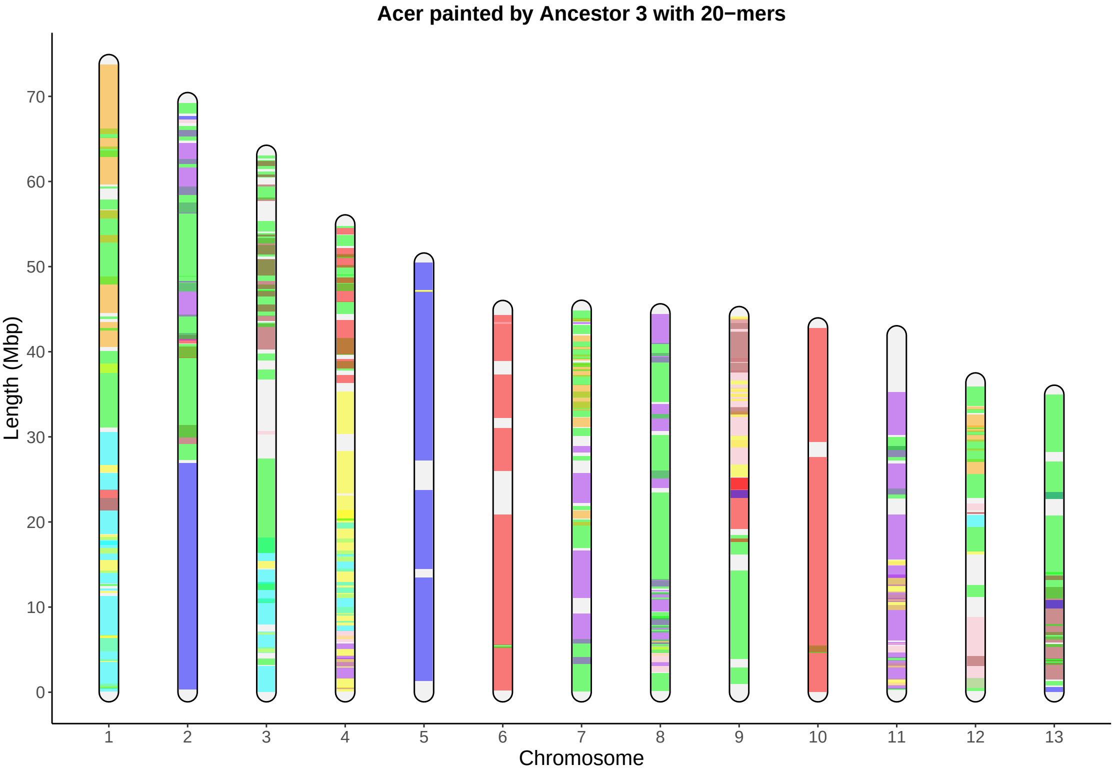

Acer painted by Ancestor 3 with 20−mers
Length (Mbp)
0
10
20
30
40
50
60
70
1
2
3
4
5
6
7
8
9
10
11
12
13
Chromosome

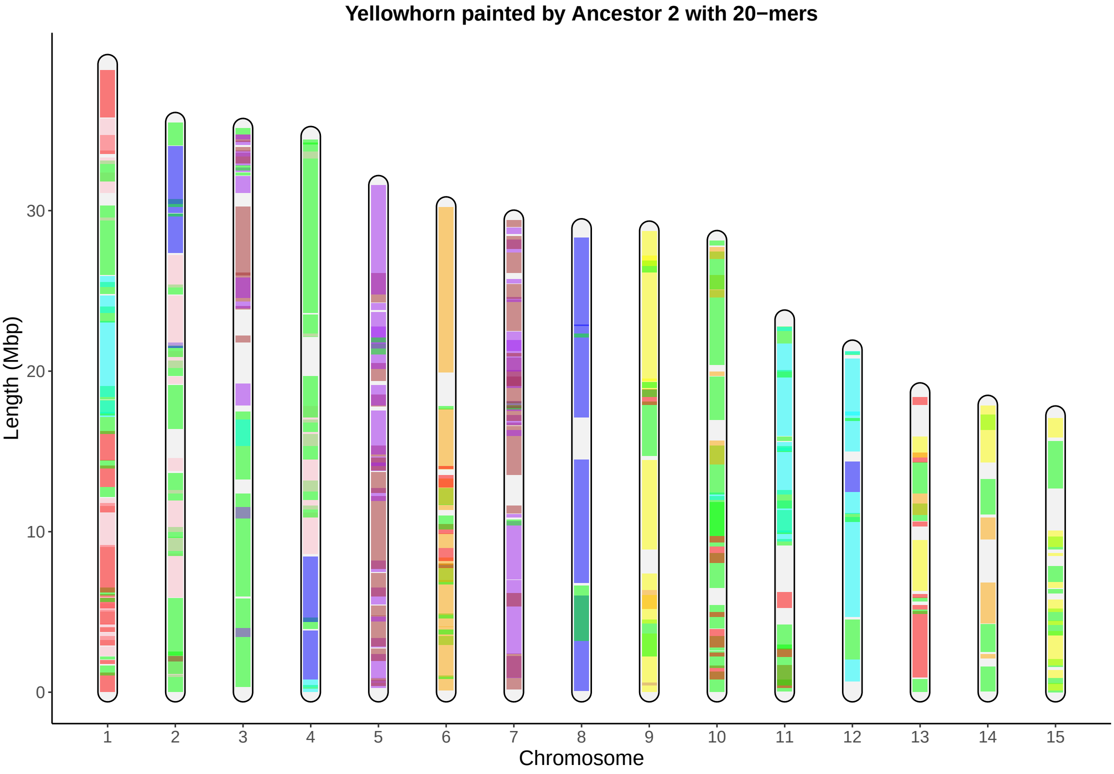

Yellowhorn painted by Ancestor 2 with 20−mers
Length (Mbp)
30
20
10
Chromosome

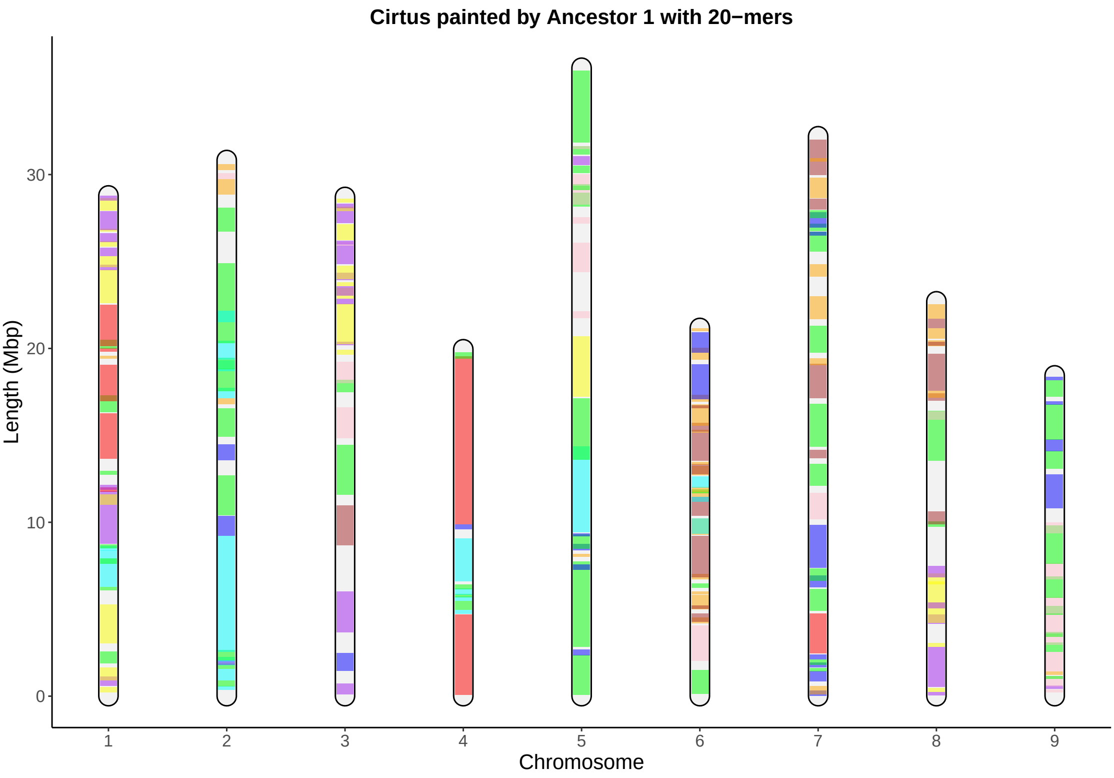

Cirtus painted by Ancestor 1 with 20−mers
Length (Mbp)
0
10
20
30
Chromosome
1
2
3
4
5
6
7
8
9

**Corymb painted by Ancestor 3 with 20−mers**

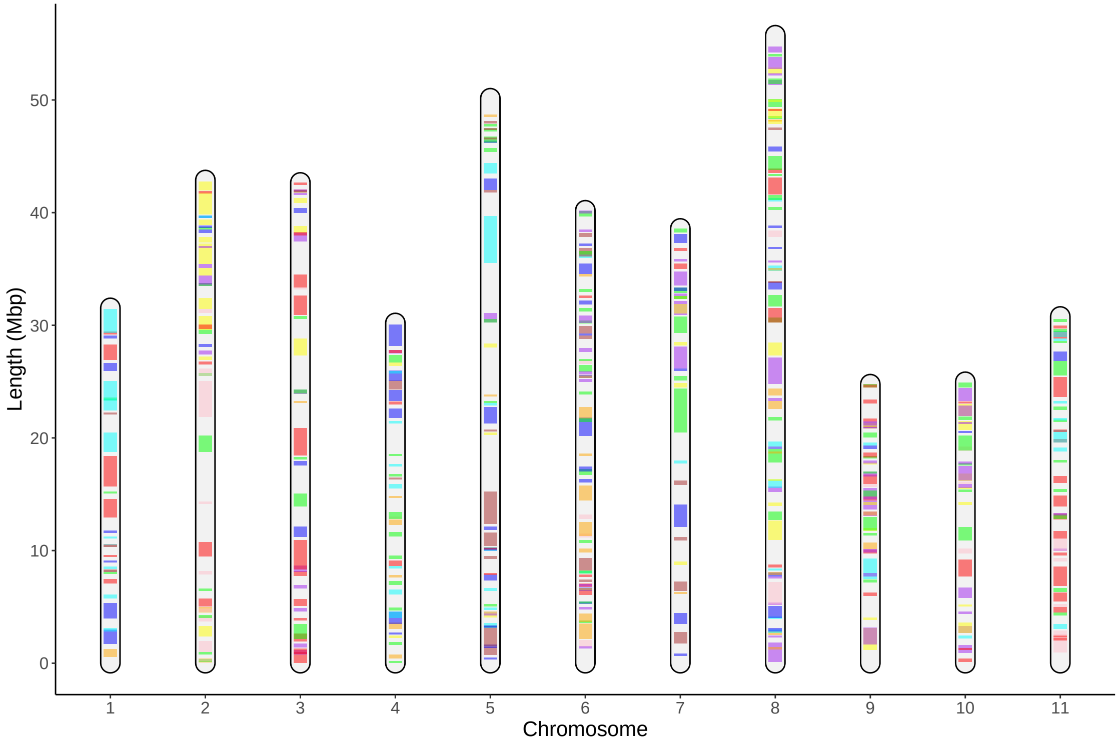

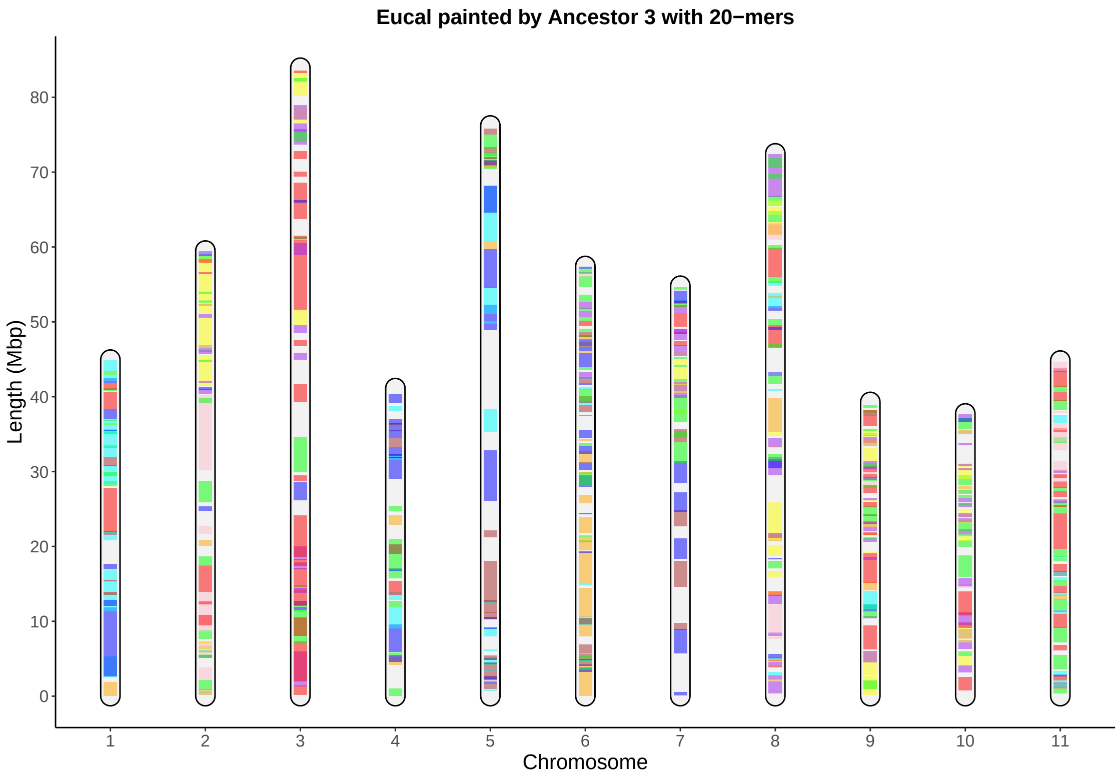

Eucal painted by Ancestor 3 with 20−mers
Length (Mbp)
0
10
20
30
40
50
60
70
80
Chromosome
1
2
3
4
5
6
7
8
9
10
11

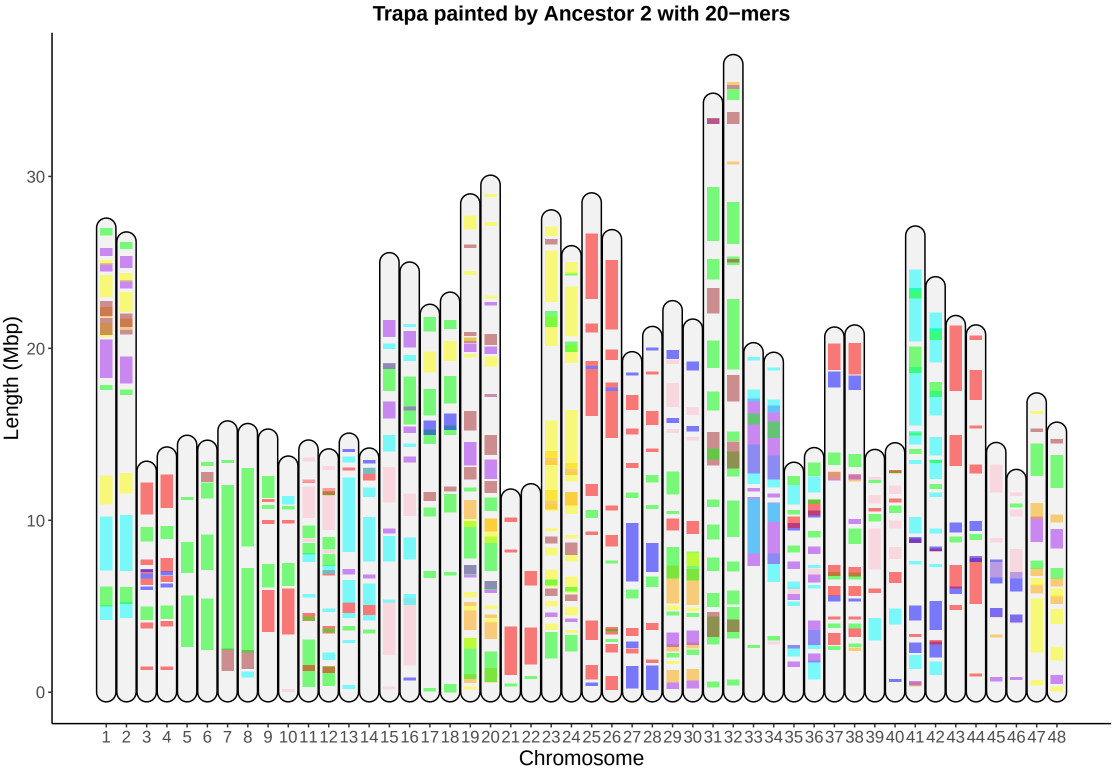

Trapa painted by Ancestor 2 with 20−mers
Length (Mbp)
0
10
20
30
1 2 3 4 5 6 7 8 9 10 11 12 13 14 15 16 17 18 19 20 21 22 23 24 25 26 27 28 29 30 31 32 33 34 35 36 37 38 39 40 41 42 43 44 45 46 47 48
Chromosome

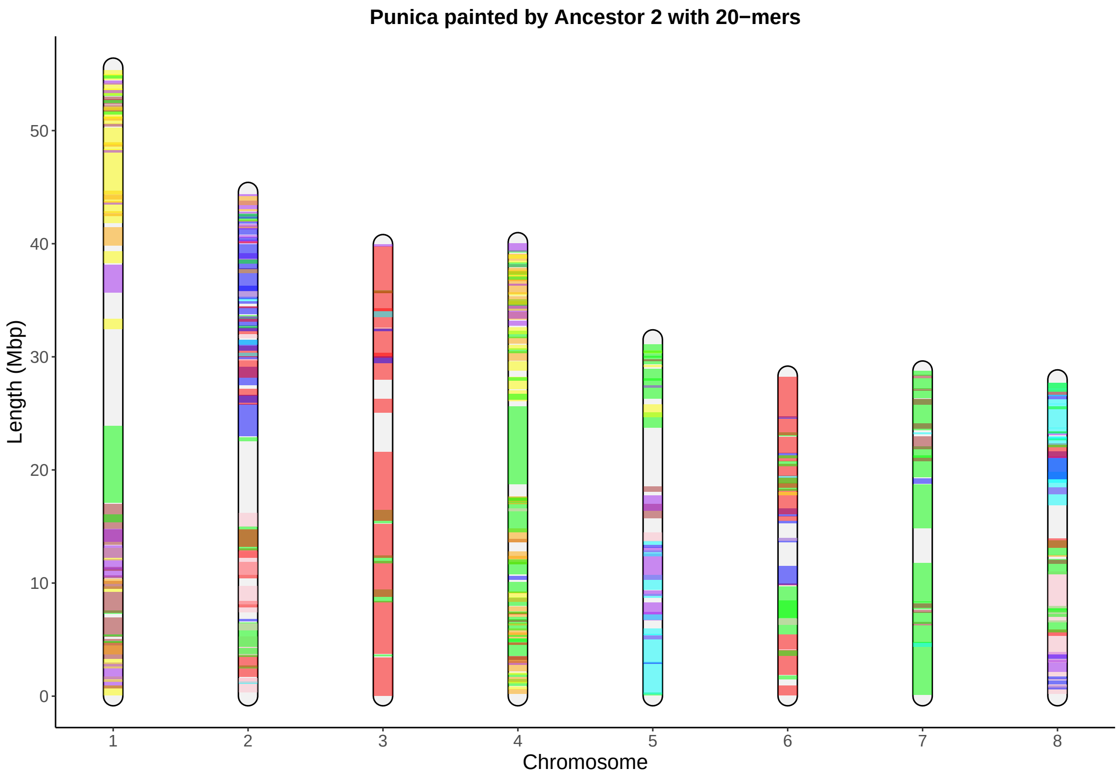

Punica painted by Ancestor 2 with 20−mers
Length (Mbp)
0
10
20
30
40
50
Chromosome
1
2
3
4
5
6
7
8

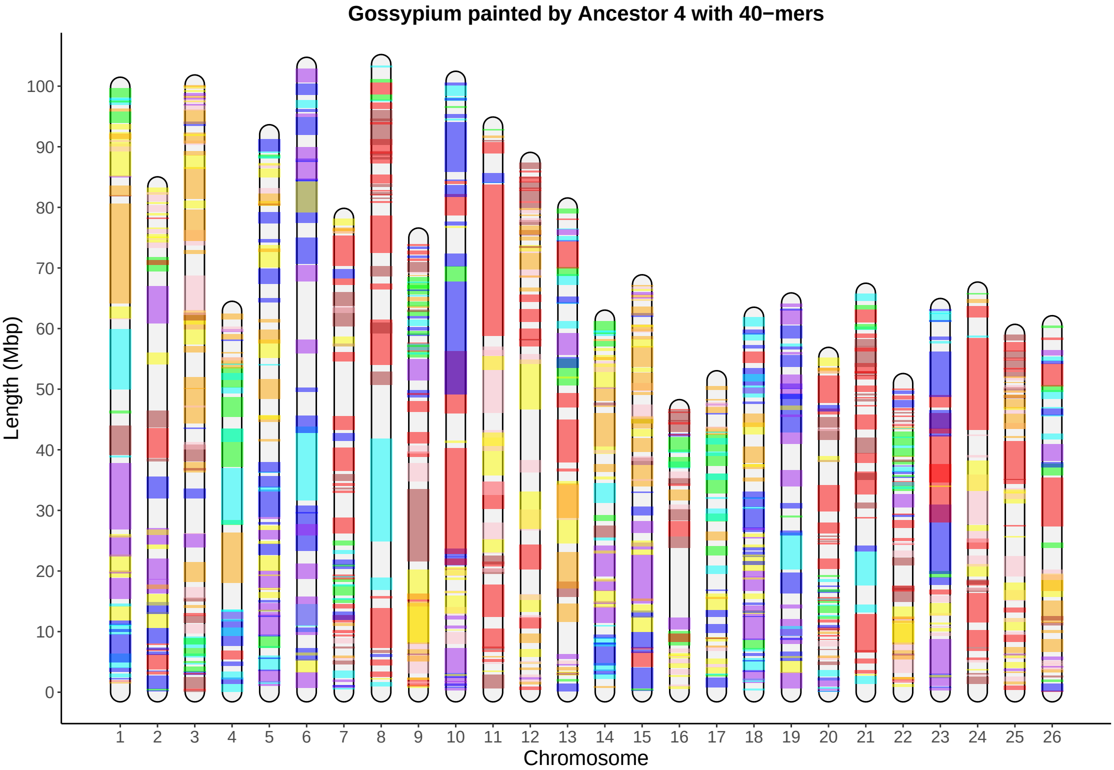
Gossypium painted by Ancestor 4 with 40−mers
Length (Mbp)
0
10
20
30
40
50
60
70
80
90
100
Chromosome

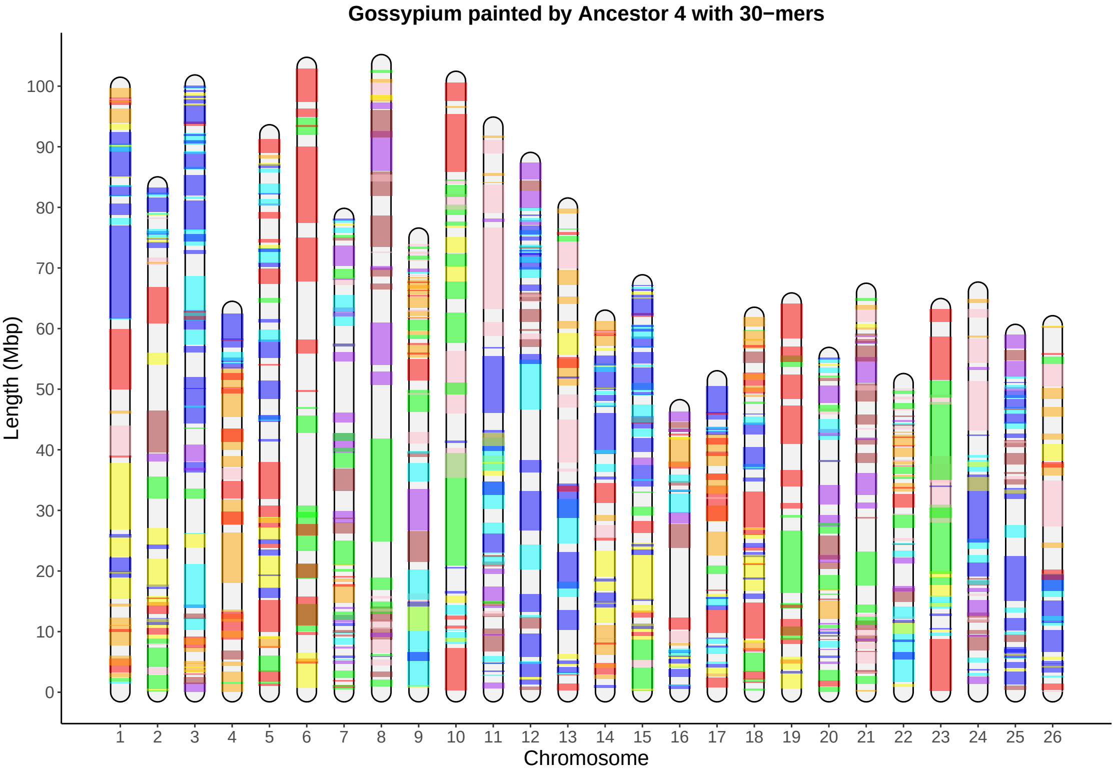

Gossypium painted by Ancestor 4 with 30−mers
Length (Mbp)
0
10
20
30
40
50
60
70
80
90
100
Chromosome
1 2 3 4 5 6 7 8 9 10 11 12 13 14 15 16 17 18 19 20 21 22 23 24 25 26

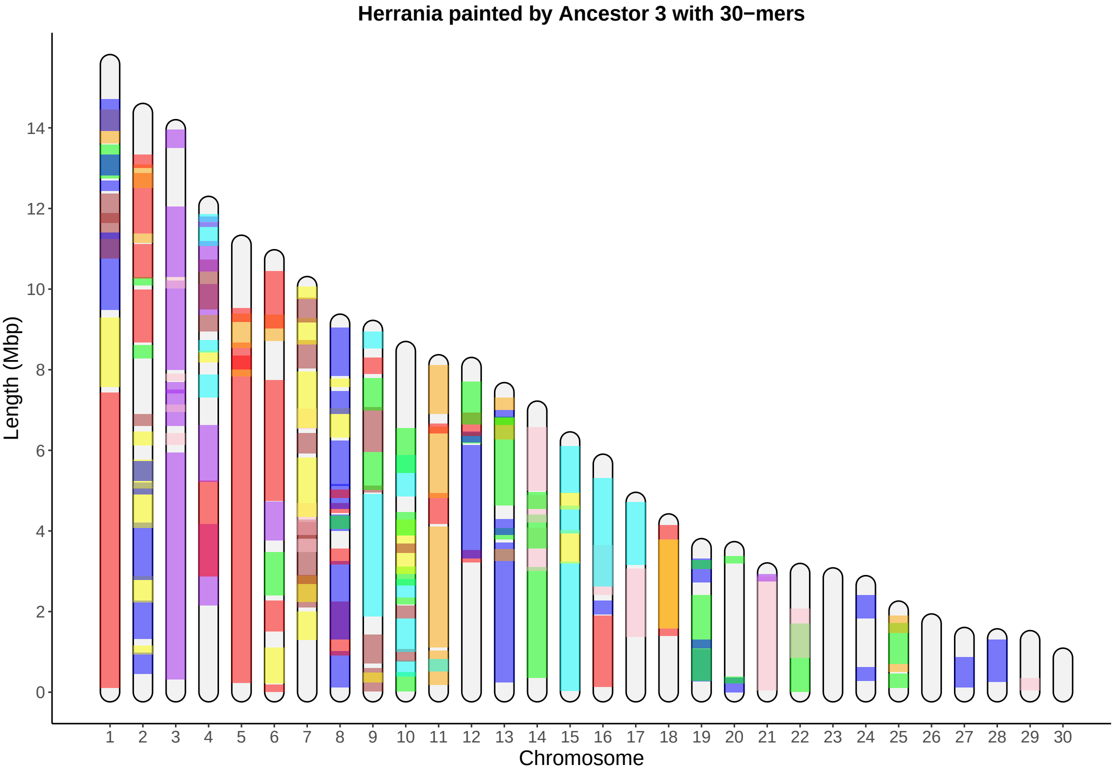
Herrania painted by Ancestor 3 with 30−mers
Length (Mbp)
0
2
4
6
8
10
12
14
Chromosome

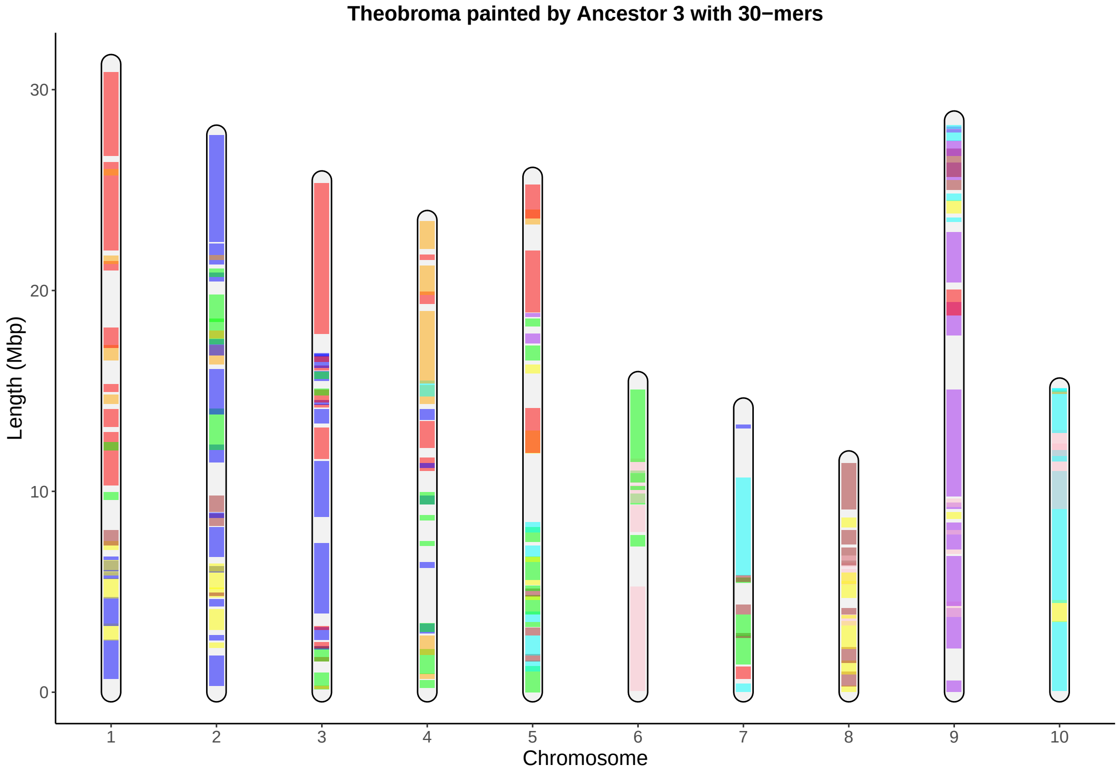
Theobroma painted by Ancestor 3 with 30−mers
Length (Mbp)
30
20
10
0
1
2
3
4
5
6
7
8
9
10
Chromosome

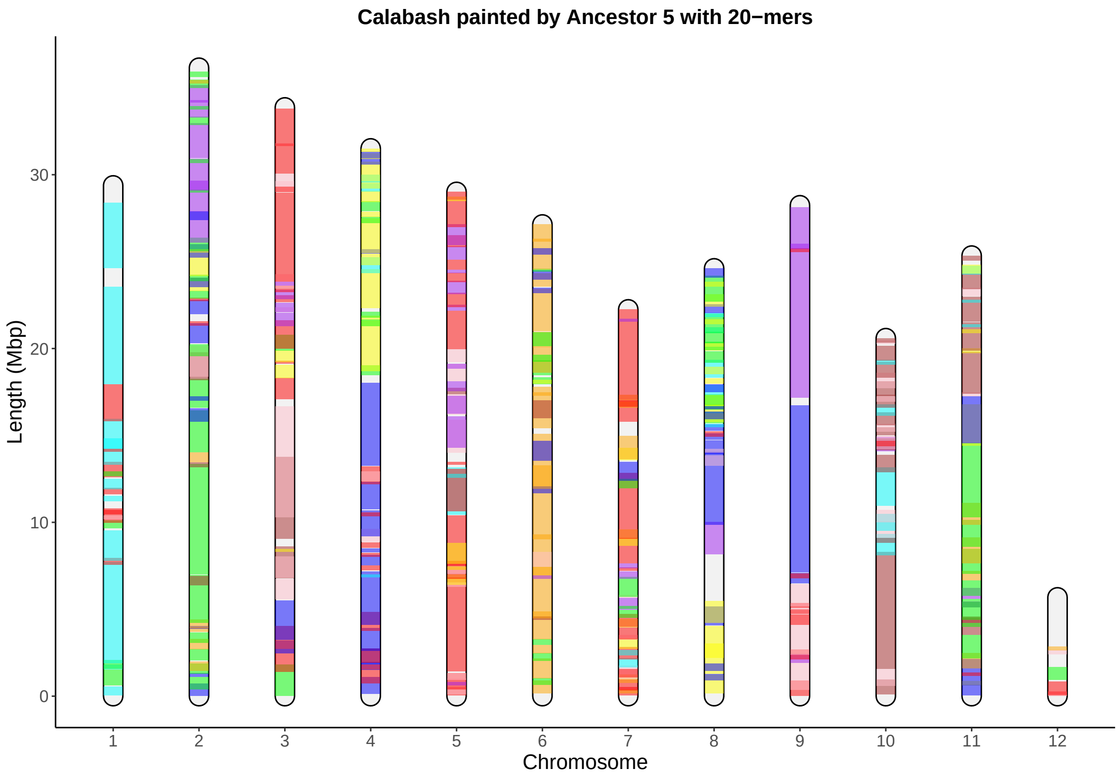
Calabash painted by Ancestor 5 with 20−mers
Length (Mbp)
0
10
20
30
1
2
3
4
5
6
7
8
9
10
11
12
Chromosome

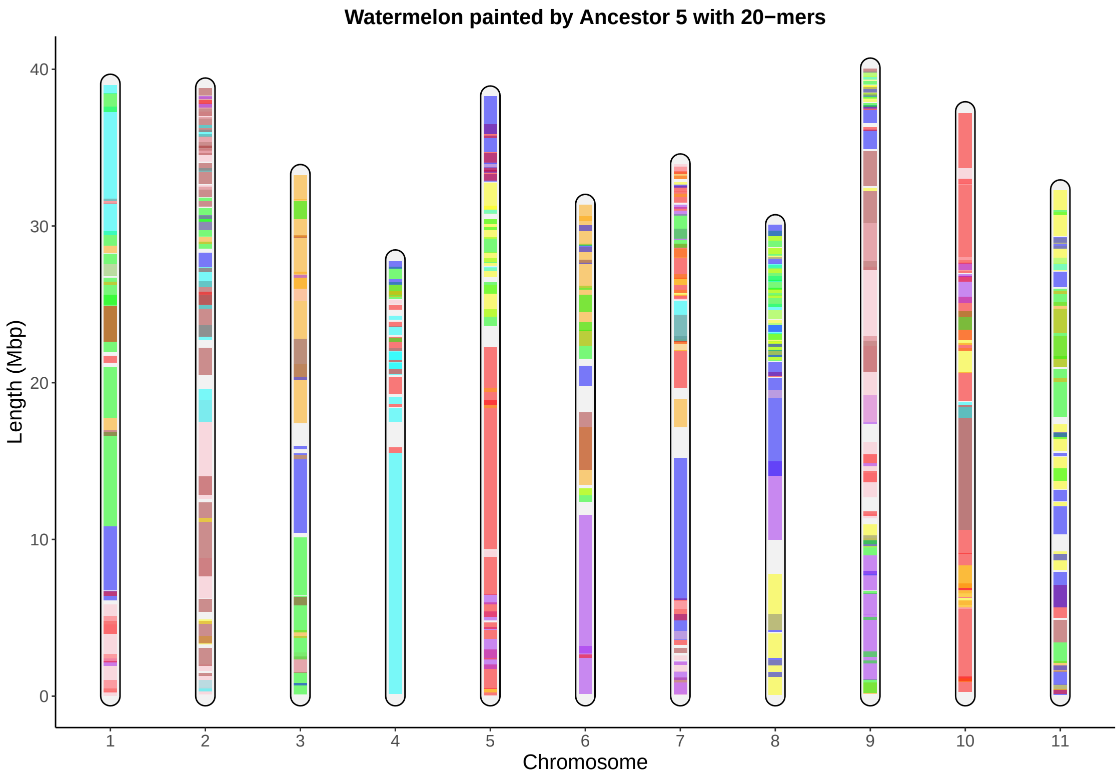
Watermelon painted by Ancestor 5 with 20−mers
Length (Mbp)
0
10
20
30
40
1
2
3
4
5
6
7
8
9
10
11
Chromosome

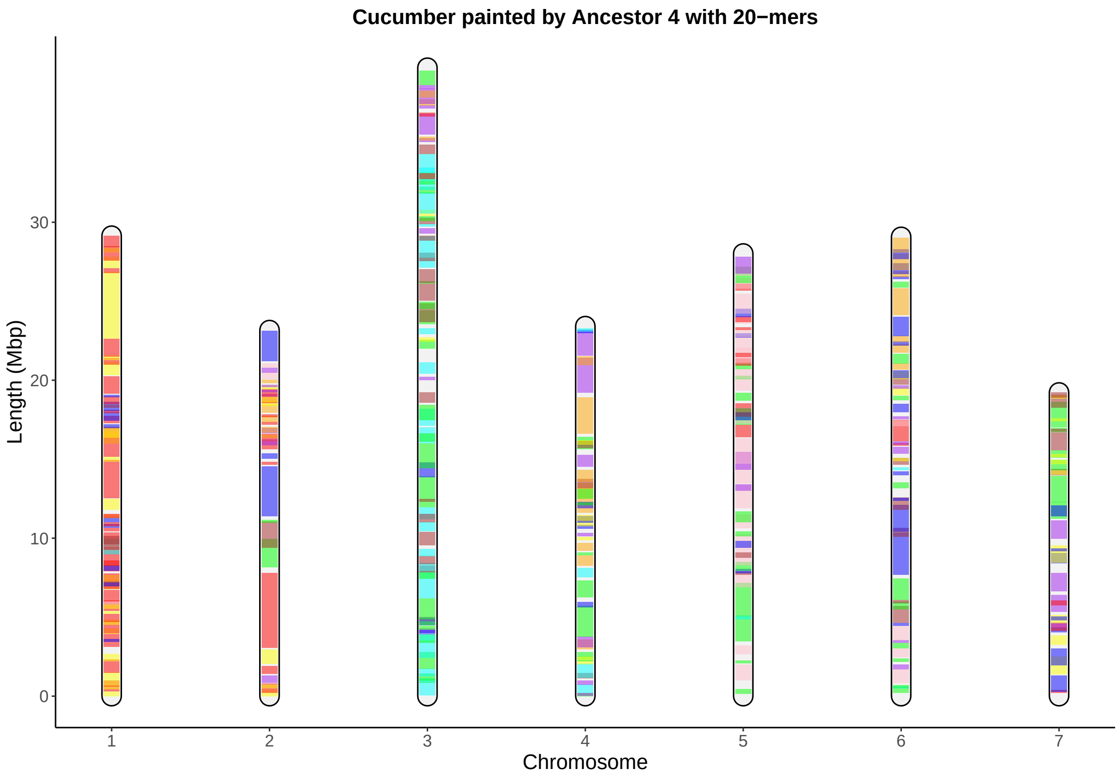

Cucumber painted by Ancestor 4 with 20−mers
Length (Mbp)
0
10
20
30
1
2
3
4
5
6
7
Chromosome

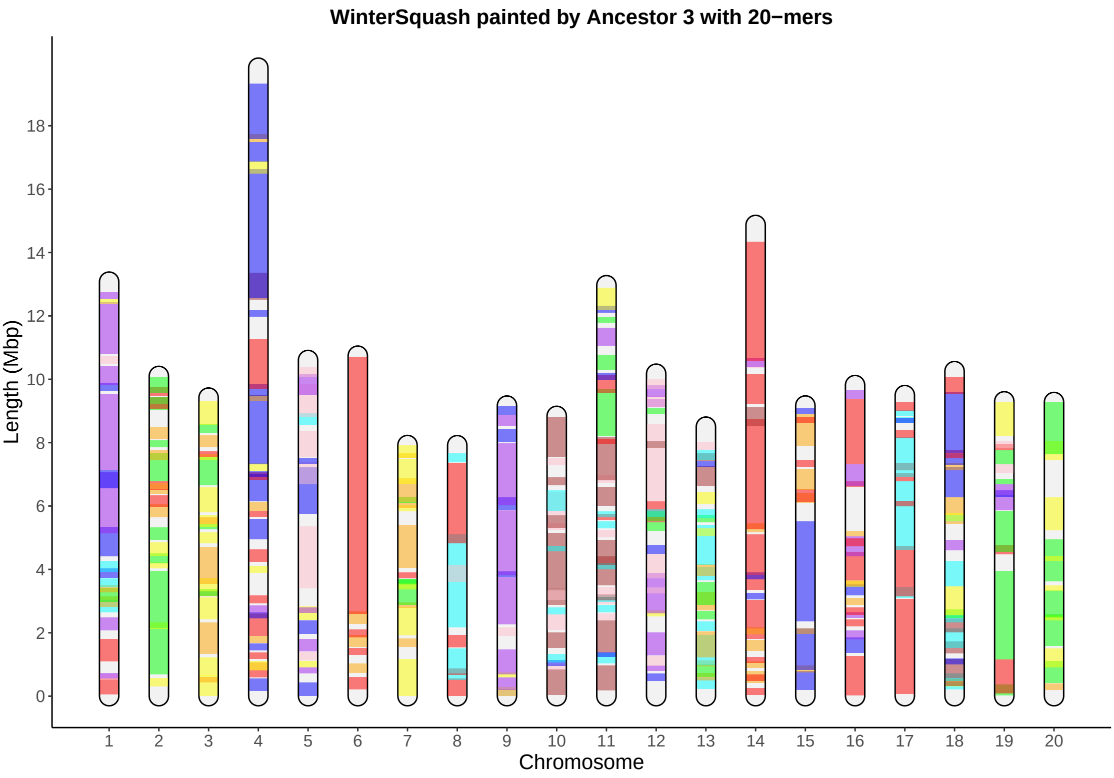
WinterSquash painted by Ancestor 3 with 20−mers
Length (Mbp)
0
2
4
6
8
10
12
14
16
18
Chromosome

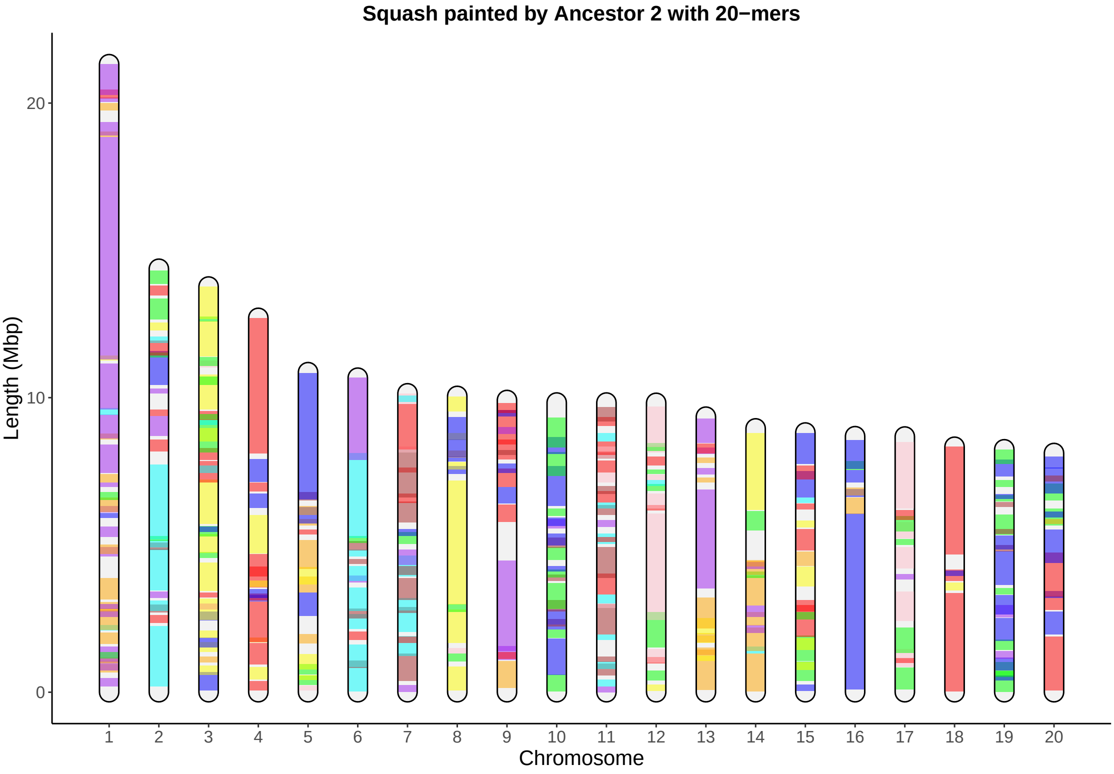
Squash painted by Ancestor 2 with 20−mers
Length (Mbp)
20
10
Chromosome

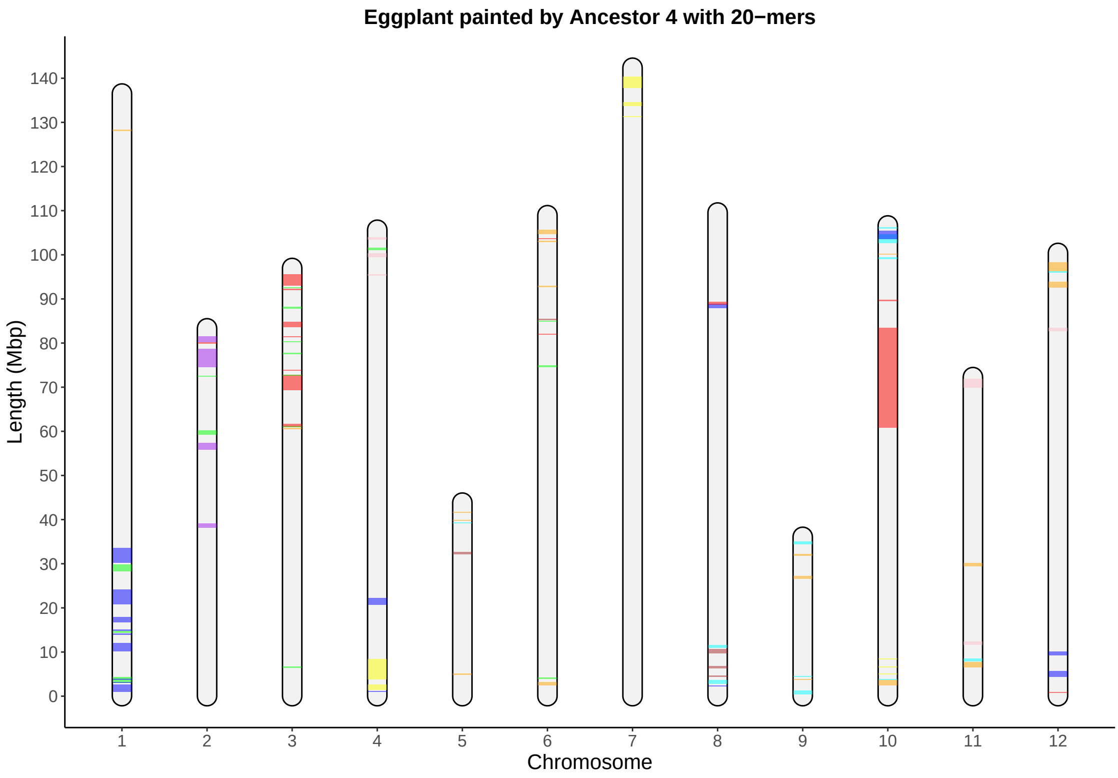

Eggplant painted by Ancestor 4 with 20−mers
Length (Mbp)
0
10
20
30
40
50
60
70
80
90
100
110
120
130
140
Chromosome
1
2
3
4
5
6
7
8
9
10
11
12

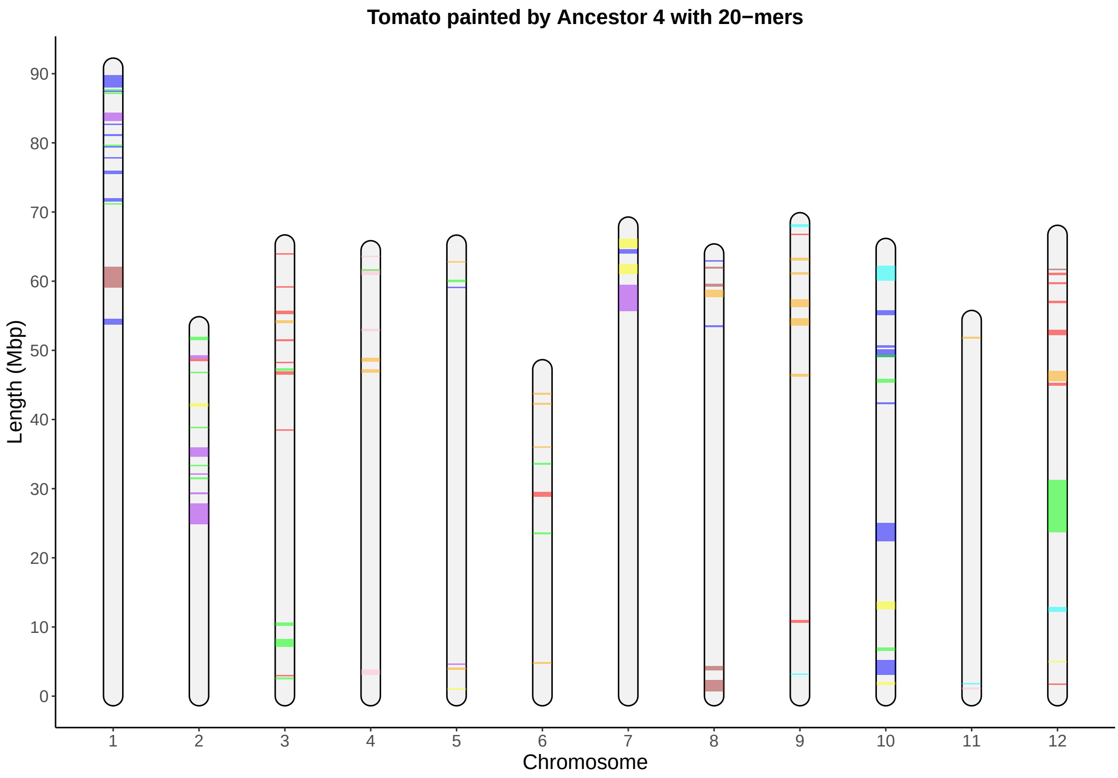

Tomato painted by Ancestor 4 with 20−mers
Length (Mbp)
0
10
20
30
40
50
60
70
80
90
Chromosome
1
2
3
4
5
6
7
8
9
10
11
12

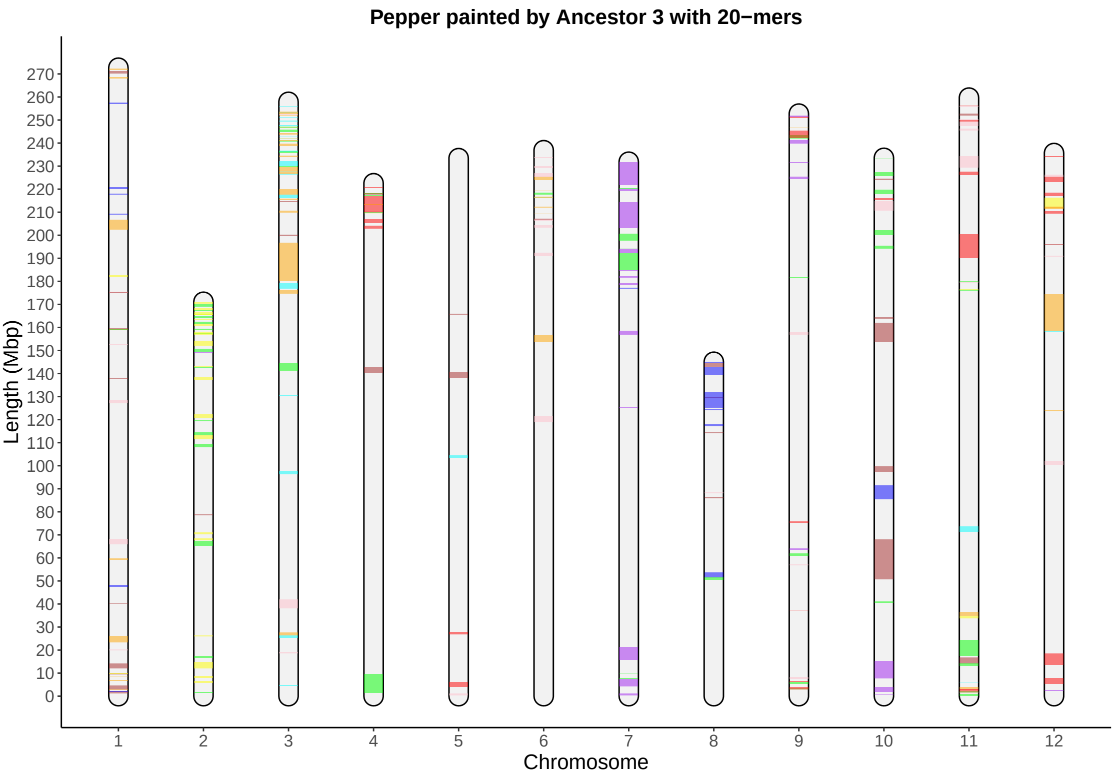

Pepper painted by Ancestor 3 with 20−mers
Length (Mbp)
270
260
250
240
230
220
210
200
190
180
170
160
150
140
130
120
110
100
90
80
70
60
50
40
30
20
10
0
1
2
3
4
5
6
7
8
9
10
11
12
Chromosome

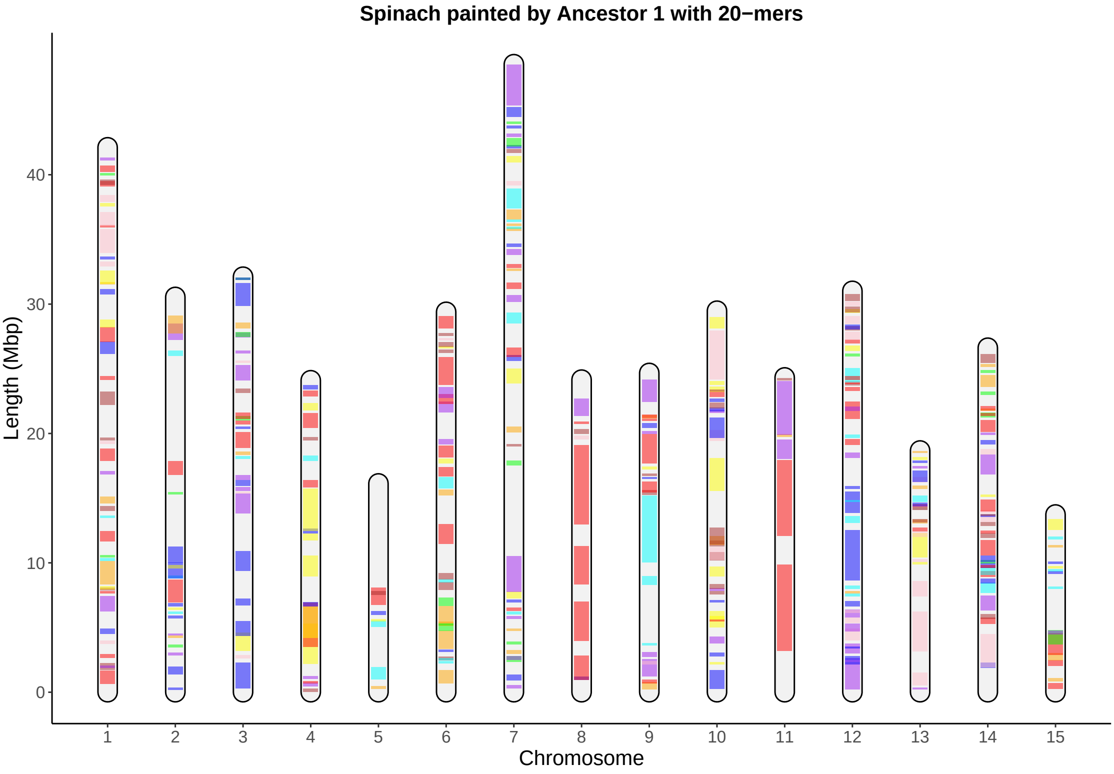

Spinach painted by Ancestor 1 with 20−mers
Length (Mbp)
0
10
20
30
40
Chromosome

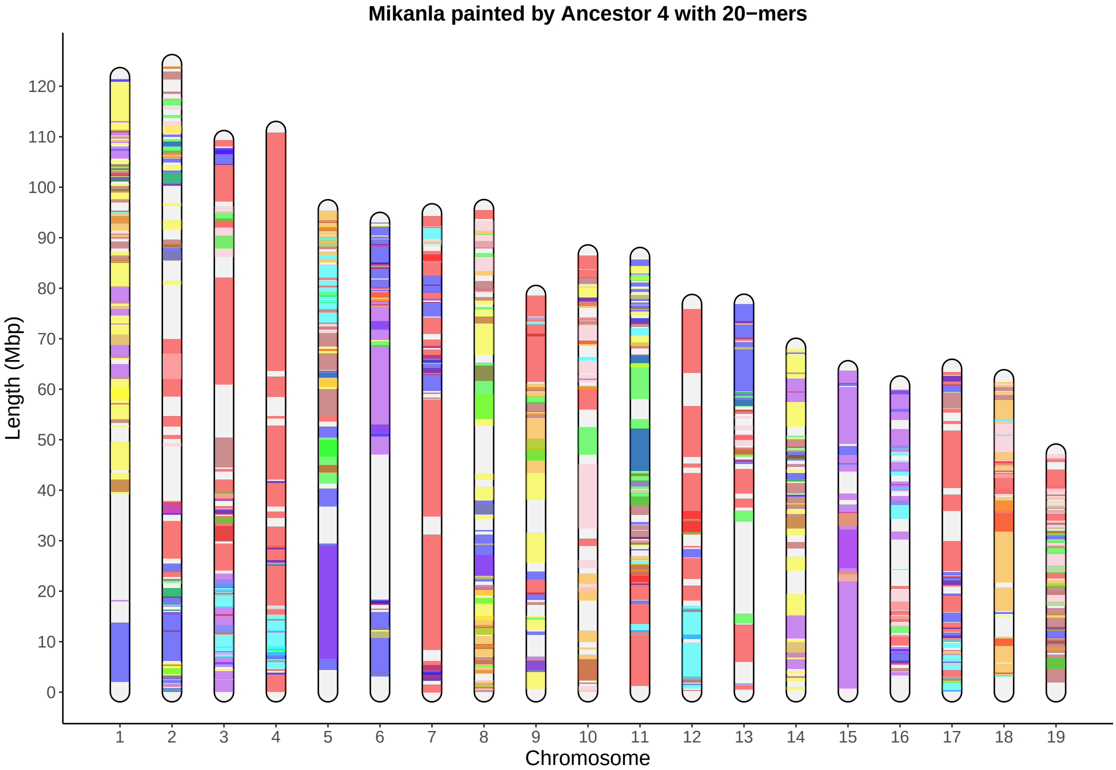

Mikanla painted by Ancestor 4 with 20−mers
Length (Mbp)
0
10
20
30
40
50
60
70
80
90
100
110
120
Chromosome

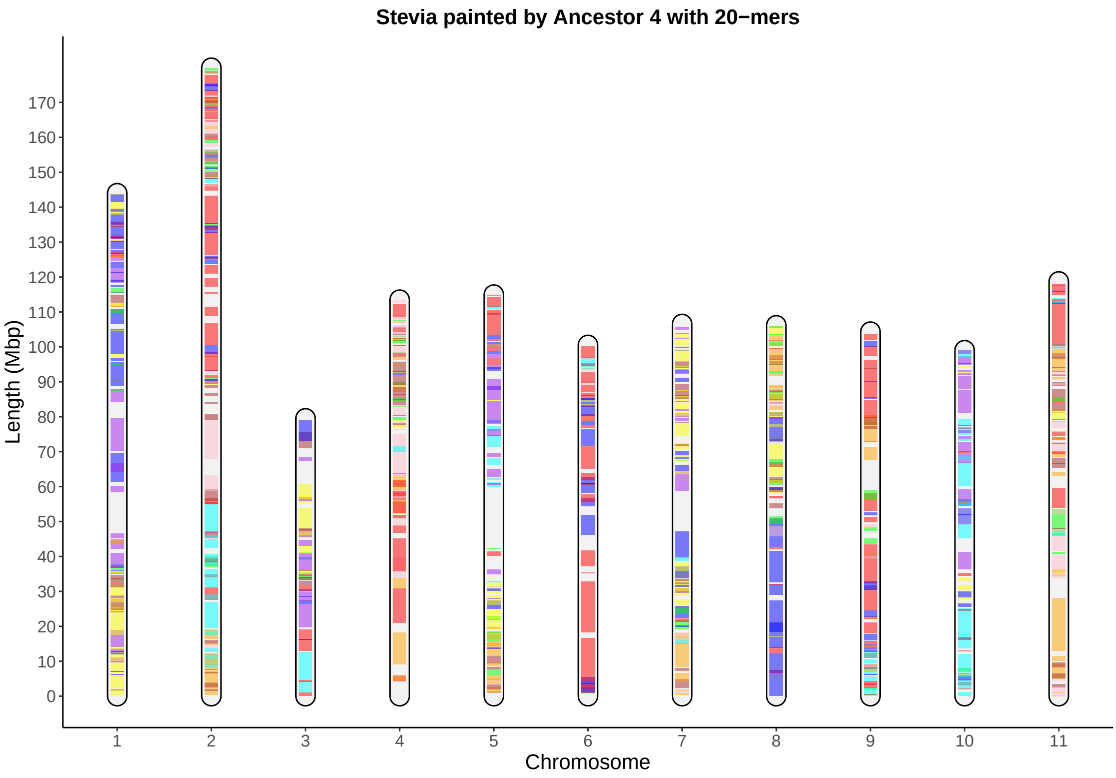

Stevia painted by Ancestor 4 with 20−mers
Length (Mbp)
0
10
20
30
40
50
60
70
80
90
100
110
120
130
140
150
160
170
Chromosome
1
2
3
4
5
6
7
8
9
10
11

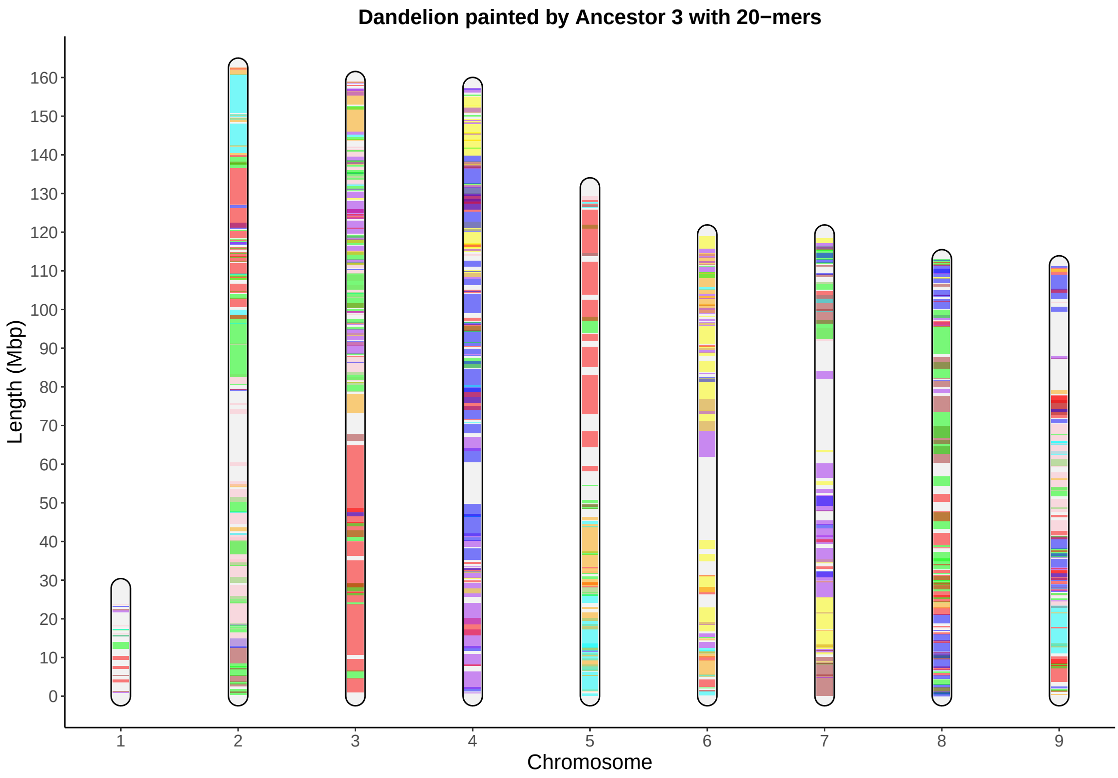
Dandelion painted by Ancestor 3 with 20−mers
Length (Mbp)
0
10
20
30
40
50
60
70
80
90
100
110
120
130
140
150
160
Chromosome
1
2
3
4
5
6
7
8
9

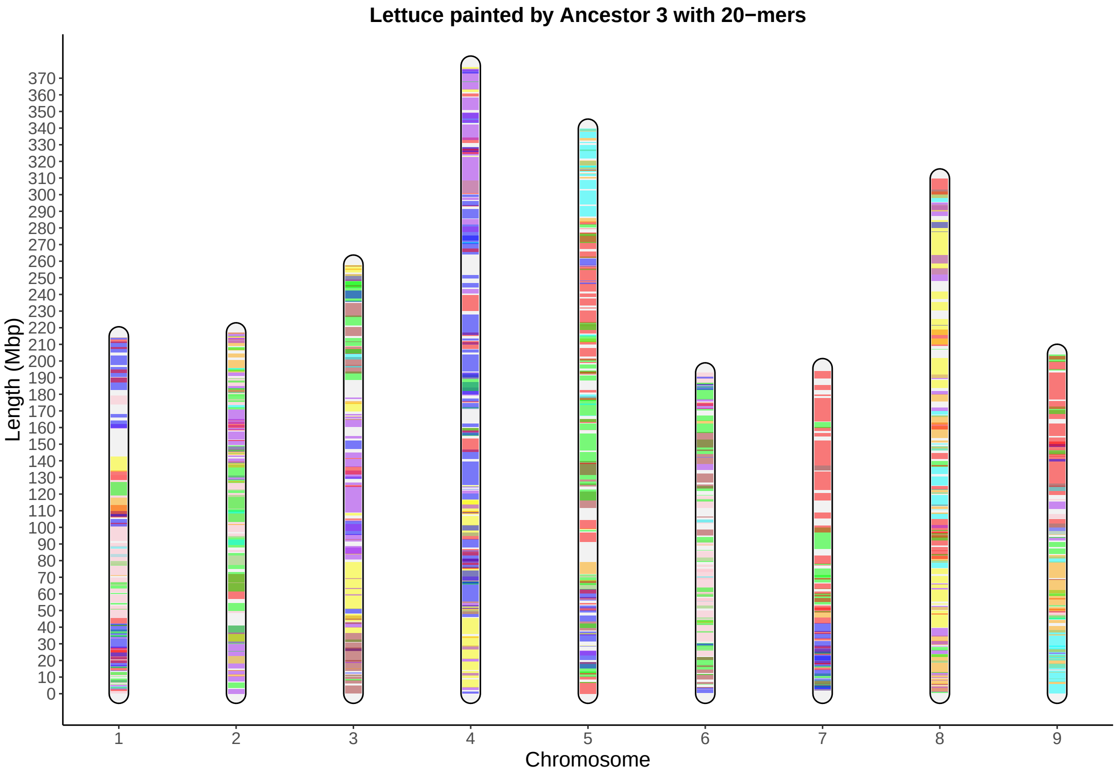

Lettuce painted by Ancestor 3 with 20−mers
Length (Mbp)
Chromosome

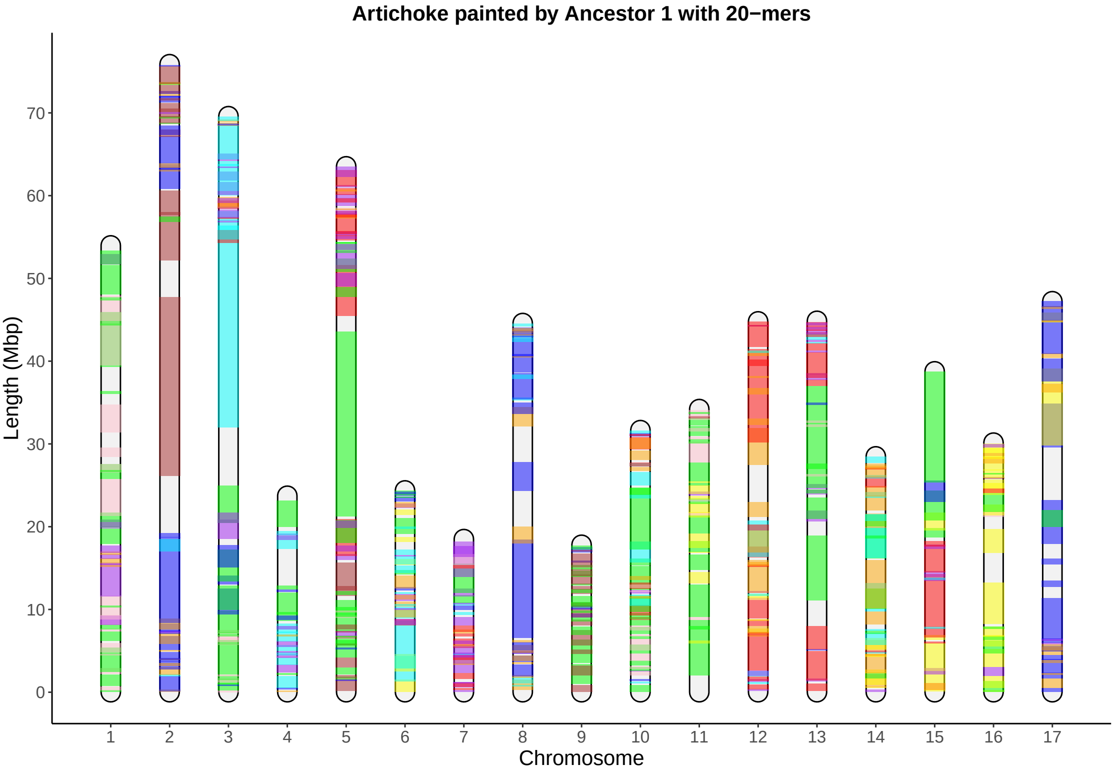

Artichoke painted by Ancestor 1 with 20−mers
Length (Mbp)
0
10
20
30
40
50
60
70
Chromosome

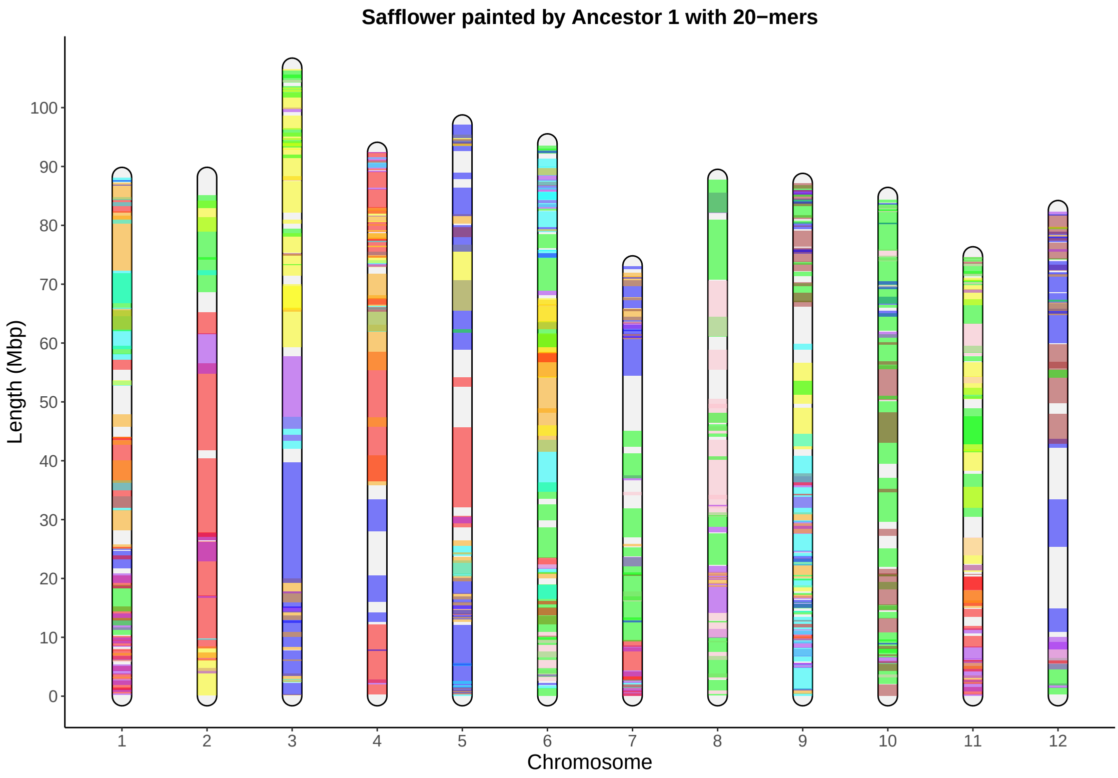

Safflower painted by Ancestor 1 with 20−mers
Length (Mbp)
0
10
20
30
40
50
60
70
80
90
100
Chromosome
1
2
3
4
5
6
7
8
9
10
11
12

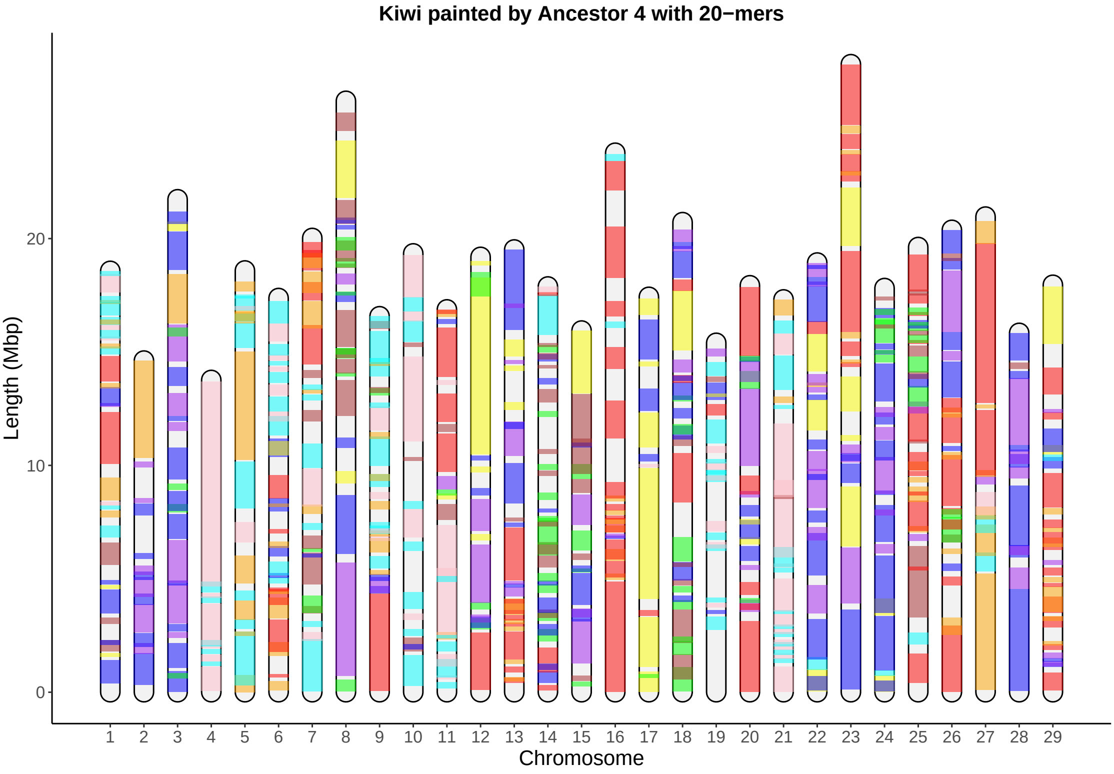
Kiwi painted by Ancestor 4 with 20−mers
Length (Mbp)
0
10
20
Chromosome

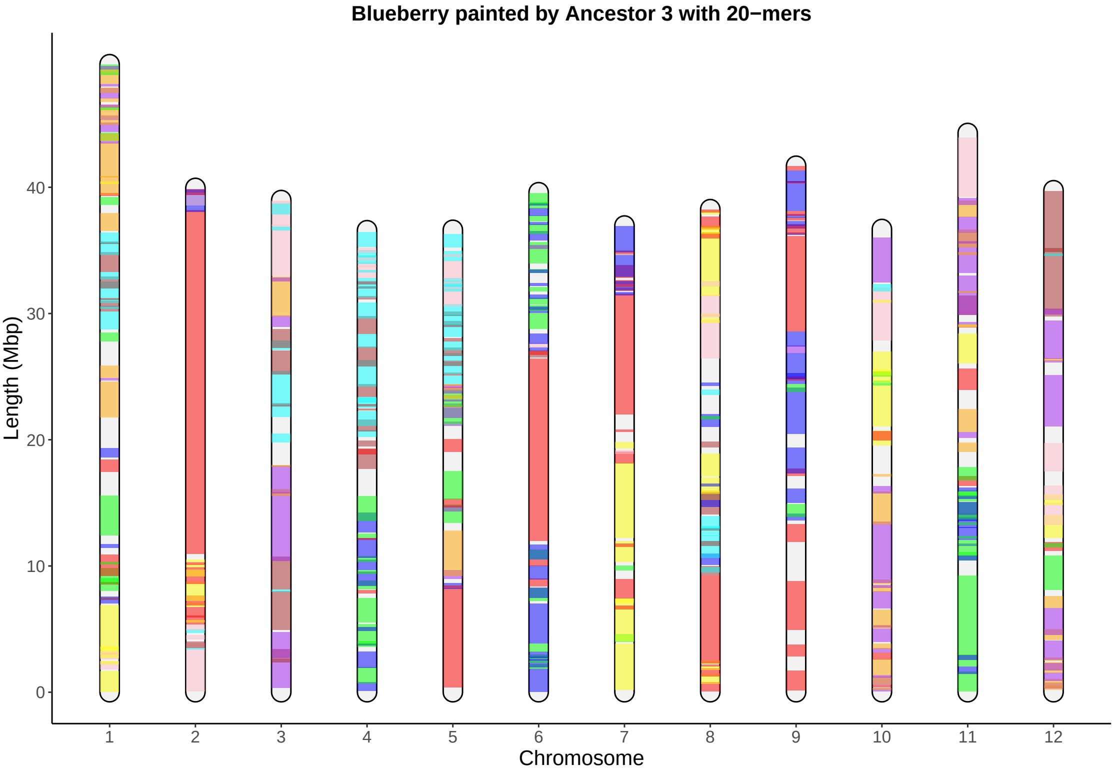

Blueberry painted by Ancestor 3 with 20−mers
Length (Mbp)
0
10
20
30
40
Chromosome
1
2
3
4
5
6
7
8
9
10
11
12

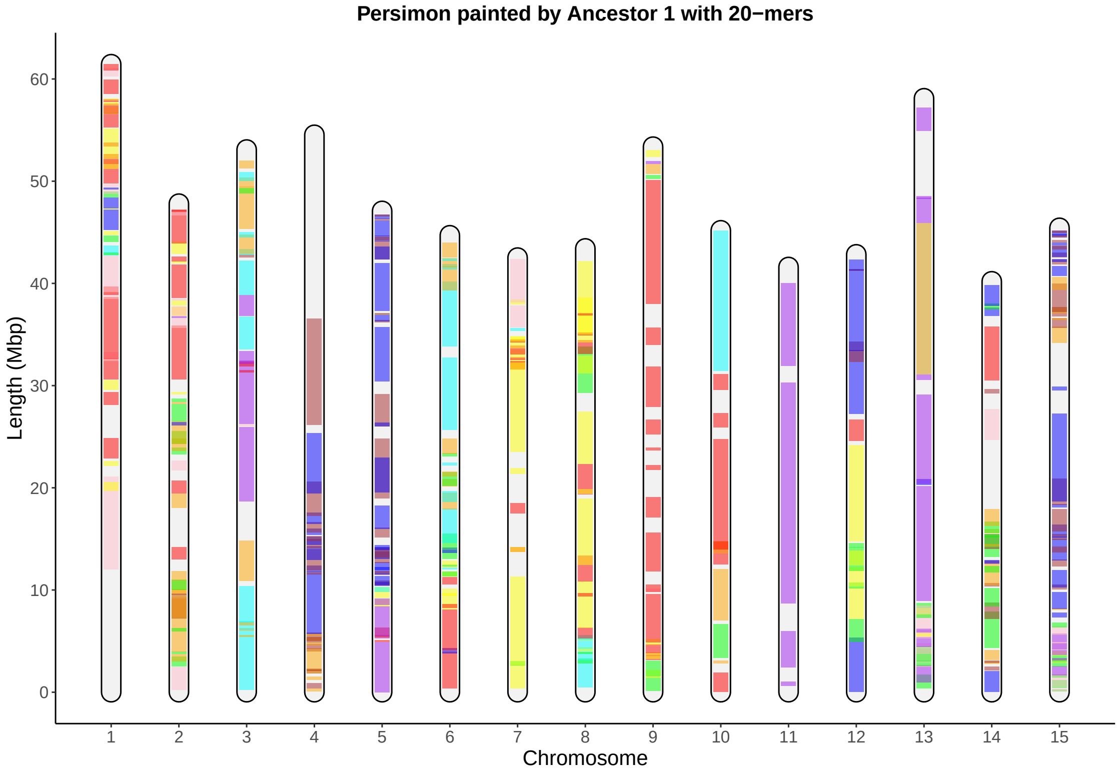

Persimon painted by Ancestor 1 with 20−mers
Length (Mbp)
0
10
20
30
40
50
60
Chromosome

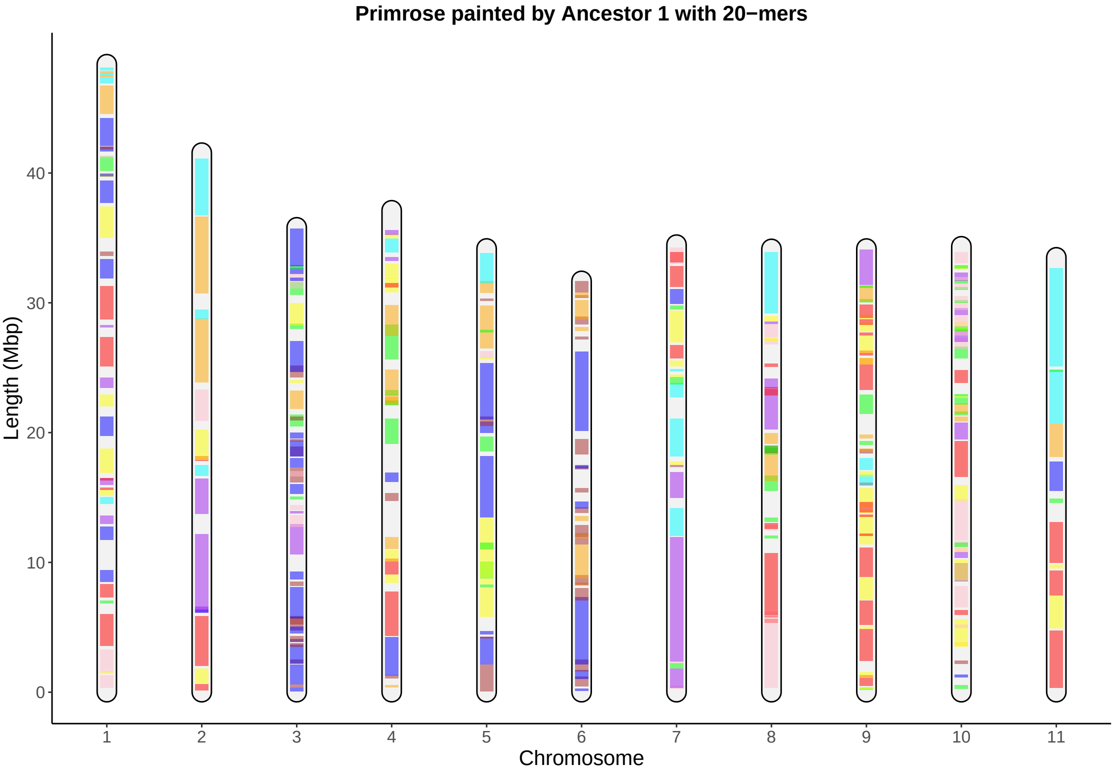

Primrose painted by Ancestor 1 with 20−mers
Length (Mbp)
0
10
20
30
40
Chromosome
1
2
3
4
5
6
7
8
9
10
11

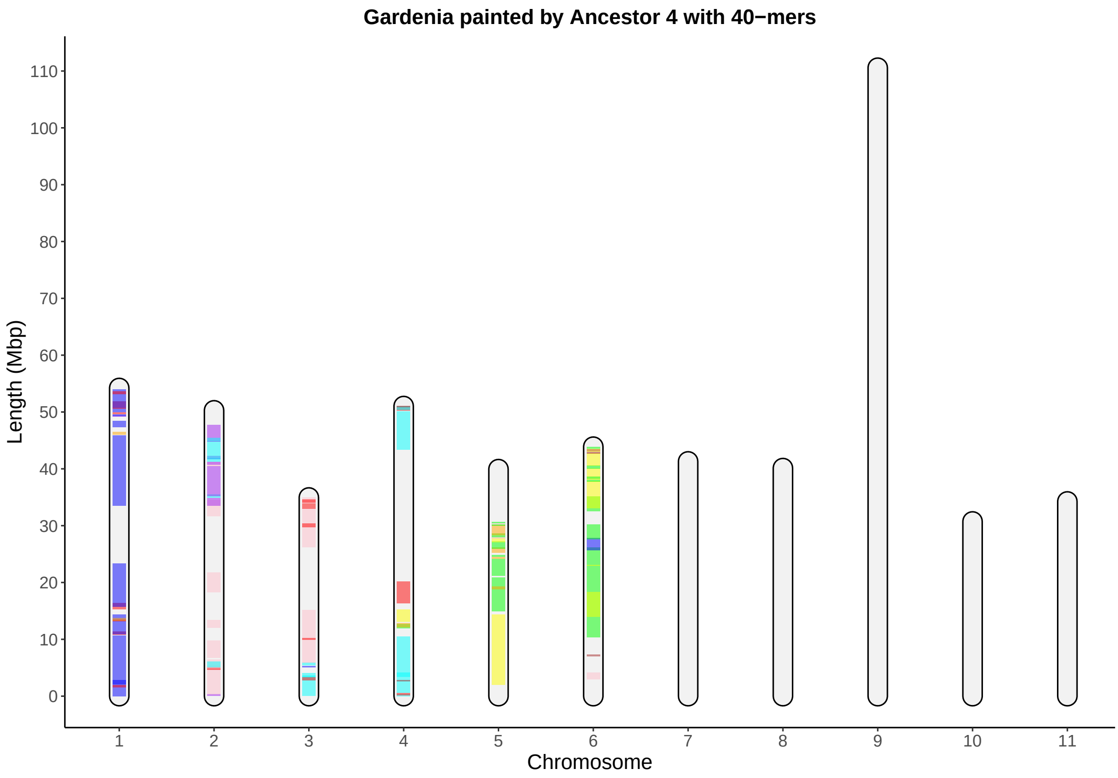

Gardenia painted by Ancestor 4 with 40−mers
Length (Mbp)
0
10
20
30
40
50
60
70
80
90
100
110
Chromosome
1
2
3
4
5
6
7
8
9
10
11

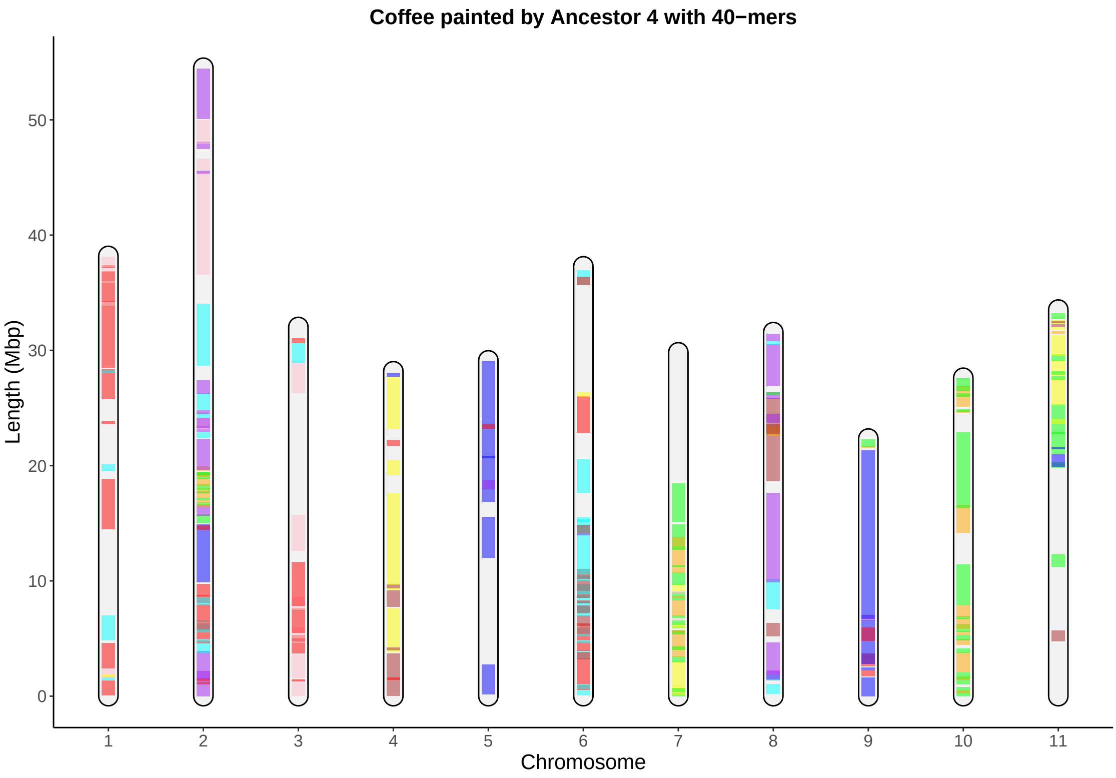

Coffee painted by Ancestor 4 with 40−mers
Length (Mbp)
0
10
20
30
40
50
1
2
3
4
5
6
7
8
9
10
11
Chromosome

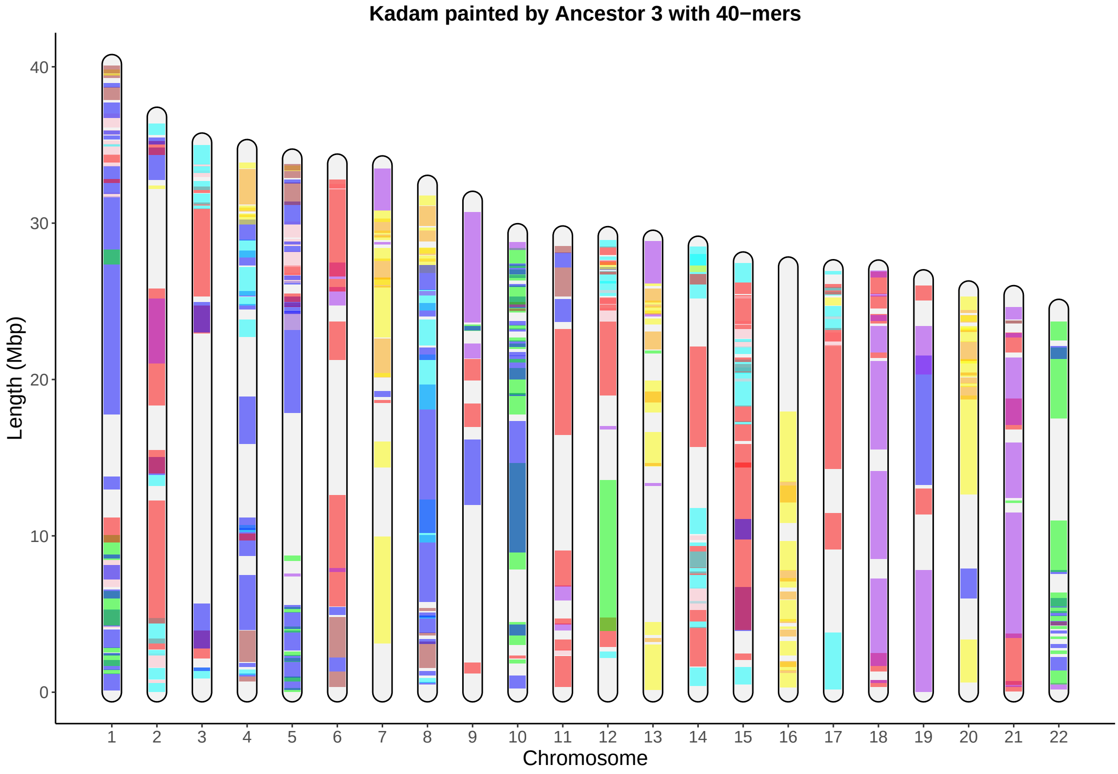

Kadam painted by Ancestor 3 with 40−mers
Length (Mbp)
0
10
20
30
40
Chromosome

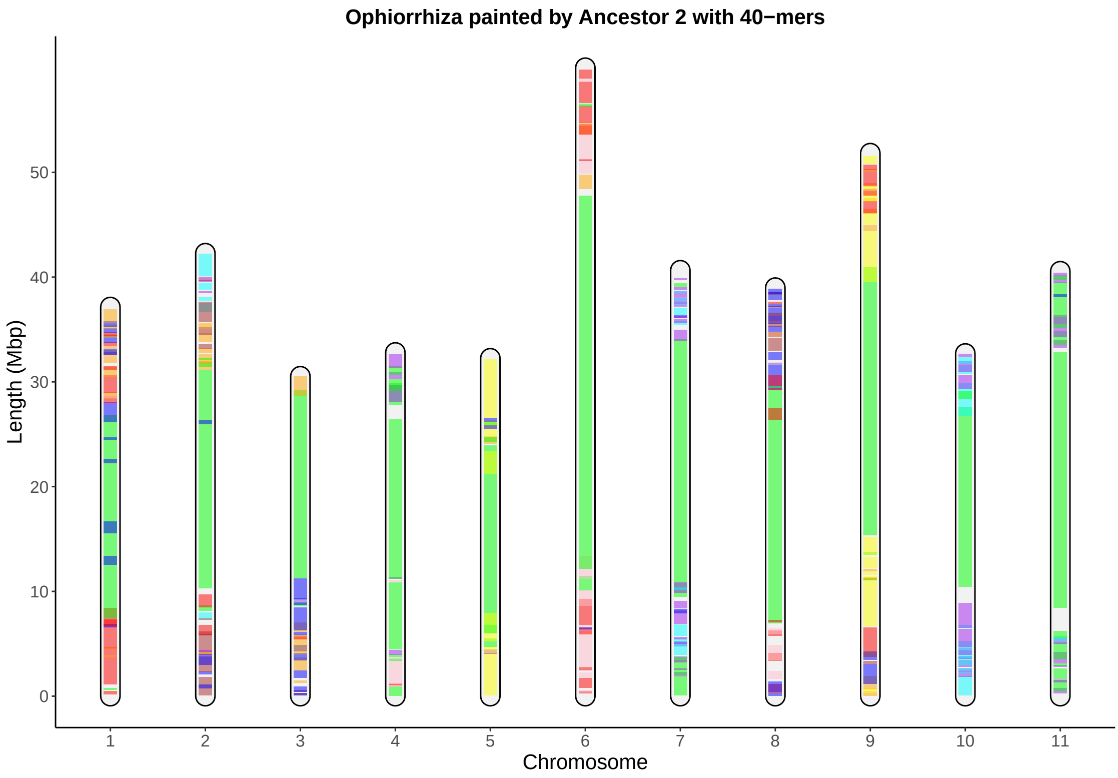

Ophiorrhiza painted by Ancestor 2 with 40−mers
Length (Mbp)
0
10
20
30
40
50
1
2
3
4
5
6
7
8
9
10
11
Chromosome

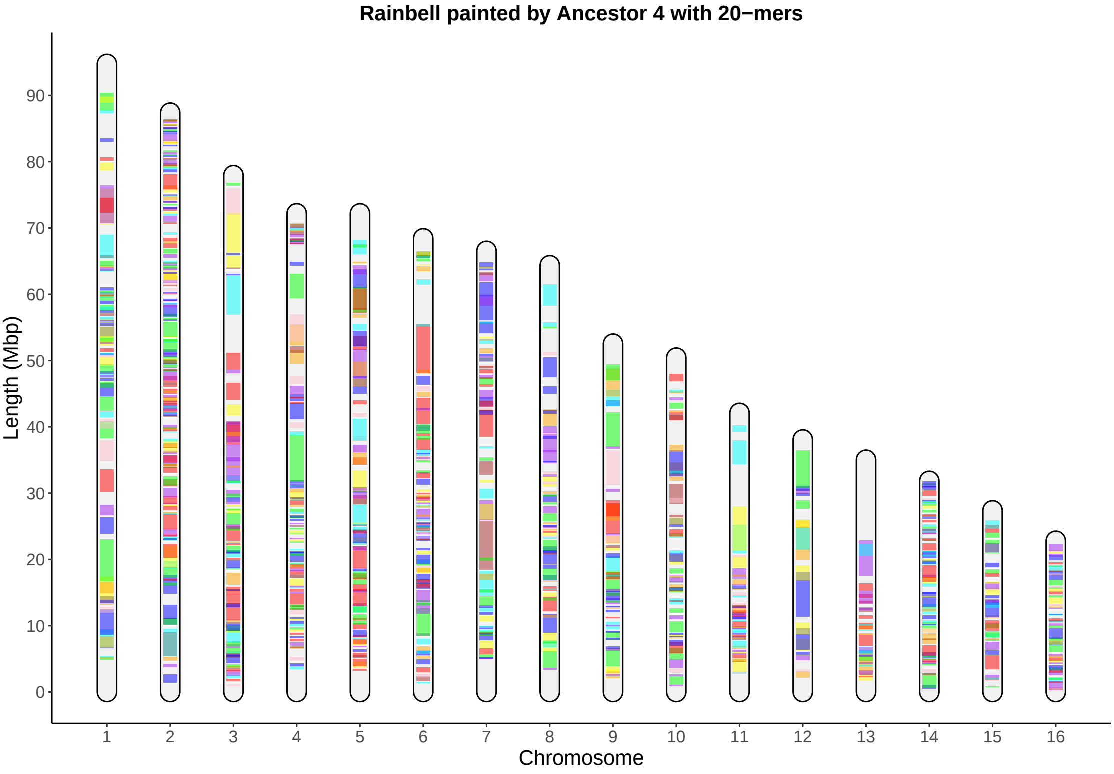

Rainbell painted by Ancestor 4 with 20−mers
Length (Mbp)
0
10
20
30
40
50
60
70
80
90
Chromosome

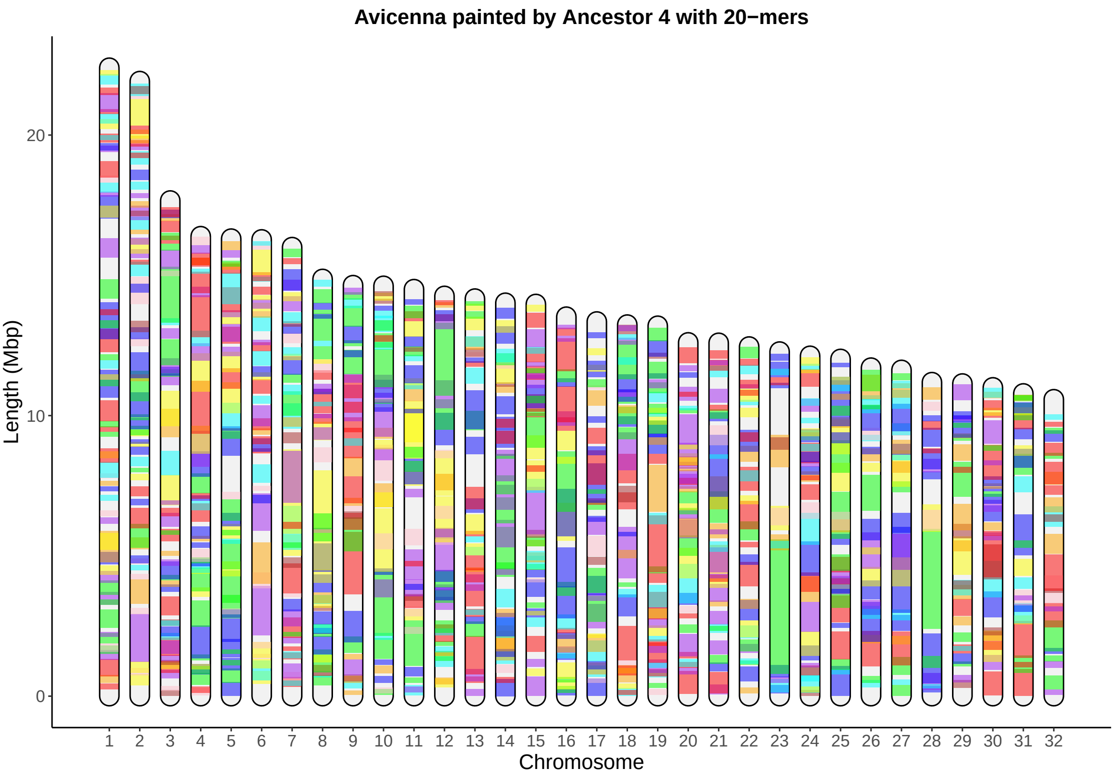
Avicenna painted by Ancestor 4 with 20−mers
Length (Mbp)
20
10
0
1 2 3 4 5 6 7 8 9 10 11 12 13 14 15 16 17 18 19 20 21 22 23 24 25 26 27 28 29 30 31 32
Chromosome

#### Teak painted by Ancestor 3 with 20−mers

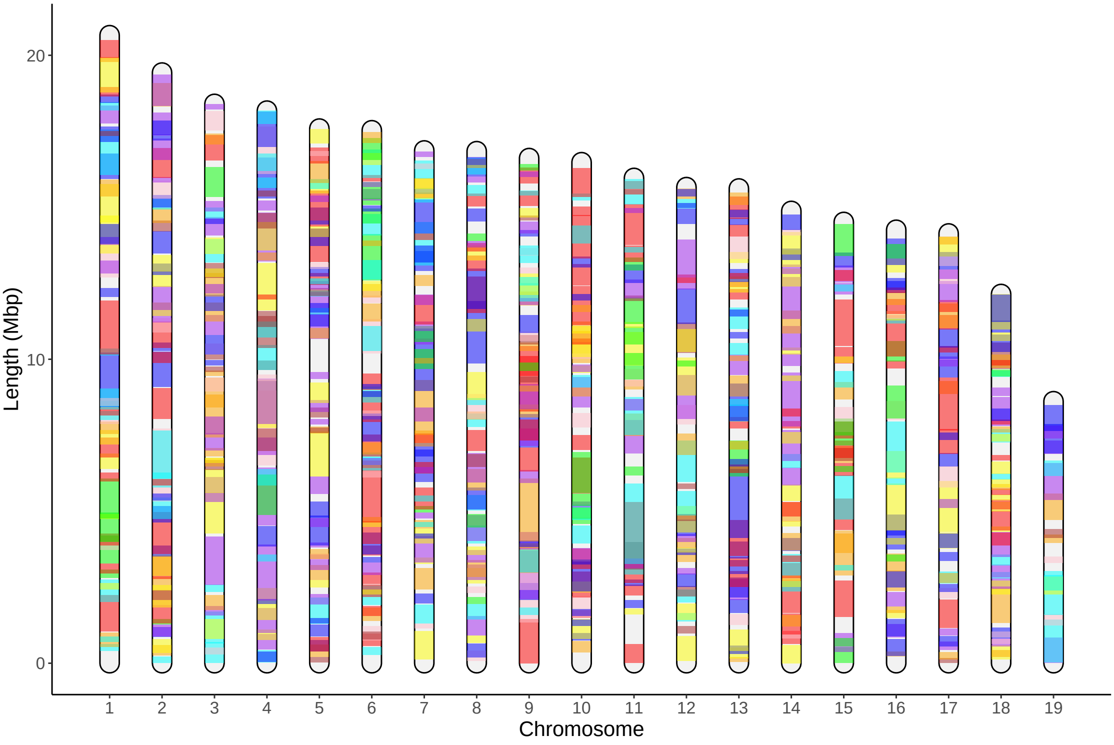

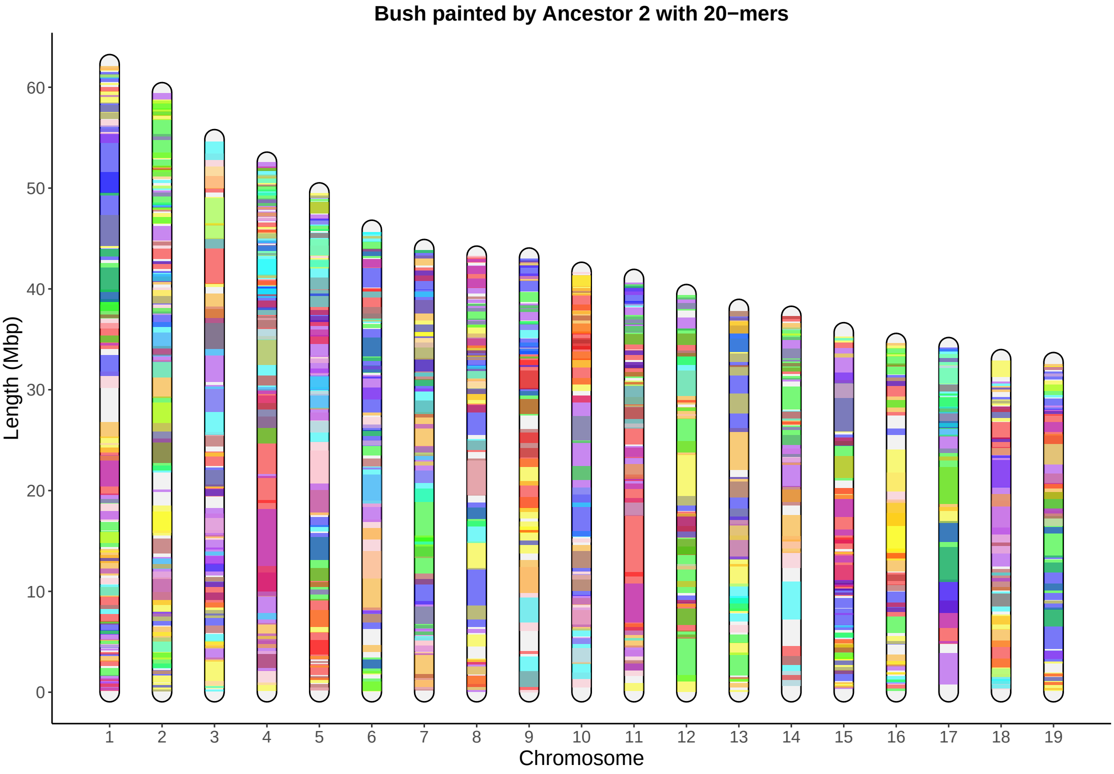

Bush painted by Ancestor 2 with 20−mers
Length (Mbp)
0
10
20
30
40
50
60
Chromosome

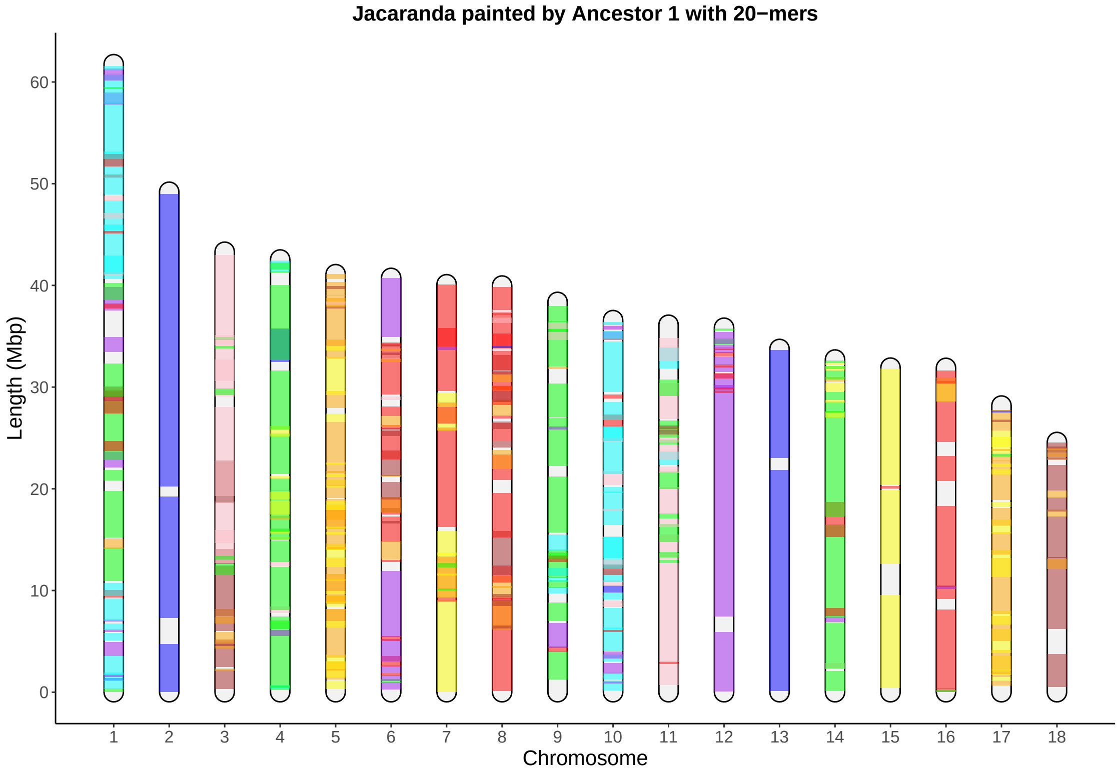

Jacaranda painted by Ancestor 1 with 20−mers
Length (Mbp)
0
10
20
30
40
50
60
Chromosome

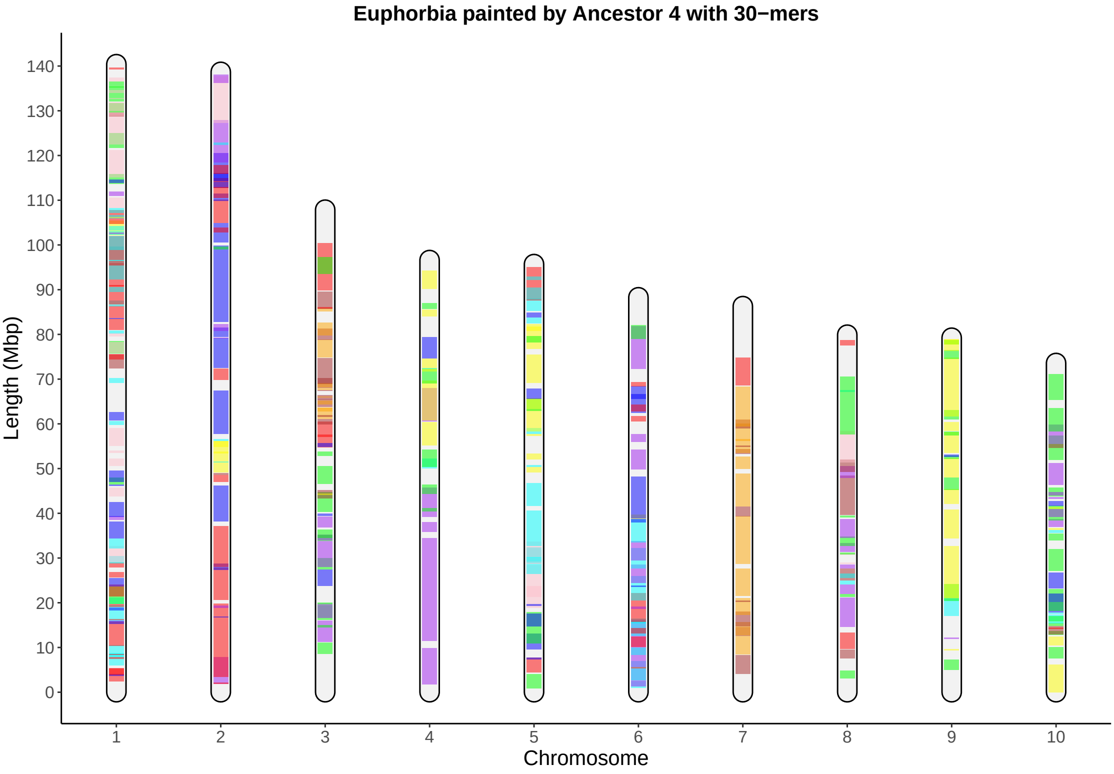

Euphorbia painted by Ancestor 4 with 30−mers
Length (Mbp)
0
10
20
30
40
50
60
70
80
90
100
110
120
130
140
Chromosome
1
2
3
4
5
6
7
8
9
10

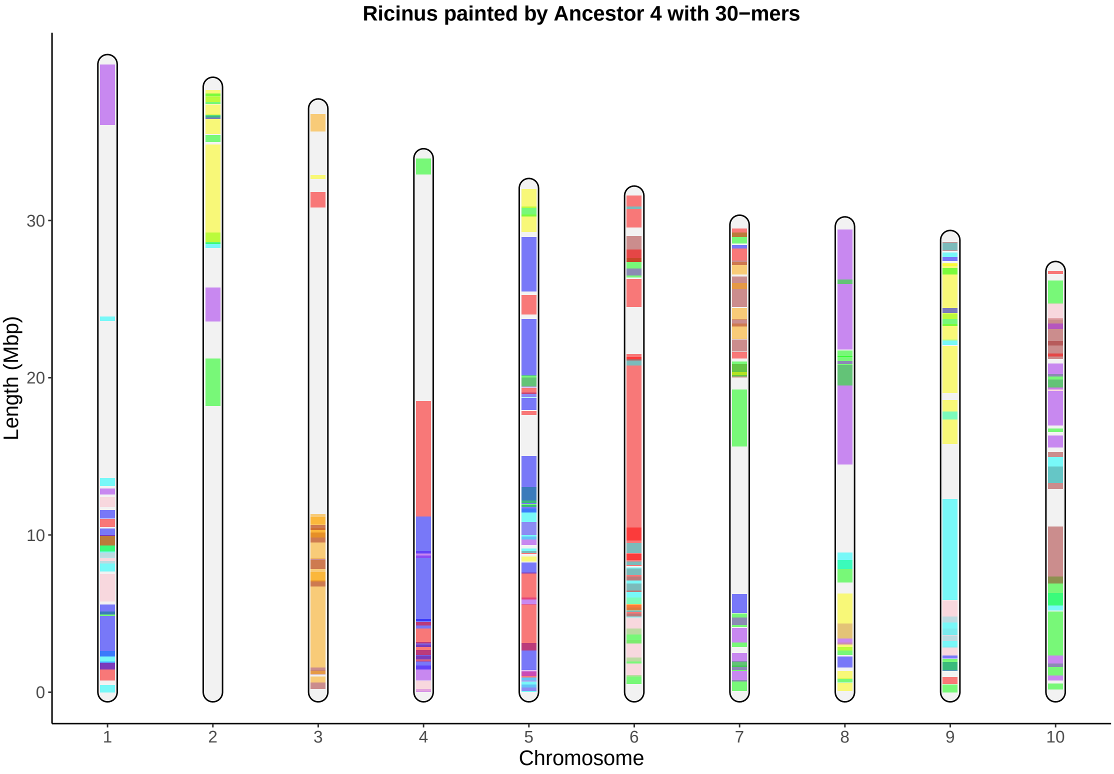

Ricinus painted by Ancestor 4 with 30−mers
Length (Mbp)
0
10
20
30
1
2
3
4
5
6
7
8
9
10
Chromosome

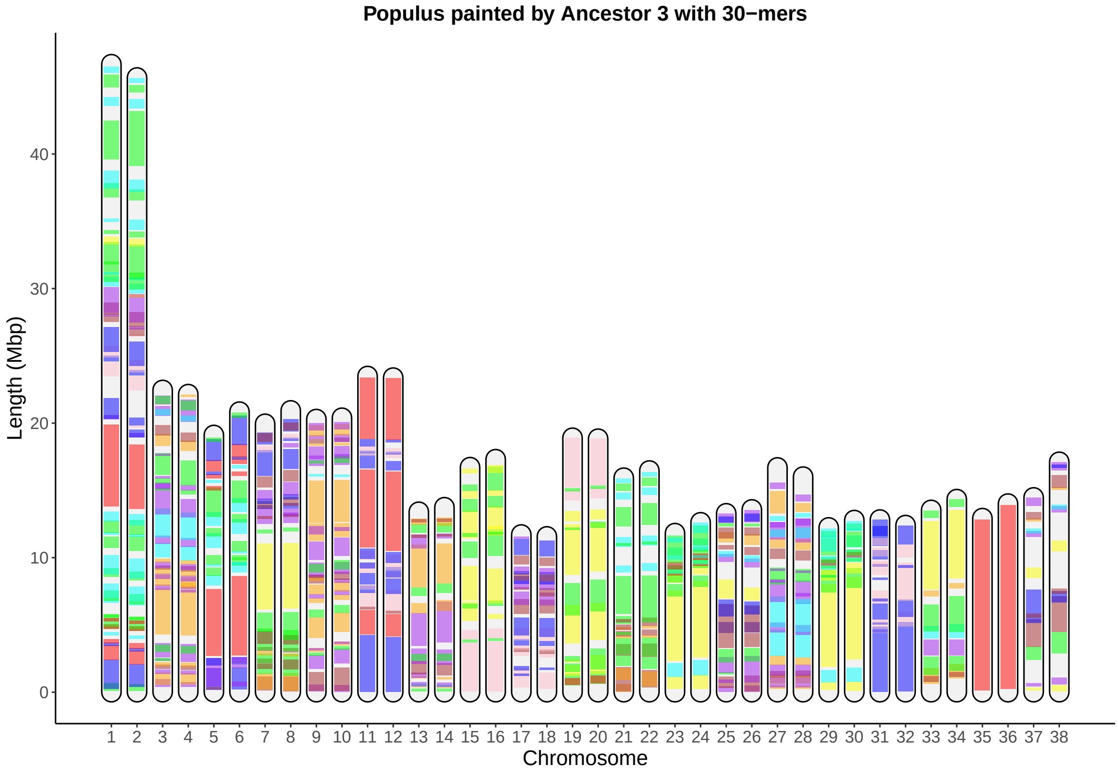
Populus painted by Ancestor 3 with 30−mers
Length (Mbp)
0
10
20
30
40
1 2 3 4 5 6 7 8 9 10 11 12 13 14 15 16 17 18 19 20 21 22 23 24 25 26 27 28 29 30 31 32 33 34 35 36 37 38
Chromosome

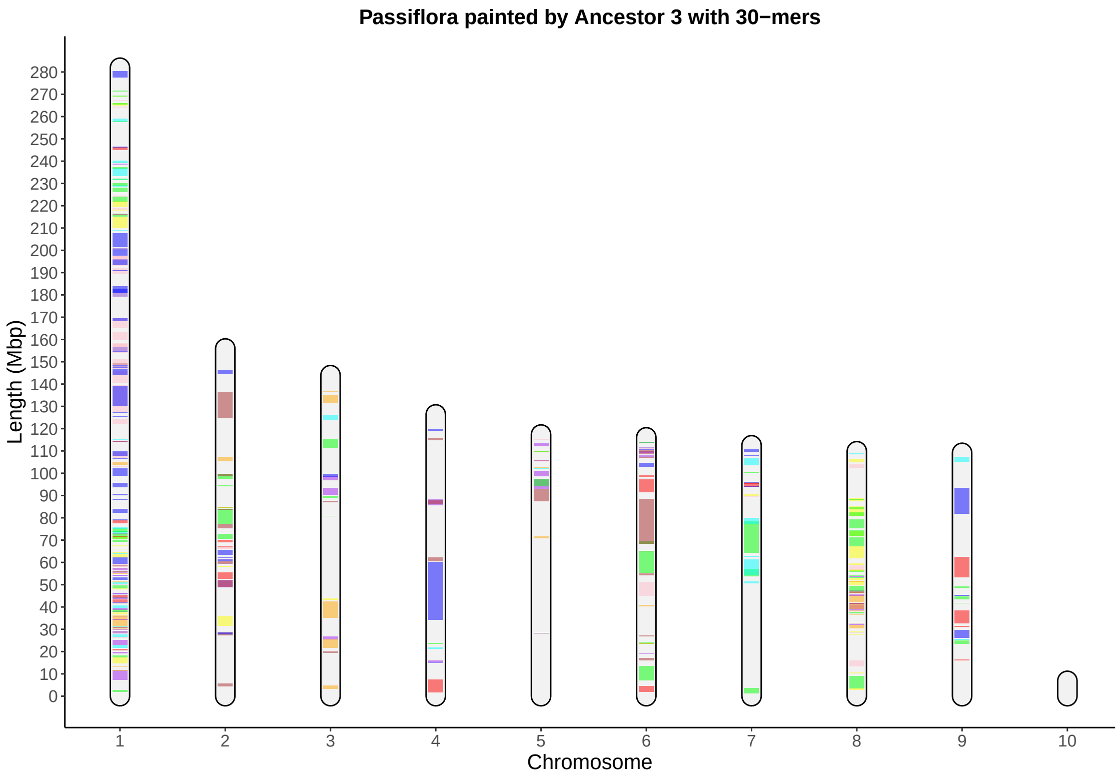

Passiflora painted by Ancestor 3 with 30−mers
Length (Mbp)
0
10
20
30
40
50
60
70
80
90
100
110
120
130
140
150
160
170
180
190
200
210
220
230
240
250
260
270
280
Chromosome
1
2
3
4
5
6
7
8
9
10

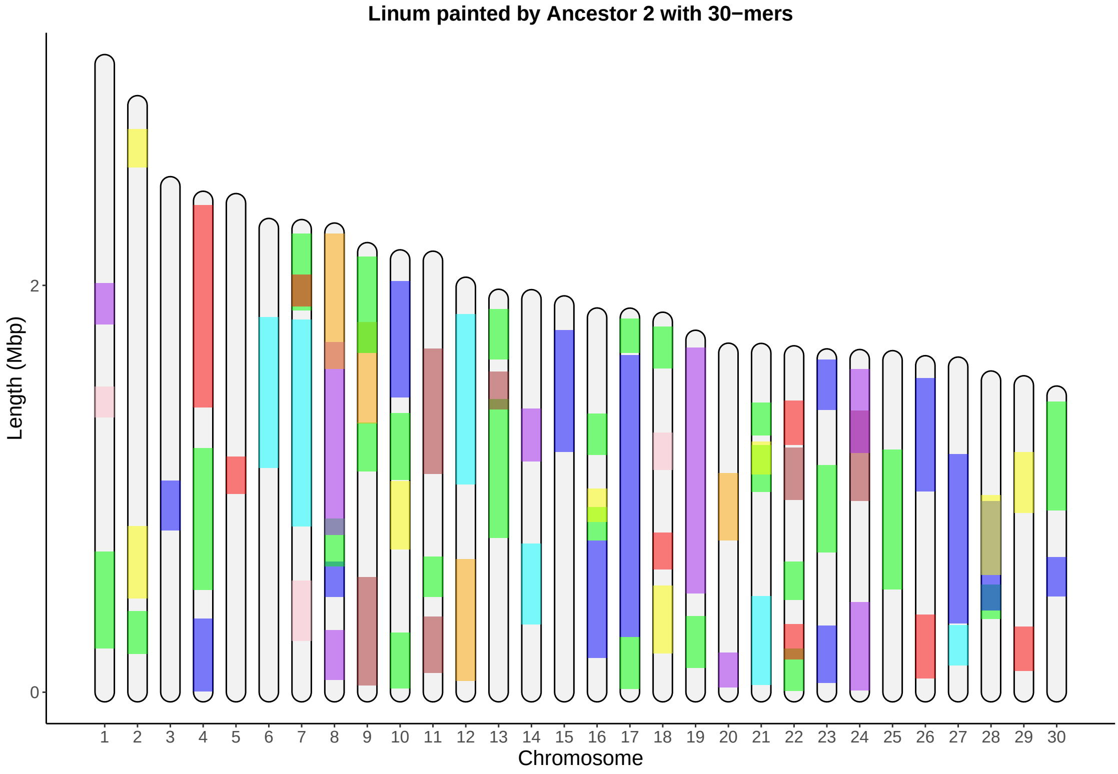

Linum painted by Ancestor 2 with 30−mers
Length (Mbp)
0
2
Chromosome

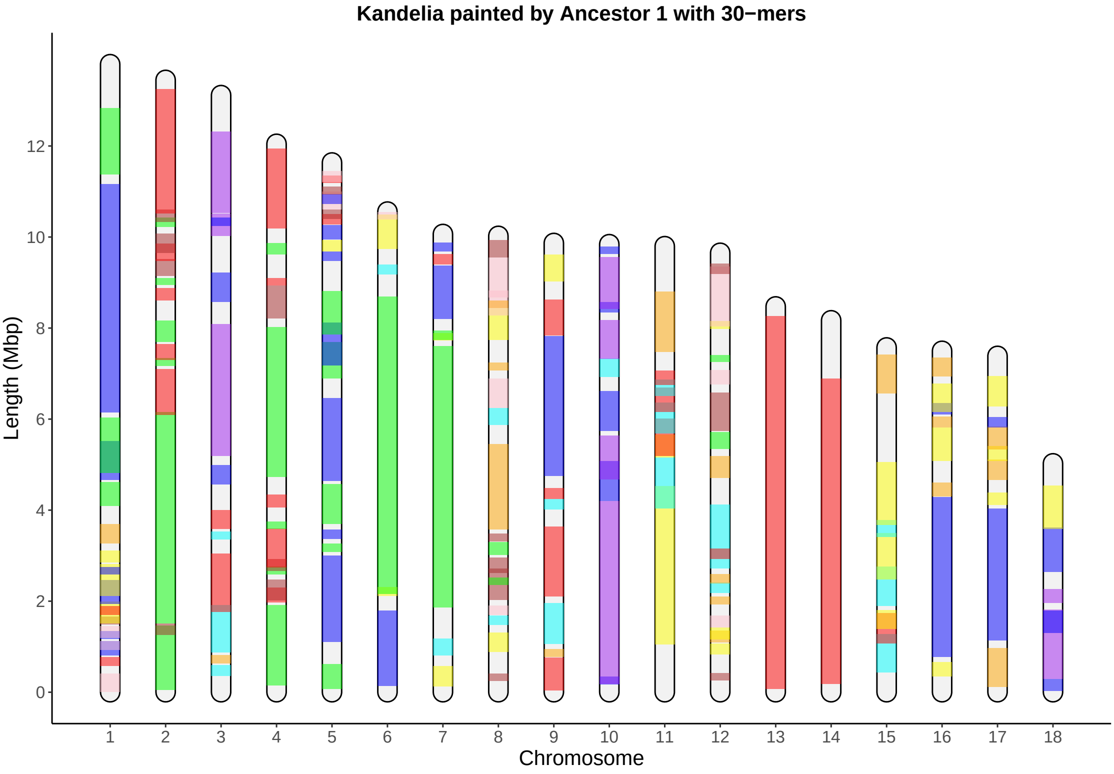
Kandelia painted by Ancestor 1 with 30−mers
Length (Mbp)
0
2
4
6
8
10
12
Chromosome

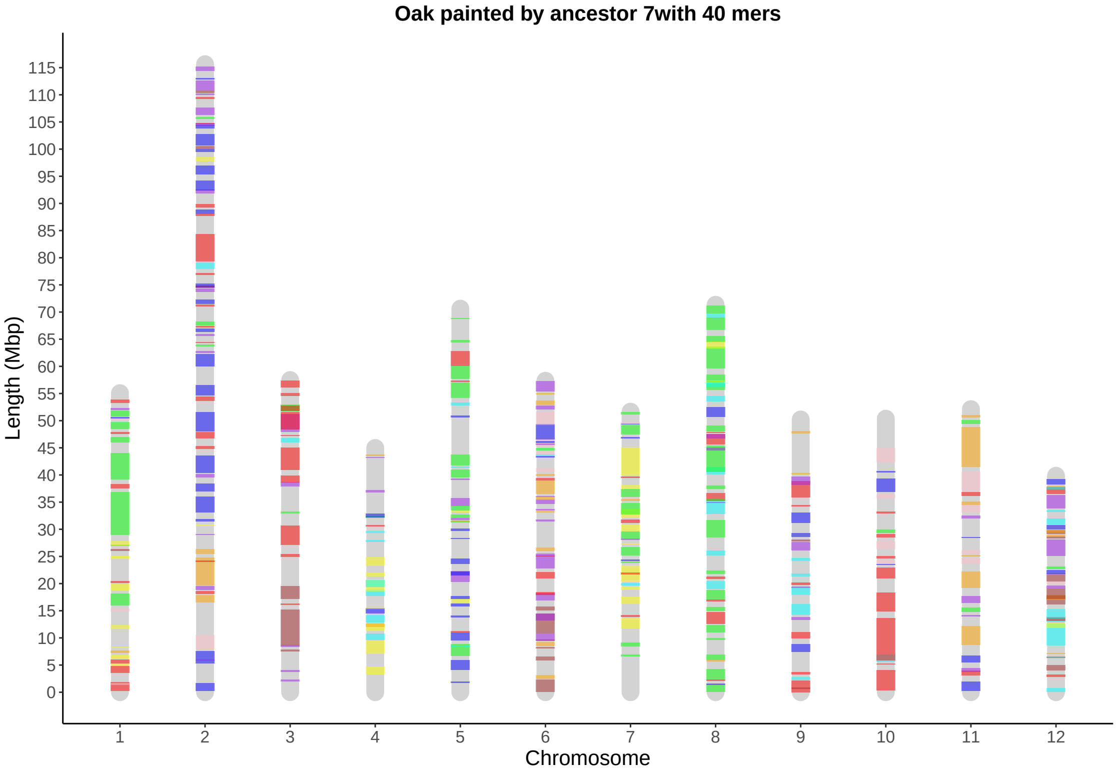

Oak painted by ancestor 7with 40 mers
Length (Mbp)
115
110
105
100
95
90
85
80
75
70
65
60
55
50
45
40
35
30
25
20
15
10
5
0
1
2
3
4
5
6
7
8
9
10
11
12
Chromosome

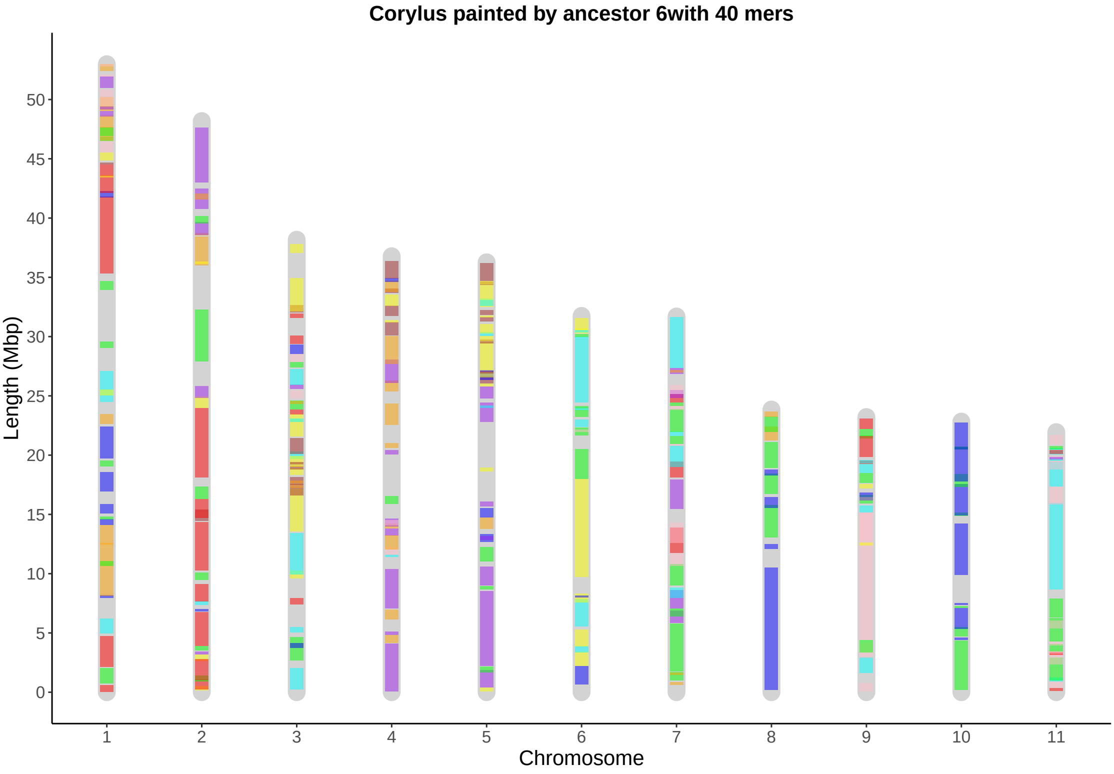

Corylus painted by ancestor 6with 40 mers
Length (Mbp)
0
5
10
15
20
25
30
35
40
45
50
Chromosome
1
2
3
4
5
6
7
8
9
10
11

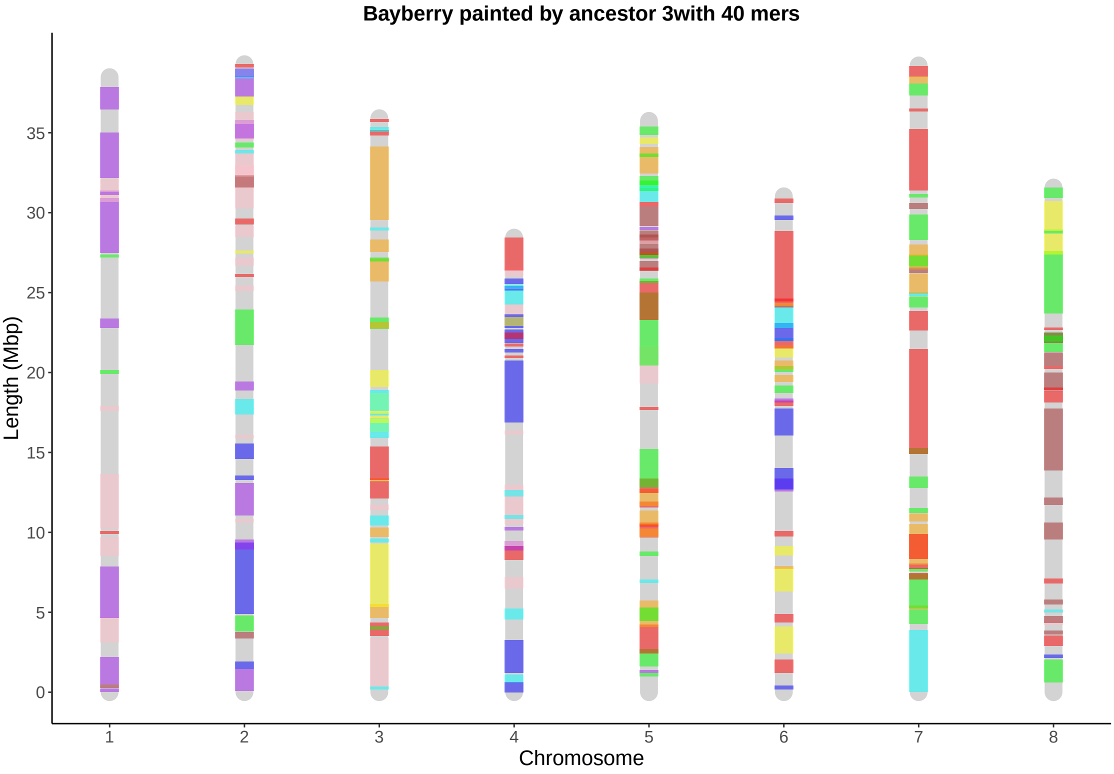

Bayberry painted by ancestor 3with 40 mers
Length (Mbp)
0
5
10
15
20
25
30
35
1
2
3
4
5
6
7
8
Chromosome

Springer Nature 2021 LaTeX template

### Supplement E. PCA clustering $g = 20$

Painting also available for 15-mers, 30-mers and 40-mers.

Fagales Ancestor 1

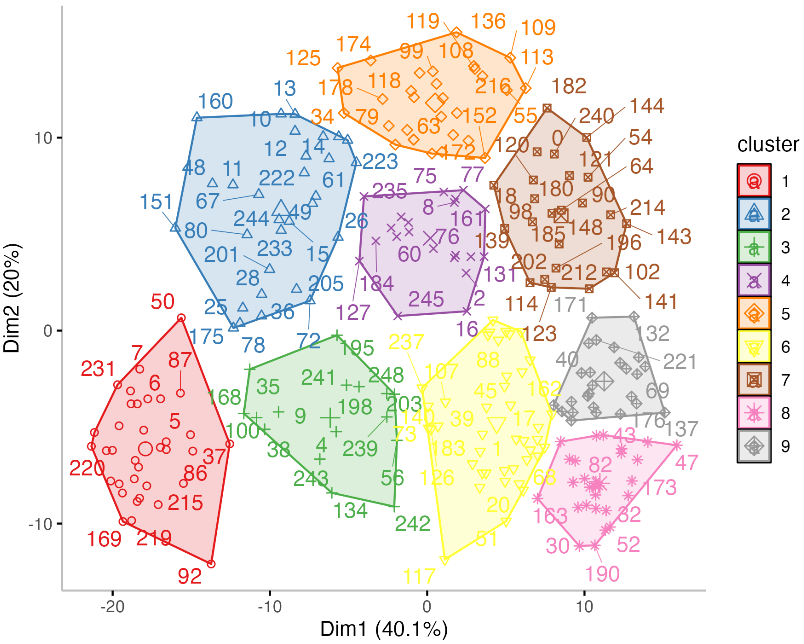

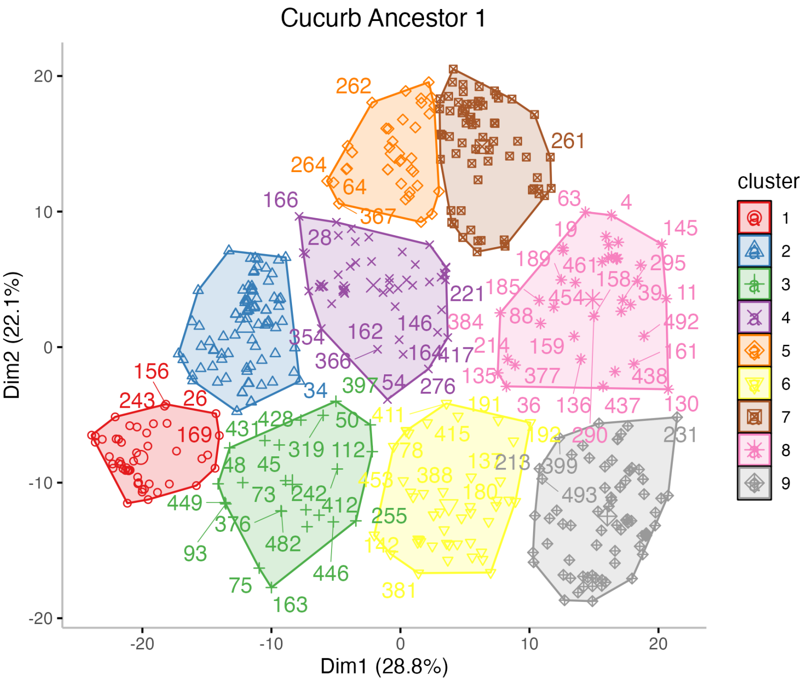

Cucurb Ancestor 1
Dim2 (22.1%)
Dim1 (28.8%)
20
10
0
-10
-20
-20
-10
0
10
20
cluster
1
2
3
4
5
6
7
8
9

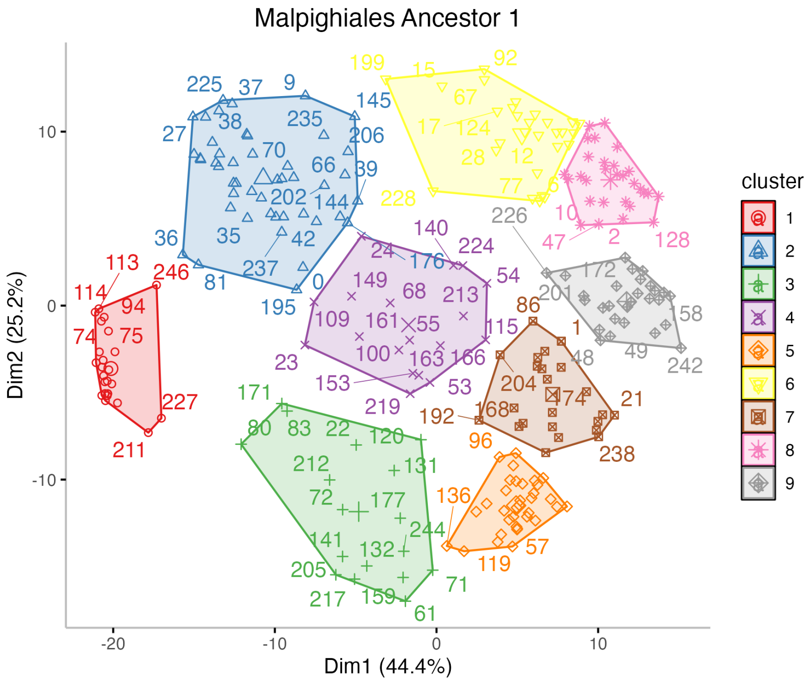

Malpighiales Ancestor 1
Dim1 (44.4%)
Dim2 (25.2%)
cluster
1
2
3
4
5
6
7
8
9

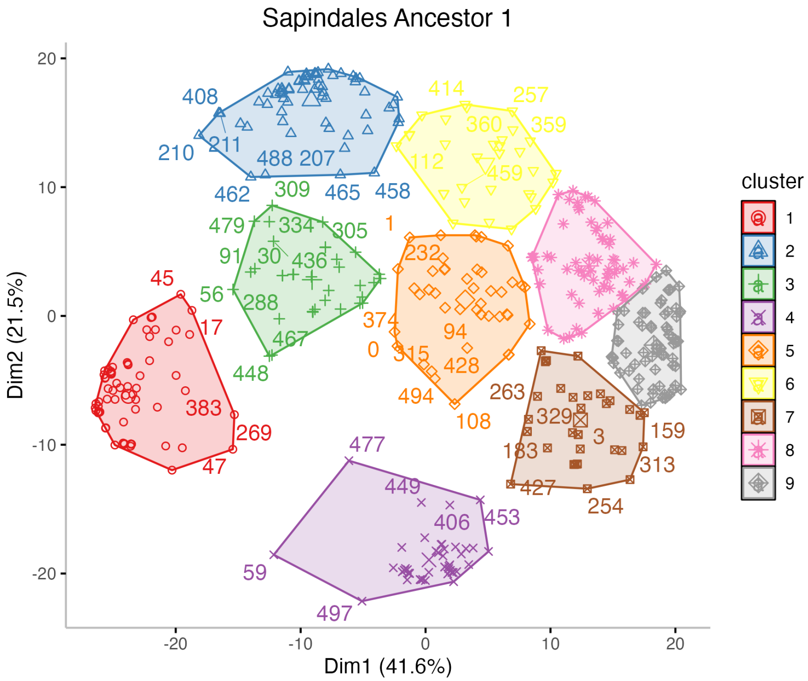

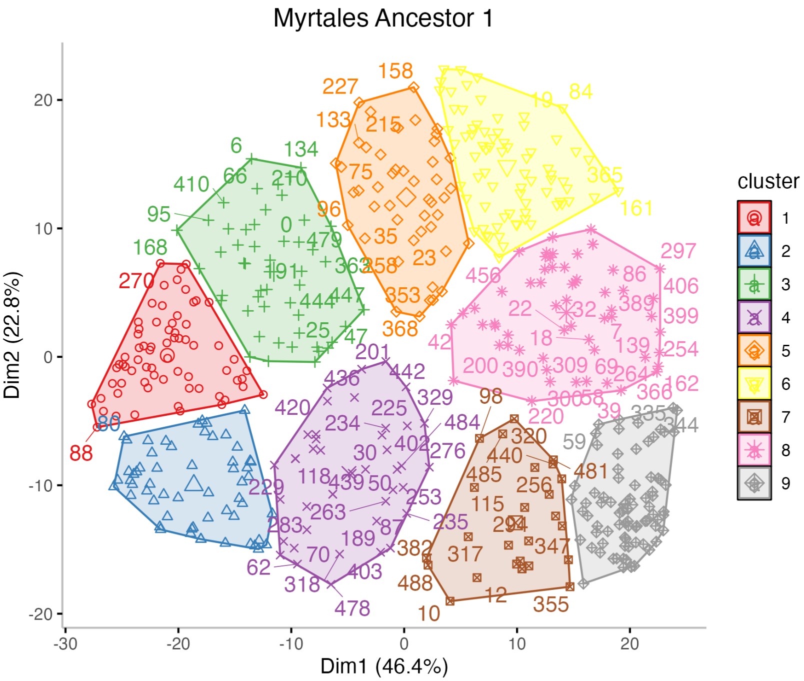
Myrtales Ancestor 1
Dim1 (46.4%)
Dim2 (22.8%)
-30
-20
-10
0
10
20
cluster
1
2
3
4
5
6
7
8
9

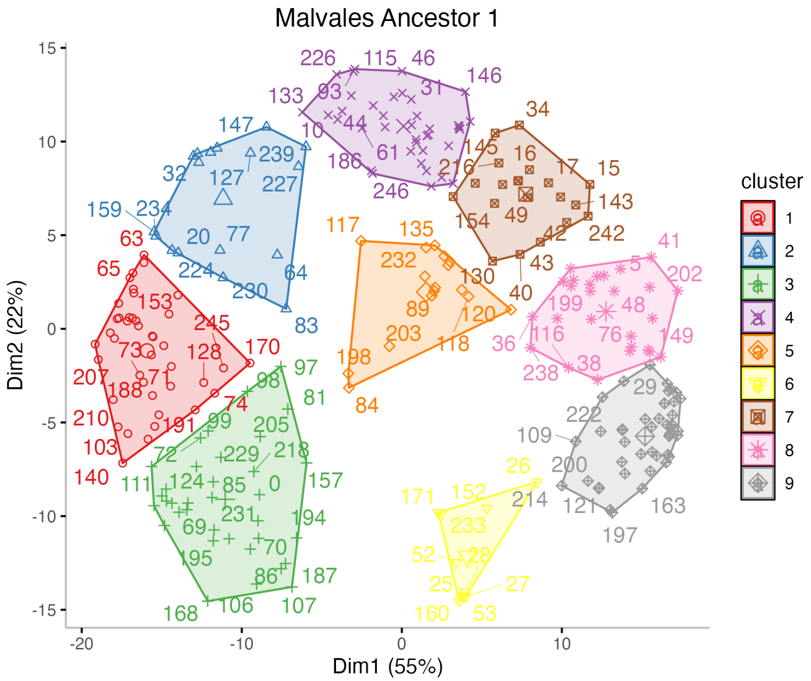
Malvales Ancestor 1
Dim1 (55%)
Dim2 (22%)
-20
-10
0
10
15
10
5
0
-5
-10
-15
cluster
1
2
3
4
5
6
7
8
9

Ericales Ancestor 1

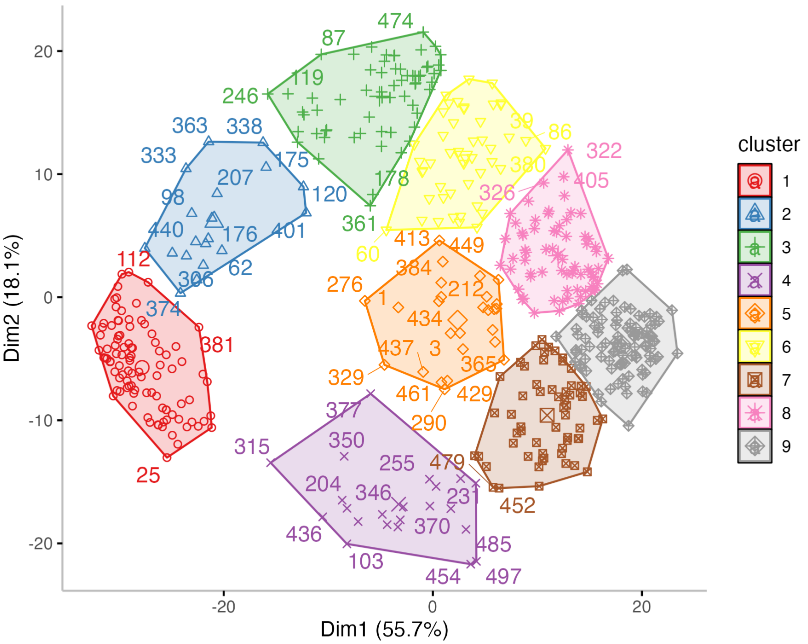

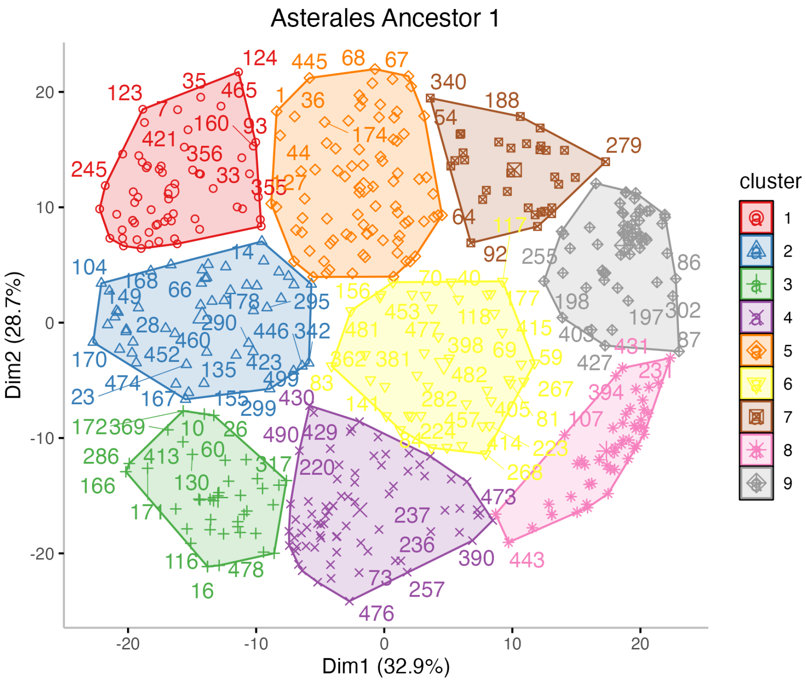
Asterales Ancestor 1
Dim2 (28.7%)
Dim1 (32.9%)
20
10
0
-10
-20
-20
-10
0
10
20
cluster
1
2
3
4
5
6
7
8
9

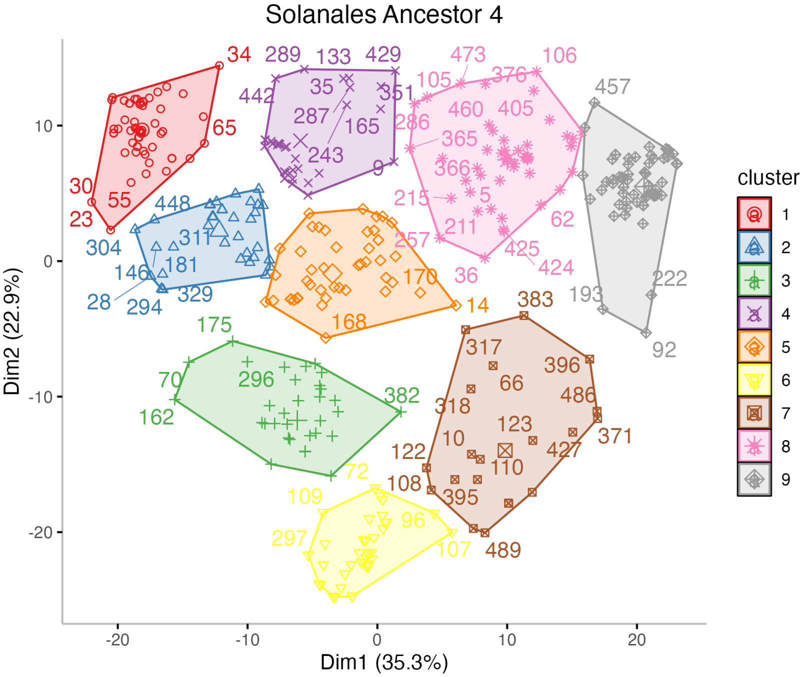

Solanales Ancestor 4
Dim1 (35.3%)
Dim2 (22.9%)
cluster
1
2
3
4
5
6
7
8
9

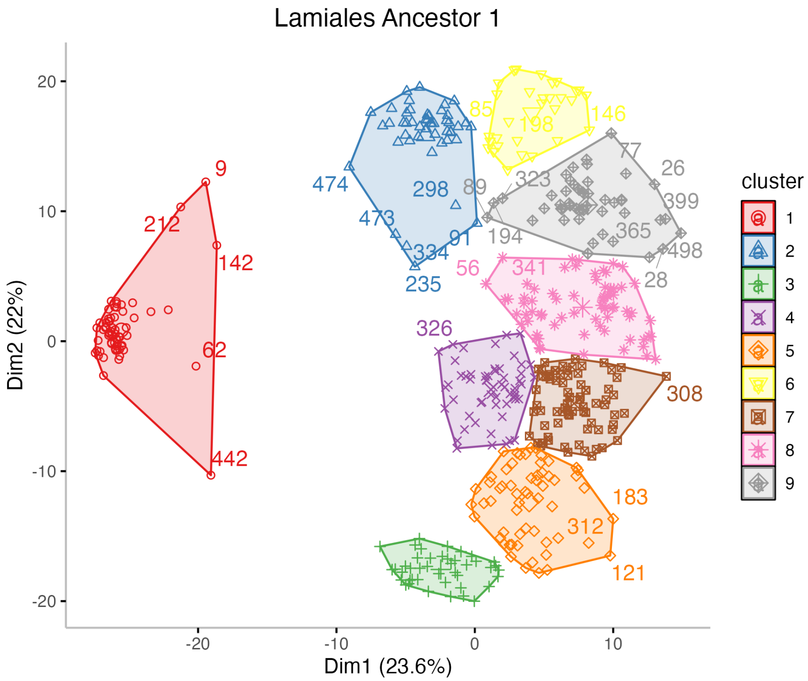

Lamiales Ancestor 1
Dim1 (23.6%)
Dim2 (22%)
cluster
1
2
3
4
5
6
7
8
9
20
10
0
-10
-20
-20
-10
0
10
9
212
142
62
442
474
298
473
334
91
235
85
198
146
89
323
77
26
399
194
365
498
28
56
341
326
308
183
312
121

Gentianales Ancestor 1

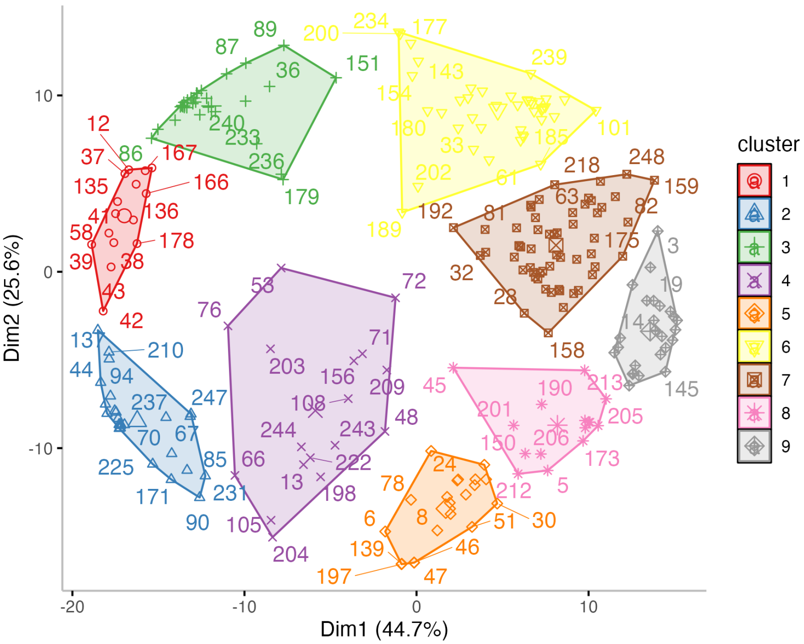
